# Universal modules for decoding amplitude and frequency of Ca^2+^ signals in plants

**DOI:** 10.64898/2025.12.13.694100

**Authors:** Fernando Vergara-Valladares, María E. Rubio-Meléndez, Myriam Charpentier, Erwan Michard, Ingo Dreyer

## Abstract

- Calcium signals are fundamental for plants and play a crucial role in long-term processes such as growth and development, as well as in rapid responses to environmental stimuli and stress factors. Nevertheless, the mechanisms involved in decoding calcium signal in plants are still largely unclear.
- Here, we have addressed the question of calcium signal decoding in a bottom-up modelling approach. We started with the thermodynamics of Ca^2+^ binding to a Ca^2+^ binding protein (CBP), e.g. via EF hands. Remarkably, Ca^2+^ binding properties of the EF hands do not coincide with the Ca^2+^ sensitivity of the protein containing these EF hands.
- In analysing the next levels of complexity, we identified six universal fundamental Ca^2+^-decoding modules, in which Ca^2+^ either interacts directly with a target protein (TP) or modulates its activity via a CBP. These modules are the basic units that enable the amplitude and frequency of Ca^2+^ signals to be decoded. Representatives of these modules are omnipresent in plant cells. They straightforwardly explain the puzzling finding that Ca^2+^-dependent kinases exhibit different Ca^2+^-sensitivities when tested with different substrates.
- In-depth analysis of the properties of the modules provides a fundamental theoretical basis for understanding Ca^2+^ signal decoding and may contribute to finding the “Rosetta Stone” for Ca^2+^ signals in plants.

## Introduction

Calcium (Ca^2+^) is a divalent cation that acts as a ubiquitous second messenger in cells of all phyla (Luan & Wang, 2021). One feature of Ca^2+^ signals is their ability to propagate within and between cells, with varying kinetics depending on the molecular processes involved (Allan *et al*., 2022; Matsui, 2022). In plants, calcium signals, whether transient or oscillatory, are triggered by a variety of stimuli, such as biotic, mechanical, chemical, or nutritional (Tian *et al*., 2020; HJ Lee & Seo, 2021), and play a crucial role in long-term processes such as growth and development, as well as in rapid responses to environmental cues and stress factors. Calcium gradients promote and control polarized cell growth (Hepler *et al*., 2001), and symplastic waves propagate in vascular tissues (Nguyen *et al*., 2018; Toyota *et al*., 2018).

Under physiological conditions, the cytosolic Ca^2+^ concentration ([Ca^2+^]_cyt_) in plant cells is typically in the range of 100 to 200 nM as measured with high-affinity reporters (Kader & Lindberg, 2010; Grenzi *et al*., 2021) and is significantly lower than that of K^+^, Na^+^, and Mg^2+^, but comparable to that of H^+^, which exhibits remarkable similarities in terms of buffer and signalling functions (Contador-Álvarez *et al*., 2025). Given that prolonged elevated levels of [Ca^2+^]_cyt_ can lead to precipitation with inorganic phosphate, resulting in cellular toxicity, cells tightly regulate calcium homeostasis (Carafoli & Krebs, 2016). This low [Ca^2+^]_cyt_ is maintained by the concerted action of membrane transporters, in particular primary active transporters (Ca^2+^-ATPases: ACA, ECA) and secondary active transporters (CAX), promoting the secretion of Ca^2+^ ions into the apoplast as well as their compartmentalization in organelles such as the vacuole, mitochondria, and the endoplasmic reticulum (Brownlee & Wheeler, 2025). In addition, buffering proteins and lipids could act as calcium sponges (Williams, 2002; Eisner *et al*., 2023) and may modulate calcium signalling and responses (Allan *et al*., 2022).

Upon various stimuli, Ca^2+^-permeable channels open in the plasma membrane and endomembrane, leading to a sharp increase in calcium levels in the cytosol, but sometimes also in the nucleus, the ER and mitochondria (Resentini *et al*., 2021; Cook *et al*., 2025). [Ca^2+^]_cyt_ can then transiently fluctuate between 100 nM and 5 μM (Malhó *et al*., 1998; Grenzi *et al*., 2021). These dynamic changes in Ca^2+^ concentration are perceived by a diverse array of Ca^2+^-sensing proteins, including calmodulins (CaMs), calmodulin-like proteins (CMLs), Ca^2+^-dependent protein kinases (CDPKs), Ca^2+^/calmodulin-dependent protein kinases (CCaMKs), and CBL-CIPK protein complexes (Kudla *et al*., 2018; Pirayesh *et al*., 2021), but also ion channels (Kintzer & Stroud, 2016) and enzymes like RBOH (Ogasawara *et al*., 2008; Kaya *et al*., 2014).

With a few exceptions, such as the C2 domain in animal PKC, Ca^2+^-sensitive proteins have a common structural motif for calcium binding: the EF-hand. This structure is remarkably well conserved in prokaryotes and eukaryotes and binds calcium in protein sensors (calmodulin), buffers (calbindin), and storage proteins (calsequestrin). The EF hand is a helix-loop-helix structure, characterized by a 12-residue sequence within the loop that coordinates a Ca^2+^ ion (Persechini *et al*., 1989). A binding event leads to a conformational change that causes the EF hand to close and the two alpha-helices to converge. Typically, Ca^2+^-sensing proteins possess four EF hands organized in two independent domains (DeFalco *et al*., 2010). Within each domain, the EF hands often exhibit cooperativity, whereby the binding of a Ca^2+^ ion to one EF hand facilitates the binding of a second Ca^2+^ ion to the other (Ikura, 1996). EF hands can be further classified as canonical and non-canonical. Canonical EF hands exhibit a high degree of conservation of negatively charged residues, enabling them to coordinate Ca^2+^ ions with dissociation constants ranging from hundreds of nanomolar to tens of micromolar. In contrast, non-canonical EF hands, lacking these negatively charged residues, often display reduced or absent calcium-binding capacity, suggesting that they may play primarily structural roles or exhibit higher affinity for other ions (Gifford *et al*., 2007).

There are two types of calcium-binding proteins: proteins that are directly regulated by calcium, and calcium signalling proteins that bind to downstream proteins and regulate them, ultimately orchestrating the plant’s physiological response to the stress it is experiencing. Based on their mode of action, this latter group of proteins can be broadly categorized into those that modulate target proteins by direct binding and those that modulate them by phosphorylation.

The first category of directly binding proteins encompasses calmodulins (CaMs) and calmodulin-like proteins (CMLs) (Ranty *et al*., 2006; Perochon *et al*., 2011). Upon Ca^2+^ binding, these proteins undergo a conformational change, particularly in the linker region connecting the two homologous domains, the N- and C-lobes, which each contain two Ca^2+^-binding EF-hands. The conformational change generates an amphipathic alpha-helix that can interact with specific binding sites on target proteins (Ishida & Vogel, 2006). These binding sites include IQ domains, as observed in ion channels and transporters such as NHXs (Daniel-Mozo *et al*., 2024), and calmodulin-binding domains, as found in the Ca^2+^/calmodulin-dependent protein kinase CCaMK (Swainsbury *et al*., 2012). Protein activation is generally associated with Ca^2+^ binding, but there is also a way known as modulation by apocalmodulin (apoCaM), which has been reported to open Ca_v_ and Na_v_ ion channels of animals (Adams *et al*., 2014), and to activate enzymes such as Phosphorylase b kinase, Adenylyl cyclase and iNO synthase even without Ca^2+^ (Jurado *et al*., 1999). At present, modulation by apoCaM remains poorly documented in plants. Nevertheless, recent research has identified some Cyclic-Nucleotide-Gated-Channels that are modulated by apoCaM in plants. In *Arabidopsis thaliana* pollen tubes, the channel complex CNGC18/8 is activated by apoCaM2 binding and becomes inactive with increased cytosolic calcium (Pan *et al*., 2019). Similarly, AtCNGC12 shows both positive and negative activity changes depending on the binding of apoCaM and CaM at multiple channel binding sites (DeFalco *et al*., 2016).

Calcineurin B-like (CBL) proteins also possess the ability to bind calcium ions. However, due to the presence of fewer canonical EF hands, their calcium-binding capacity is generally considered to be lower (Sánchez-Barrena *et al*., 2005). CBL proteins form functional complexes with Calcineurin B-like interacting protein kinases (CIPKs), enabling the phosphorylation of various intracellular targets (Weinl & Kudla, 2009; Tang *et al*., 2020), leading to the second category of proteins that modulate through phosphorylation. This class includes Ca^2+^-dependent protein kinases (CDPKs), Ca^2+^/calmodulin-dependent protein kinases (CCaMKs), and CIPKs (Tang *et al*., 2020). These kinases modulate the activity of target proteins by phosphorylating specific amino acid residues, typically serine, threonine, or tyrosine. Notably, several CDPKs and CBL-CIPK proteins exhibit autophosphorylation activity, further influencing their kinase activity (Oh *et al*., 2012; Tang *et al*., 2020) or forming potentiating kinase modules (Köster *et al*., 2025).

While CaMs and CMLs exert their influence through direct protein-protein interactions, kinases do not necessarily require sustained binding to their target proteins. Thus, a single kinase can phosphorylate multiple proteins. The reverse reaction (dephosphorylation) requires the action of protein phosphatases. Some studies suggest a balance between kinase and phosphatase activities in response to stresses such as salinity (Senadheera & Maathuis, 2009; Praat *et al*., 2021). This balance represents an important point of intersection between Ca^2+^ signalling and other cellular signalling pathways, making it challenging to definitively attribute specific effects to kinase activity alone.

Significant progress has been made in elucidating the interactions between proteins involved in Ca^2+^ signalling pathways. *In vitro* studies have employed fluorescence-based techniques to monitor CaM activity (Mukherjee *et al*., 2022) and to investigate the catalytic activity and conformational changes of CDPK (Liese *et al*., 2024). Techniques such as isothermal titration calorimetry (ITC; Quinn *et al*., 2016) and stopped-flow methods have been instrumental in determining kinetic parameters for Ca^2+^ binding to proteins and protein-peptide interactions (Northrop & Simpson, 1997; Mazzei *et al*., 2016; Tso *et al*., 2022). These experimental approaches have provided valuable insights into the molecular mechanisms underlying cellular responses to various stresses. However, several challenges remain in the study of Ca^2+^ signalling. *In vitro* studies often encounter difficulties in maintaining physiologically relevant Ca^2+^ concentrations while observing measurable protein activity. Likewise, studies involving overexpression of proteins in cellular models may require the use of unnaturally high Ca^2+^ concentrations or prolonged exposure times, which can be toxic and limit the reproducibility of results. Furthermore, *in vivo* studies face the challenge of distinguishing the specific contributions of individual proteins within a complex network of interacting signalling pathways. Since cells utilize multiple signalling pathways to respond to stress, experimental approaches must carefully consider the involvement of other signalling molecules. In addition, gene silencing techniques can provide valuable insights by attenuating the activity of specific proteins and observing the resulting impact on cellular responses.

To supplement the diverse experimental work, a couple of theoretical approaches were carried out (Martins *et al*., 2013; Allan *et al*., 2022), which show that modelling can indeed provide deeper insights into that complex topic. Here, we have pursued this avenue further and addressed the question of whether the principles of biochemistry and thermodynamics that govern the interactions between proteins and Ca^2+^ ions can contribute to a deeper understanding of the decoding of Ca^2+^ signals in plants. In a rigorous bottom-up approach, we started with the thermodynamics of Ca^2+^ binding to a “Ca^2+^ binding protein” (CBP) and built on this basis the next layer of complexity represented by four fundamental Ca^2+^-decoding modules, in which a CBP modulates the activity of a target protein (TP). All of the physiological examples presented above could be assigned to one of these four modules. The in-depth analysis of the kinetics of the modules presented here provides a fundamental theoretical basis for understanding Ca^2+^ signal decoding in plants. The models may contribute to the targeted planning of future experimental investigations and serve as a basis for further in-depth follow-up studies.

## Materials and Methods

### Ca^2+^-binding to a protein

Compared to other cellular processes, the direct interaction of Ca^2+^ ions with a target protein is so fast that it can be considered almost instantaneous (HY Park *et al*., 2008; Swainsbury *et al*., 2012) and always in steady state. In this condition, the binding of the ligand Ca^2+^ to a receptor protein can be described with the theoretical framework developed for ligand-receptor complexes (Klotz, 2004), which confirms the experimentally observed sigmoidal curves. To substantially reduce the number of free parameters, the sigmoidal dependency can be satisfactorily approximated by the empirical Hill equation (Hill, 1910), which is in its most general form:

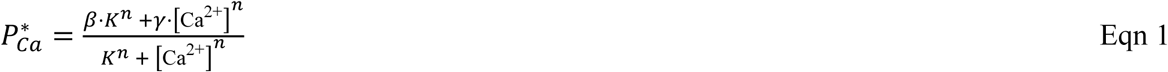

Here, [Ca^2+^] is the free calcium concentration, 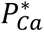 is the fraction of active proteins (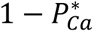 is the fraction of inactive protein), 0 ≤ *β* ≤ 1 is the fraction of active proteins at zero calcium, 0 ≤ *γ* ≤ 1 the fraction of active proteins under saturating calcium, *K* is the apparent dissociation constant that denotes the [Ca^2+^] at the midpoint of the curve 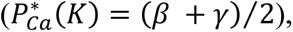 and *n* is the Hill coefficient that – together with *K*, *β* and *γ* – determines the slope of the sigmoidal curve at [Ca^2+^] = *K*: *slope*([Ca^2+^] = *K*) = *n* ⋅ (*γ* − *β*)⁄4*K* (**Figure 1a**, **Figure S1**). With *β* < *γ*, equation 1 describes the situation of a target protein that is activated by calcium (module 1, **Notes S1**), while with *β* > *γ* it represents module 2, where a target protein is inactivated by increasing calcium. Further considerations for more complex Ca^2+^-sensing modules 3-6 are provided in **Notes S1** (including **Figs. S1-S15** and **Tables S1-S3**) and in the following. Often, the activation of a target protein by Ca^2+^ is not instantaneous, but proceeds with a relevant delay that may be caused by a Ca^2+^-induced conformational change or the interaction of a Ca^2+^ binding protein with a target protein. In these cases, the activation of a target protein follows kinetics with an exponential time course

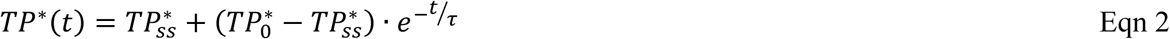

with the time constant τ, whose Ca^2+^-dependence can be specified by

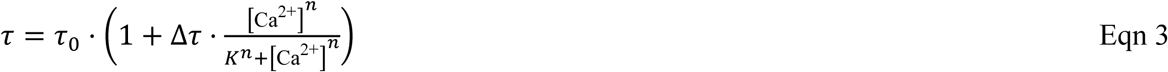

Here *τ*_0_ is the *τ*-value at zero Ca^2+^ and *τ*_0_ ⋅ (1 + Δ*τ*) the *τ*-value at very high Ca^2+^ (**Figure 1b**). Further details and comprehensive mathematical descriptions that lead to equations 2 and 3 are provided in the **Supplementary Material**.

**Figure 1.**
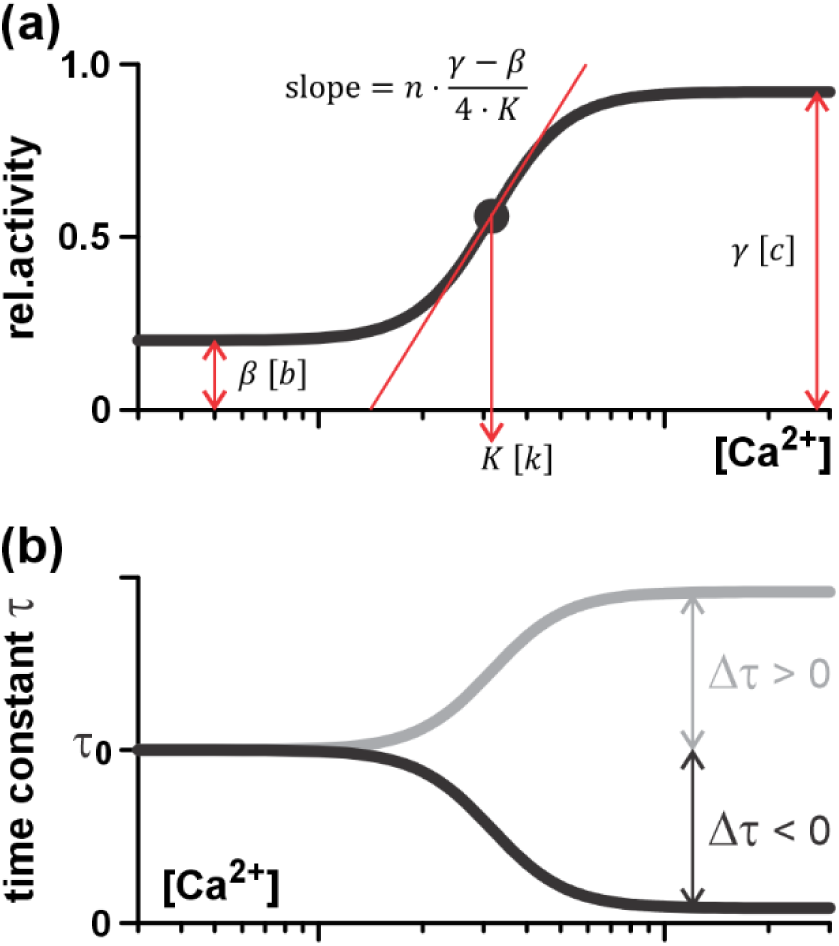
Fundamental characteristics of Ca^2+^-sensing modules. (**a**) In steady state the Ca^2+^-dependence of a target protein (TP) in a Ca^2+^-sensing module is characterized by four empiric parameters: *β* [*b*] represents the fraction of TPs at zero Ca^2+^, *γ* [*c*] the fraction at high Ca^2+^, *K* [*k*] is the [Ca^2+^] at the midpoint of the sigmoidal curve, and *n* is linked to the slope of the curve at its midpoint. *b*, *c*, *k* are the modified *β*, *γ*, *K* in modules 3-6 (Tabs. S1-S3). (**b**) For time-dependent modules, the time constant τ is Ca^2+^-dependent and is characterized by τ_0_, the value at zero Ca^2+^, and Δτ, the difference between τ at high Ca^2+^ and τ_0_. For modules 3 and 4 it is Δτ < 0, while Δτ > 0 holds for modules 5 and 6.

## Results

To gain insight into the fundamental basis of Ca^2+^ signal decoding at the mechanistic level, we chose a theoretical bottom-up approach using physical, chemical and mathematical principles. We described the binding of Ca^2+^ to perceptive proteins and the subsequent reactions with chemical reaction kinetics and analysed basic Ca^2+^-decoding modules, starting with the simplest and then increasing their complexity.

### Parameters for EF-hands differ from those of whole proteins

We started our bottom-up approach by modelling the simplest event in the signalling cascade that is the binding of a Ca^2+^ ion to an EF hand. This binding could be described by a simple chemical reaction: Ca^2+^ + EF ⇌ EF^*Ca*^. In steady state the fraction of EF hands bound with Ca^2+^ was given by EF^*Ca*^ = [Ca^2+^]⁄(*K* + [Ca^2+^]), i.e. Eqn. (1) with *β* = 0, *γ* = 1, *n* = 1 and an EF hand-dependent *K*-value. Usually, these *K*-values can be determined experimentally by calorimetry (Keeler *et al*., 2013) or computationally in molecular dynamics simulations (Project *et al*., 2006; Neamtu *et al*., 2023). Thus, the interaction of Ca^2+^ with EF hands in isolation could be described with *microscopic* parameters.

In the next step of complexity, we considered downstream effects in a calcium-binding protein (CBP), where calcium binding induces trans-conformation and activity changes. Typically, a CBP bears four EF hands arranged in two pairs, referred to as N- and C-lobes, with each of them capable of binding two calcium ions (**Figure 2**; DeFalco *et al*., 2010; Hashimoto & Kudla, 2011; Troilo *et al*., 2022). As explained before, the binding of Ca^2+^ to an EF hand could be characterized by an individual microscopic *K*-value (**Figure S18**; Bayley *et al*., 1984; Teleman *et al*., 1986; Christodoulous *et al*., 2004; La Verde *et al*., 2018; Troilo *et al*., 2022). If all EF hands are occupied, the CBP switches to an activated state, which could be described analogous to a chemical reaction with the forward and backward rates *k_a_* and *k_d_* respectively. Interestingly, the concerted interaction of all EF hands, regardless of whether Ca^2+^ binding occurred independently or cooperatively, consistently resulted in a sigmoidal S-shaped Ca^2+^ dependence of the CBP (**Figures 2, S18**, red curves), which could also be well described empirically by Eqn. (1) with – this time – *macroscopic* parameters (**Figures 2, S18**, blue curves). For illustration, **Figures 2** and **S18** show several examples of different, randomly selected *K_1-4_* and *k_a_*/*k_d_* values. Hence, Eqn. (1) provided an excellent description with a minimal set of parameters that eclipse the original microscopic parameters. Attempts to determine the microscopic parameters, including the *K*-values of the individual EF hands, by fitting the blue or red curves with the respective mathematical functions (**Figure S18**), did not lead to a reliable determination of these microscopic parameters. Remarkably, the macroscopic parameters γ (β), *K* and *n* were strongly influenced by the ratio *k_a_*/*k_d_* of the terminal reaction, but only slightly by the *K*-values of the EF hands (**Figure 2**; **Notes S2; Figures S16-S18**). Thus, the macroscopic parameters had a barely traceable relationship with the original microscopic parameters of the Ca^2+^ binding reaction(s) to the EF hands, which implies an important first conclusion: There are fundamental obstacles that make it nearly impossible to quantitatively correlate the Ca^2+^ binding properties of the EF hands and the Ca^2+^ sensitivity of the protein containing these EF hands. While the microscopic parameters (*K*-values of the EF hands) might be of interest from a biophysical perspective, they are less informative for physiological processes. The physiological processes are characterized by the empirical macroscopic parameters that depend also on other Ca^2+^ independent protein properties such as the *k_a_*/*k_d_* ratio.

**Figure 2.**
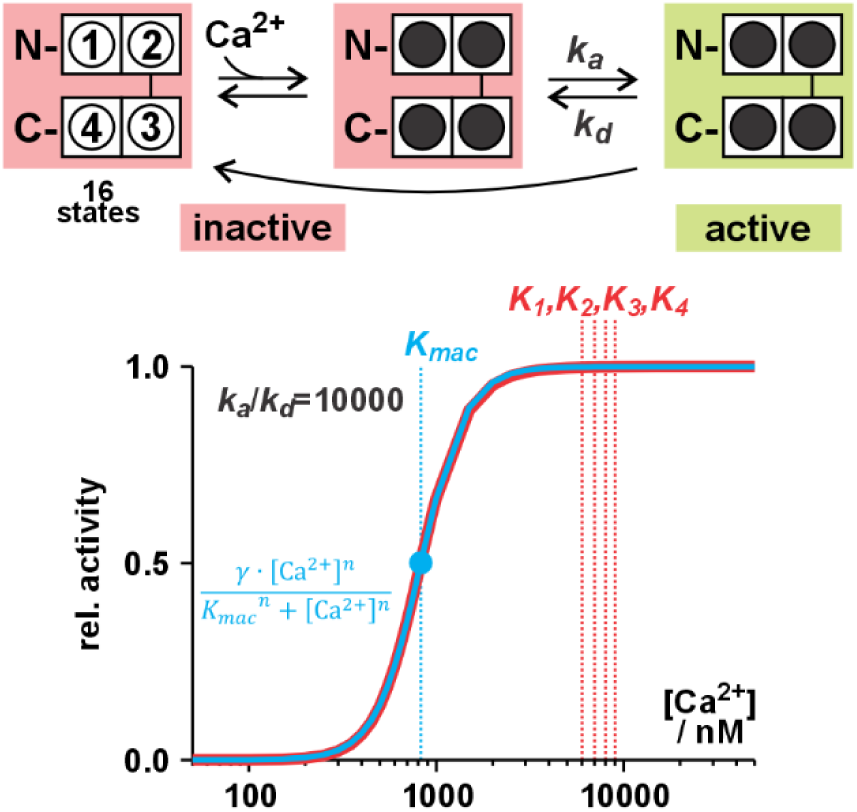
Ca^2+^-dependent activation of a protein with EF hands. Upper panel: A protein may contain four EF hands arranged in pairs in an N- and a C-lobe. The binding of Ca^2+^ to the EF hands can be described by individual *K*-values (*K_1_*, *K_2_*, *K_3_*, *K_4_*), resulting in 16 different states of the system. From the state in which all EF hands are occupied, the protein can transition to the activated state (*green*) through a further reaction with the rates *k_a_* and *k_d_*. Dissociation of Ca^2+^ renders the protein inactive. Lower panel: Example of a protein with *K_1_*=6μM, *K_2_*=8μM, *K_3_*=7μM, *K_4_*=6μM, and *k_a_*/*k_d_*=10000 (red curve). The resulting Ca^2+^-dependence of the activity of the protein could be well described by the empiric Eqn. (1) with the macroscopic parameters γ (β), *K_mac_* (=826.7 nM) and *n* (=3.61). These parameters are strongly influenced by the *k_a_*/*k_d_* ratio but only weakly by the *K*-values of the EF hands.

### Basic modules for Ca^2+^ signal decoding

As next, we examined the effects of calcium binding to proteins in more detail and identified six different fundamental Ca^2+^ decoding modules (modules 1–6; **Notes S1, Table 1**) formed by calcium-binding proteins and their target proteins. Each of the six has unique properties and can be described by a characteristic set of macroscopic parameters. In the following, these modules are presented systematically step by step.

**Table 1.**
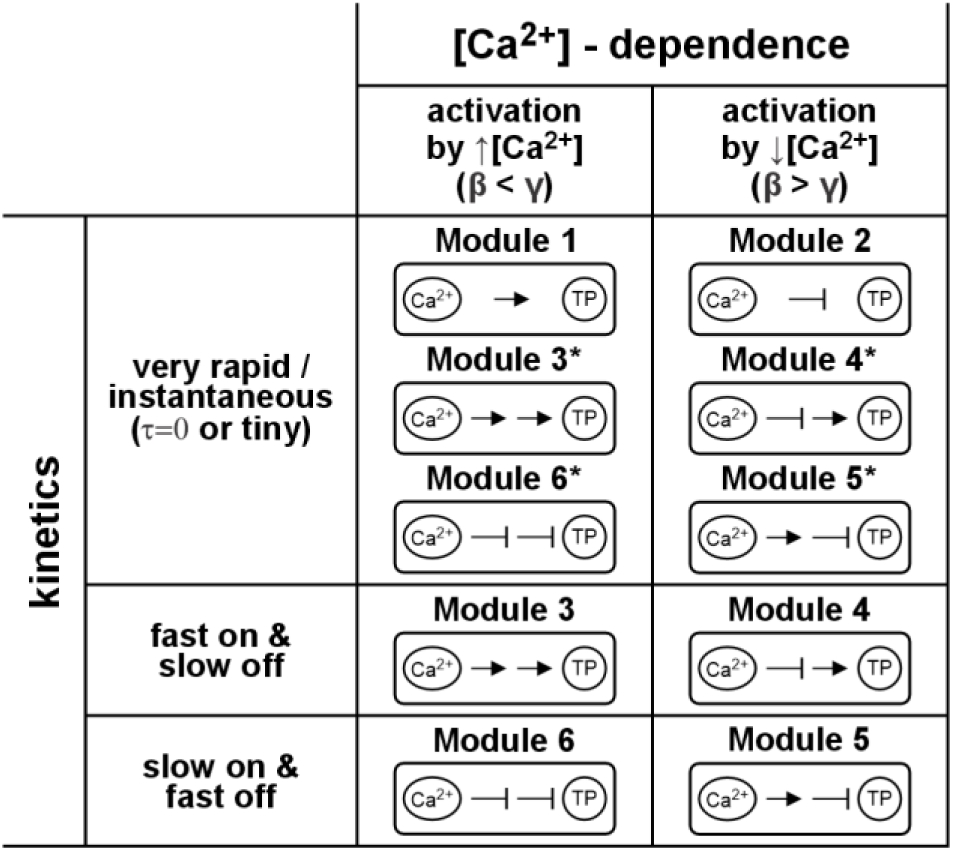
Classification of the six basic Ca^2+^ sensor modules. Calcium can influence a target protein (TP) by cascades of activating (→) or inactivating (⊣) steps. The basic units show one or two steps and allow to assemble six different modules: Module 1 (one step, →); Module 2 (one step, ⊣); Module 3 (two steps, →→); Module 4 (two steps, ⊣→); Module 5 (two steps, →⊣); Module 6 (two steps, ⊣⊣). According to their Ca^2+^-dependence the six modules can be distinguished into those that activate the TP with increasing [Ca^2+^] (↑, β < γ; modules 1, 3, 6) and others that activate the TP with decreasing [Ca^2+^] (↓, β > γ, modules 2, 4, 5). According to their kinetics, the response of a module to a change in [Ca^2+^] could be very rapid / almost instantaneous (modules 1, 2, 3*, 4*, 5*, 6*, τ=0 or very tiny) or could proceed with a relevant kinetical component (τ). In two of the modules, the time constant τ is larger at [Ca^2+^], at which the steady state activity of the TP is low, but smaller at [Ca^2+^], at which the activity of the TP is high. Here, a stimulating Ca^2+^-pulse causes a fast activation (fast on) and slower inactivation (slow off) after the stimulus (modules 3, 4). In the other two modules (5, 6) it is inverse. Here, an activating stimulus causes a slow on and a fast off response. * Please note that the time-dependent modules 3-6 could be very rapid that they appear to be almost instantaneous. In this case, they do not exhibit a physiologically important kinetic component. However, their parameters *β*[*b*], *γ*[*c*], *k*[*K*] might be modified by the interaction between CBP and TP (Tabs. S1-S3).

### Modules 1 and 2 can decode amplitudes but not frequency

At first, we considered the case that Ca^2+^ ions bind to a target protein (TP), which changed very rapidly its conformation and thus its activity (**Fig. S1**). Ca^2+^-binding could activate the TP (module 1) or inactivate it (module 2). From a mechanistic point of view, a Ca^2+^ ion binds to EF-hands in a TP and the occupation of several or all EF-hands resulted in the conformational/activity change of the protein. In steady-state conditions, which may establish almost instantaneously (<≈*μs*; HY Park *et al*., 2008; Swainsbury *et al*., 2012) with respect to physiological time scales (≥ *ms* - *min*), the Ca^2+^-dependence of the fraction of activated TP* could be described empirically according to the Hill equation (Eqn 1) with its four free parameters, as shown above. Due to the fast kinetics of the reactions, the activity of the target protein TP* immediately followed even a transient Ca^2+^-signal (**Figure 3**, case 1). In conclusion, these simple modules could extract information about the amplitude of a Ca^2+^ signal but they were not well suited to decode frequency information.

**Figure 3.**
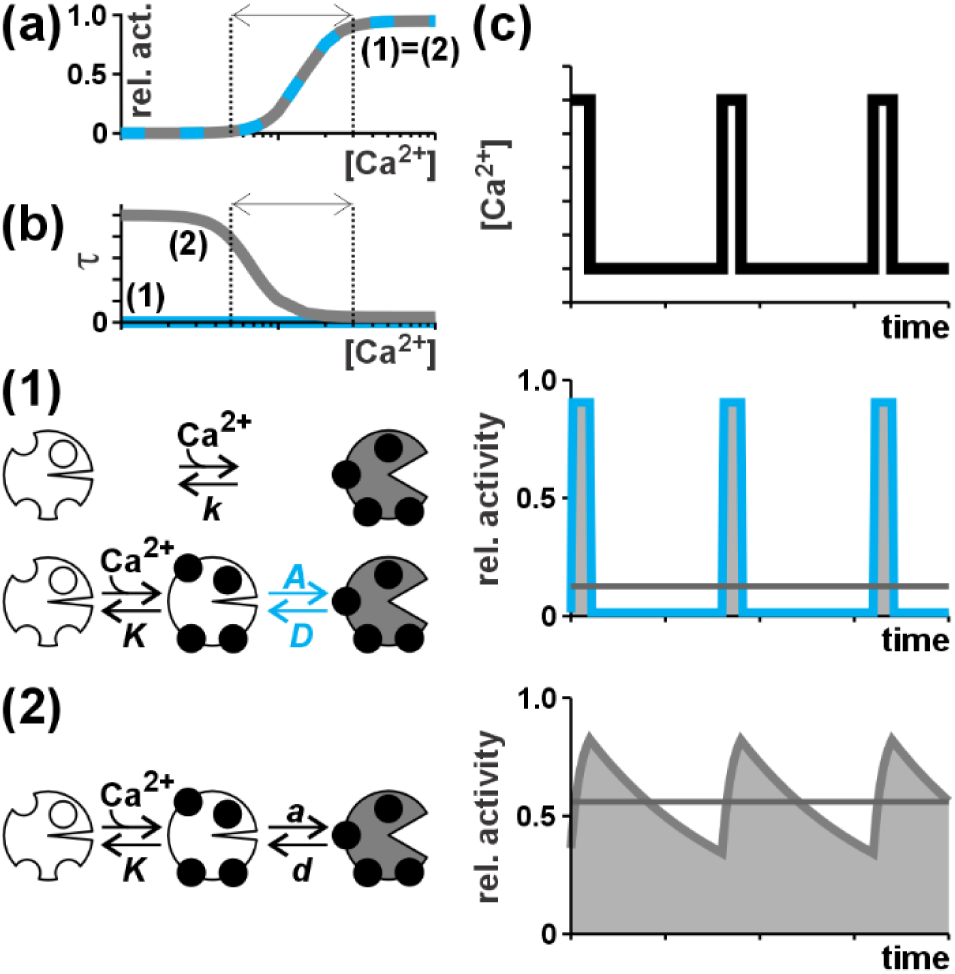
A slow intermediate step creates a kinetic component. (1) Ca^2+^-binding almost instantaneously activates the TP in a one-step process (parameter *k*) or in a very rapid two-step process (parameters *K*, *A*, *D*). (2) Ca^2+^-binding activates the target protein via a delaying intermediate step (parameters *K*, *a*<*A*, *d*<*D*). (a) Ca^2+^-dependence of the steady state activity of the target protein of the models (1, blue line) and (2, grey line). Note that both curves superimpose. (b) Ca^2+^-dependence of the time constant τ for both cases. Note, that for the instantaneous model (1) τ = 0 s (blue curve). (c) Response of the immediate (1, blue curve) and delayed activation cascade (2, grey curve) to Ca^2+^ pulses. The horizontal grey lines indicate the time-averaged activity level. The parameters for the simulations were *n*=4, *K*=3, *β*=0, *γ*=1, *a/d*=*A/D*=19, *d*=0.01 s^-1^, *D*=10 s^-1^ for the two step mechanisms and *n*=4, *k*=1.42, *β*=0, *γ*=0.95 for the one step mechanism. The Ca^2+^-states between which the system was switched are shown as dotted lines in (a) and (b).

### Modules 3-6: An intermediate step allows frequency decoding

Frequency information could be better decoded, when the Ca^2+^-binding triggered a subsequent rather slow reaction. Such a delay between Ca^2+^-binding and a change in activity could be established in three different ways (**Notes S1**): (i) Rapid, direct binding of Ca^2+^ to the TP, followed by a slower subsequent modulatory effect such as a conformational change. An example would be the binding of Ca^2+^ to the EF-hands of the ion channel TPC1 that changes the conformation of the channel near the gate and voltage sensor (Kintzer & Stroud, 2016; Mérida-Quesada *et al*., 2022). (ii) A Ca^2+^-binding protein (CBP) is rapidly modulated by Ca^2+^ (Modules 1 and 2), and then, during a slower step, the CBP catalytically modulates the activity of the TP. An example is the Ca^2+^-dependent phosphorylation of the Arabidopsis Vacuolar K^+^ Channel TPK1 by the Ca^2+^-dependent protein kinase CPK29 (Latz *et al*., 2013). (iii) A CBP is rapidly modulated by Ca^2+^ (Modules 1 and 2), after which it binds to the TP and modulates its activity through binding. An example is the binding of calmodulin to the ion channel CNGC15 (Del Cerro *et al*., 2022). Although the mathematical descriptions of scenarios (i)-(iii) differed in detail, the physiologically relevant outcome of the modulation of a TP by Ca^2+^ was very similar for all three (**Notes S1**). In particular, the Ca^2+^ sensitivity in steady state could be represented again by the Hill equation (Eqn. 1), while the time course of activation/inactivation could be described by an exponential decay kinetics as 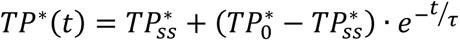 with the Ca^2+^-dependent time constant τ (Eqn. 2). Here *TP*^∗^(*t*) describes the time progression of the fraction of active *TP*, while 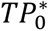 and 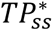 denote the fraction of active TP at *t*=0 and in steady state, respectively.

An example of the effect of a slower intermediate step is shown in **Figure 3**. Calcium bound to the target protein and activated it (1) almost instantaneously or (2) in a subsequent, slower reaction. In this example, we chose conditions so that the Ca^2+^-dependent steady-state activity of the TP was identical in both cases (**Figure 3a**). The first case could be modelled in two ways. Either as a direct immediate reaction, or as a two-stage process in which the second step is very fast. In both cases, the activity of the TP directly followed the Ca^2+^ signal (**Figure 3c, case (1)**). This characteristic changed fundamentally when the second step was slowed down. The time course of the TP activity then had its own kinetic component (**Figure 3c, case (2)**), which was characterised by the time constant τ (**Figure 3b**) and thus by the parameters *a* and *d* of the slower reaction (**Notes S1**).

The intermediate step did not only cause a new kinetic component, but introduced also a further combinatorial degree. While Ca^2+^-binding can activate (→) or inactivate (⊣) in the first step, the following step can also have activating (→) or inactivating (⊣) effects. This allowed to form four additional Ca^2+^-decoding modules (Module 3 →→, Module 4 ⊣→, Module 5 →⊣, Module 6 ⊣⊣, **Table 1**). The modules distinguished in the Ca^2+^ dependence of their activation and their kinetics. In two of them (modules 3 and 6) the TP was activated by an increase in Ca^2+^, while in the other two (modules 4 and 5) the TP was inactivated by Ca^2+^. With respect to the kinetics, two of the modules activated faster than they inactivated (“fast on / slow off”, modules 3 and 4), while the other two inactivated faster than they activated (“slow on / fast off”, modules 5 and 6). Irrespective of the different implementations (modules 1, 2, i.3-i.6, ii.3-ii.6, iii.3-iii.6, **Notes S1**) the six modules with their peculiar functional properties provide the universal foundation for Ca^2+^-signal decoding in plants and likely also in other organisms.

### Ca^2+^ sensitivity of kinases is substrate dependent

Having established the fundamentals of the modules, we continued to use them to investigate physiologically relevant situations, in particular some seemingly puzzling experimental observations. One such, at first glance incomprehensible finding is that Ca^2+^-dependent kinases exhibit different Ca^2+^-sensitivities when tested with different substrates (JY Lee *et al*., 1998; Harper *et al*., 2004). This phenomenon could be explained with module ii.3 (**Notes S1**, **Figure S6**, **Table S2, Figure 4**), in which the kinase was represented by the CBP (characterized by a Hill function with the parameters *β*, *γ*, *K* and *n*) that catalytically interacted with the substrate represented by TP. The experimental readout is usually the fraction of phosphorylated substrate at different free [Ca^2+^] (TP*), which could again be described by a Hill function (parameters *b*: TP* at zero Ca^2+^; *c*: TP* at high Ca^2+^; *k*: apparent dissociation constant; *n*: Hill coefficient). The reaction between CBP and TP was essentially determined by the rate constants *a* and *d* that described the affinity between the kinase and the substrate. Different substrates differed in their affinity to the kinase and showed consequently different values for *a* and *d*. The values of both, however, influenced the parameters *b*, *c*, and *k* (**Table S2**, **Figure 4**). This implies that the values obtained from the TP*-readout do not represent the Ca^2+^-dependent features of the kinase (*β*, *γ*, *K*, *n*) but rather are a measure of the interaction between the kinase and the substrate. Additionally, the parameter *a* contained implicitly the factor *CBP_total_*, *i.e.* the expression level of the CBP (**Notes S1**). Thus, the fraction of phosphorylated substrate depends not only on the affinity of the kinase for the substrate, but also on the amount of kinase present in the reaction. The conclusion from this consideration is that a kinase can hardly be characterized by a unique Ca^2+^ sensitivity, and that one and the same kinase can exhibit different target-dependent Ca^2+^ sensitivities in cellular processes.

**Figure 4.**
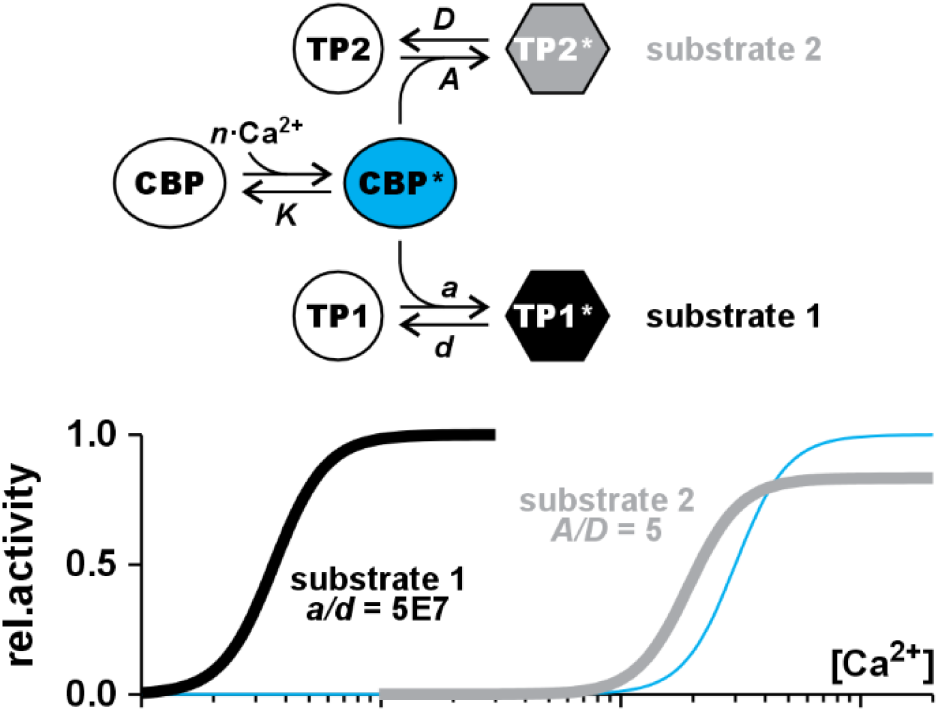
Substrate-dependent apparent Ca^2+^-sensitivity. The same calcium binding protein (CBP, *e.g.* a kinase) can interact with different target proteins (TP1, TP2) as substrates; here shown for module 3 (Ca^2+^→→TP). The Ca^2+^-dependent activity of the substrate in steady state strongly depends on the affinity between CBP and TP. A higher affinity results in an apparent higher Ca^2+^-sensitivity. The blue curve shows the Ca^2+^-sensitivity of the CBP. For illustration, the parameters were *n*=4, *K*=3, *β*=0, *γ*=1. The interaction with substrate 1 (*black*) is shown with *a/d*=5×10^7^, while for substrate 2 (*grey*) the affinity is selected as *A/D*=5. The example demonstrates that the activity curve of the TP is not a measure for the Ca^2+^-sensitivity of the CBP.

We based our conclusion on an example from module ii.3, but similar conclusions could also be drawn from all other modules. The comprehensive considerations in **Notes S1** further demonstrated that this conclusion is valid for all other CBPs that exerted an effect on a target protein, e.g. CBLs, CMLs, apoCMLs. The parameters *a* and *d*, which are a measure of the interaction between CBP and TP, did not only affect the steady state levels of TP* but influenced also the kinetics of the reaction. The higher the affinity between CBP and TP, the more pronounced the difference in the kinetics between low and high Ca^2+^, and the more pronounced the effects that will be presented in the following.

### Disturbance tolerance of Ca^2+^-sensing modules

To gain further insight into the dynamics of the different modules, it was useful to analyse their behaviour under controlled conditions, e.g. a constant Ca^2+^-amplitude or stepwise changes in amplitude since a transient Ca^2+^ signal could be interpreted as a superimposition of different Ca^2+^ pulses with constant amplitudes. Following a Ca^2+^ pulse, the time course followed by the fraction TP* was generally an exponential decay function (Eqn. 2) with the characteristic parameters TP*_ss_, the steady-state value of TP*, and τ, the time constant of the exponential course.

Among the four slow (not instantaneous) modules, modules 3 and 6 were both activated by increasing [Ca^2+^] (**Figure 5**). However, they differed in their kinetic features. Upon a stepwise change from lower to higher [Ca^2+^] and back, module 3 reacted faster to the Ca^2+^-increase (fast ‘on’) than module 6 (slow ‘on’), but it responded more slowly to a Ca^2+^-decrease (slow vs. fast ‘off’; **Figure 5a**). This difference in dynamics had fundamental consequences under conditions of fluctuating Ca^2+^. During Ca^2+^ spiking pulses (**Figure 5b**), the fast ‘off’-reaction allowed module 6 to relax quickly during inter-spike periods, resetting the system for the subsequent pulse. In comparison, module 3 did not relax as quickly and accumulated, over time, a new background activity that was higher than the basic level (**Figure 5b**, grey area). Besides, this inertial behavior rendered module 3 more robust towards disturbances in long lasting pulses than module 6 as illustrated in another simple example (**Figure 5c**). In long lasting Ca^2+^-pulses, the activity of TP* reached a Ca^2+^ amplitude-dependent steady-state (**Figure 5c**, black lines). Interruptions in such a pulse (**Figure 5c**, top, grey curve) resulted in a transient inactivation that was more pronounced for module 6, with its fast ‘off’ kinetics, than for module 3.

**Figure 5.**
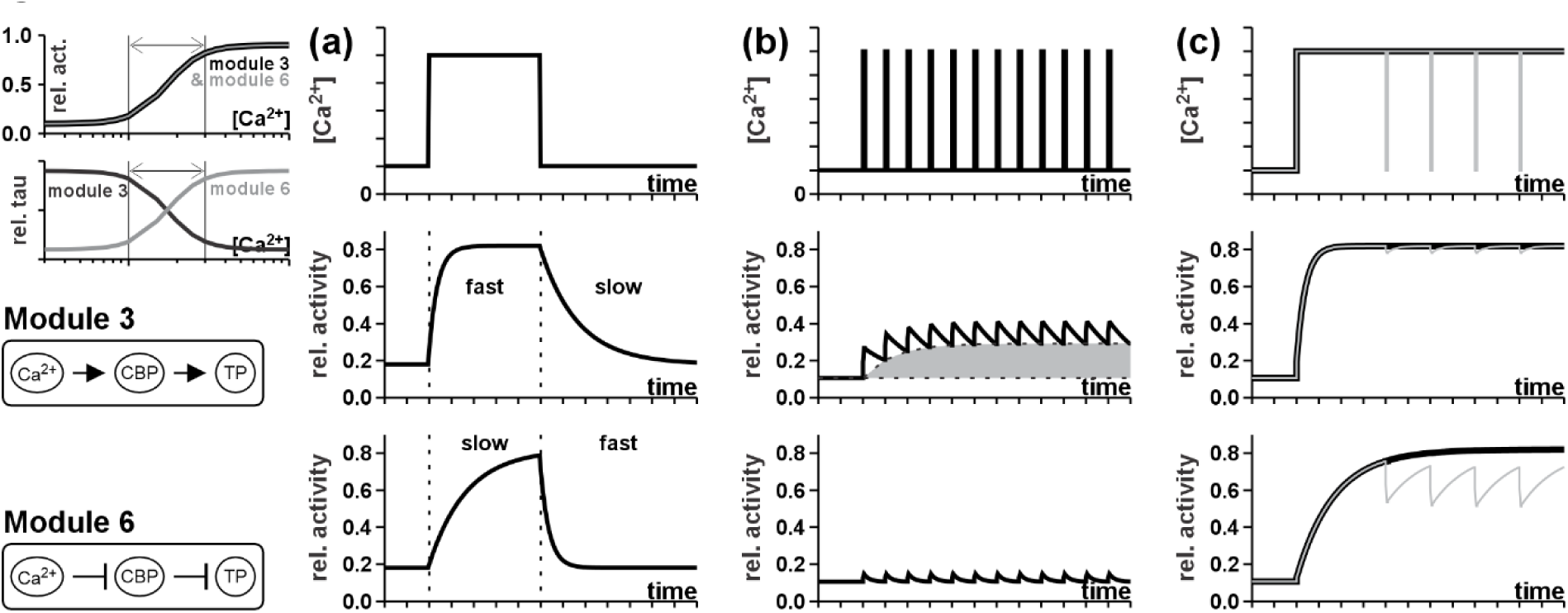
Comparison of the dynamic behavior of the Ca^2+^-sensing modules 3 and 6. In both modules the target protein (TP) is activated by rising Ca^2+^-concentrations (top left). The two modules differ in their kinetic response to fluctuating Ca^2+^-concentrations. (**a**) The module 3 (middle) reacts more quickly to a stimulating than to a relaxation Ca^2+^-pulse (upper panel). In contrast, module 6 (bottom) responds more slowly to an activating than to an inactivating pulse. (**b**) In response to Ca^2+^-spikes, the slow relaxation of module 3 can result in the accumulation of a frequency-dependent background activity of the TP (grey area), while, under identical conditions, the module 6 still allows the relaxation to the baseline. (**c**) Module 6 is more sensitive towards disturbances in long lasting Ca^2+^-pulses than module 3. For illustration, the long-lasting Ca^2+^-pulse (black line) was interrupted by 4 flickering events of low Ca^2+^ (grey line). To allow a comparison of the two modules, parameters *n*, *b*, *c*, *k*, τ_min_ and τ_max_ were identical in both cases.

A similar comparison of two modules could also be made between module 4 (fast on / slow off) and module 5 (slow on / fast off) (**Figure 6**). In these cases, however, a Ca^2+^ pulse inactivated the TP. Interestingly, module 4 represents the mechanism by which apo-calmodulin modulates the activity of the heteromeric CNGC8-CNGC18 (Pan *et al*., 2019). When apo-calmodulin binds Ca^2+^, it dissociates from the CNGC, leading to a decrease in the relative activity of this target protein. Conversely, module 5 represents the mechanism by which Ca^2+^ activates calmodulin 2 to modulate the CNGC15 (Del Cerro *et al*., 2022). Once calmodulin 2 binds Ca^2+^, it adopts its active conformation and binds to the IQ motif of the CNGC15, closing it. In both cases, Ca^2+^ had an inhibitory effect on the target protein (CNGC). Nevertheless, the two mechanisms differed in their dynamic properties. While module 4 was more robust to flickering Ca^2+^ spikes than module 5 (**Figure 6b**), it was more sensitive to interruptions in long-lasting pulses (**Figure 6c**). These mechanistic insights suggested that Ca^2+^-bound calmodulin 2 and apo-calmodulin affect similar target proteins (in this case CNGC8/18 and CNGC15) with different dynamics, although both have the same final effect on the TP, namely the Ca^2+^-induced inactivation. During Ca^2+^ fluctuations, Ca^2+^-bound calmodulin inactivates the TP faster than it activates it, while apo-calmodulin activates the TP faster than it inactivates it.

**Figure 6.**
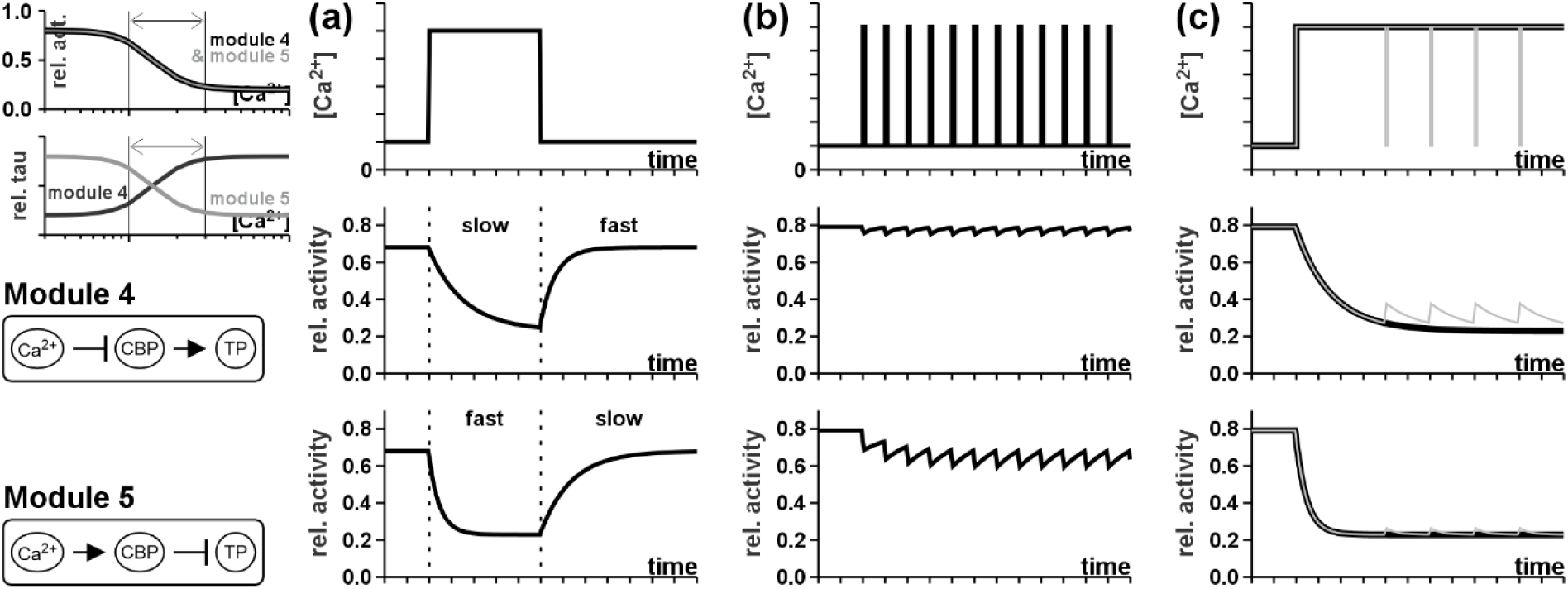
Comparison of the dynamic behavior of the Ca^2+^-sensing modules 4 and 5. In both modules the target protein (TP) is deactivated by rising Ca^2+^-concentrations (top left). The two modules differ in their kinetic response to fluctuating Ca^2+^-concentrations. (**a**) The module 4 (middle) reacts more quickly to a stimulating than to a relaxation Ca^2+^-pulse (upper panel). In contrast, module 5 (bottom) responds more slowly to an activating than to an inactivating pulse. (**b**) In response to Ca^2+^-spikes, the module 4 is hardly affected, while, under identical conditions, the module 5 is more sensitive to the short Ca^2+^ pulses. (**c**) Module 4 is more sensitive towards disturbances in long lasting Ca^2+^-pulses than module 5. For illustration, the long-lasting Ca^2+^-pulse (black line) was interrupted by 4 flickering events of low Ca^2+^ (grey line). To allow a comparison of the two modules, parameters *n*, *b*, *c*, *k*, τ_min_ and τ_max_ were identical in both cases.

### Frequency dependence of Ca^2+^-sensing modules

We next investigated the behavior of the modules upon repetitive Ca^2+^ pulses with different durations and different inter-pulse intervals (**Figure 7**, **Figures S19-S21**). Usually, cellular processes are rather slow compared to the rapid fluctuations in cytosolic [Ca^2+^]. For example, calcium concentrations oscillate within minutes during the growth of moss protonema, pollen tubes, and root hairs, regulating an apical cell growth process that lasts for hours. The same can be observed in root cells, where salt stress induces calcium oscillations, or in calcium oscillations in guard cells, which modulate stomatal opening (McAinsh *et al*., 1997; Shabala & Newman, 1997; Cárdenas *et al*., 2008; Bascom *et al*., 2018; Ogawa *et al*., 2025;). Therefore, it is not the real-time progression of TP activity (**Figure 7b**, grey surface) that is decisive, but rather its average activity over time (**Figure 7b**, horizontal line). This average activity strongly depends on the frequency of the pulses (**Figure 7**). In the case of module 3, the average activity of TP increased when the time interval between spikes shortened (**Figure 7b**, ①**→**②), and when the spike duration increased (**Figure 7b**, ③**→**④). Equivalent results were obtained for module 6 (**Figure S21**). Modules 4 and 5 showed inverse behaviour (**Figures S19, S20**). Here, the average activity of TP increased when the time interval between spikes increased, and when the spike duration shortened. Thus, the modules react on both the duration of the spike and the inter-spike interval, albeit inversely, and already allow the decoding of frequency features of the Ca^2+^ signal by converting it into a frequency-dependent average activity of the TP.

**Figure 7.**
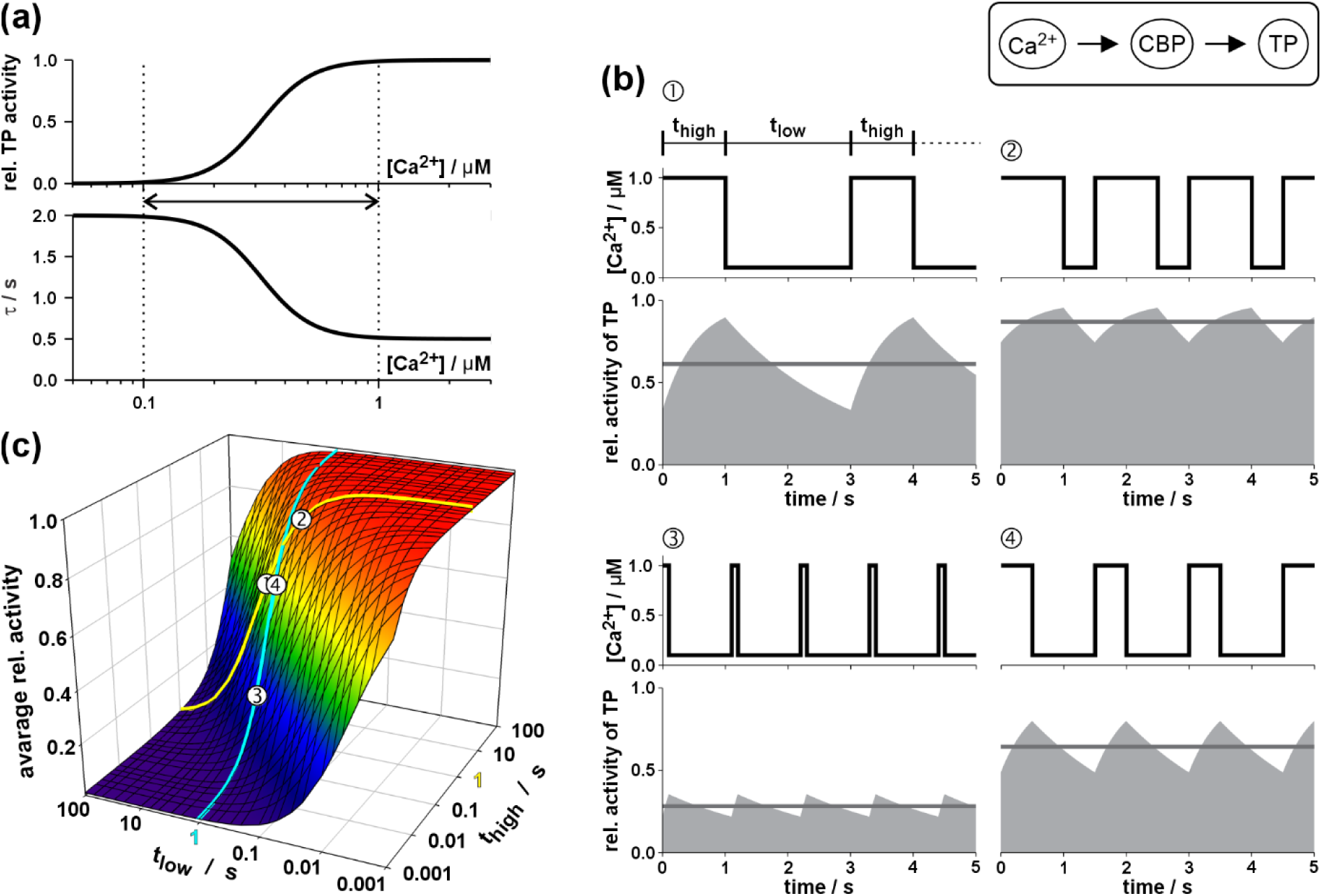
Ca^2+^-sensing modules allow adjusting the activity of a target protein (TP) towards the frequency of a flickering Ca^2+^-signal. Exemplarily module 3 is considered. The responses of the other three modules with comparable parameters for TP and τ are shown in Figures S19-S21. (a) Ca^2+^ dependence of the steady state activity of the target protein (TP) and the time constant (τ) of the relaxation processes. The dashed lines indicate the Ca^2+^ values between which oscillations occurred in the various scenarios. (b) Four scenarios with repetitive pulses between high and low Ca^2+^ (①-④) are shown with different durations t_high_ and t_low_. The average activity of TP over time (grey line) is given by the time-average over the grey surface under the curve. (c) 3D-plot of the average relative activity of TP as a function of the two parameters t_high_ and t_low_. The data of the four presented scenarios are indicated and cross-referenced by numbers ①-④. For better orientation the values for t_high_ = 1s and t_low_ = 1s are indicated by a yellow and a cyan line, respectively.

Some characteristics of Ca^2+^-sensing modules in response to oscillating Ca^2+^ signals have already been clearly described in earlier studies of individual cases of Ca^2+^-dependent protein phosphorylation (Goldbeter *et al*., 1990; Dupont & Goldbeter, 1998; Prank *et al*., 1998; Gall *et al*., 2000; Larsen *et al*., 2004; Schuster *et al*., 2005). Here, we have generalized these cases and placed them on a solid theoretical foundation. Our conclusion is that these modules are not yet optimal for frequency decoding on their own. In case of module 3 the average TP activity increased with longer spikes (longer t_high_; **Figure 7c**, cyan curve), but also with shorter inter-spike intervals (shorter t_low_; **Figure 7c**, yellow curve). The model presented here demonstrates that decreasing spike frequency while increasing spike duration produces a similar increase in TP activity to that obtained by increasing frequency while maintaining the same spike duration.

### Combinations of modules can specify the resonance interval

In the final step, we assessed the effect of combining the different modules. Indeed, by combining two modules with opposing characteristics, e.g. modules 3&5, the frequency range within which a TP can be activated was significantly restricted (**Figure 8**). Similar results were obtained for the combinations 3&4, 6&4, and 6&5 (**Figure S22**). The opposite properties of the modules define an optimal range of Ca^2+^ fluctuations within which TP exhibits its highest activity. Interestingly, such combinations of modules actually exist *in vivo*. Recently, Ming *et al*. (2025) described the activation of CNGC5/6 channels by phosphorylation via the Ca^2+^-dependent protein kinase CPK3 (module 3), together with an inhibitory effect of CaM2 binding to the C-terminus of the CNGCs (module 5). Thus, the modules presented in this study are fundamental entities for decoding Ca^2+^-signals in plants, and likely also in other organisms.

**Figure 8.**
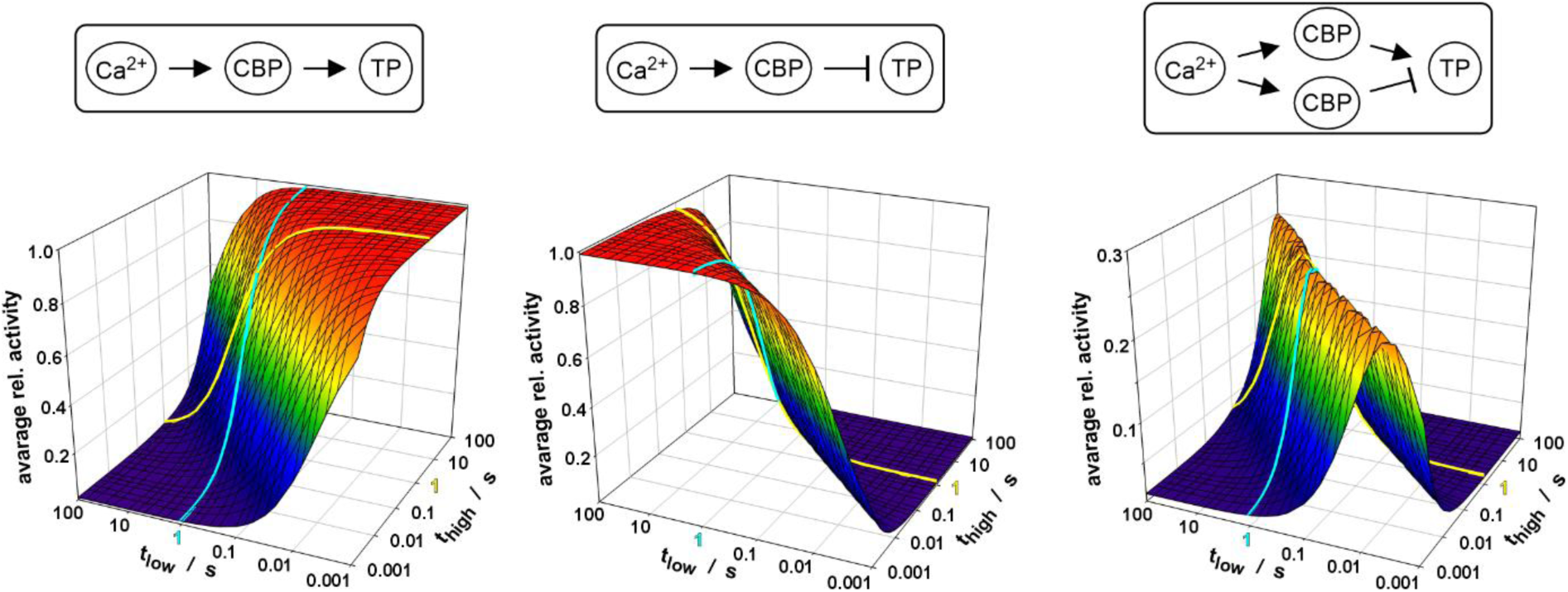
Combination of Ca^2+^ decoding modules enhance the frequency sensitivity. While module 3 (left) cannot distinguish between longer spikes (larger t_high_, cyan) and shorter inter-spike intervals (smaller t_low_, yellow), module 5 (middle) cannot distinguish between longer inter-spike intervals (larger t_low_, yellow) and shorter spikes (smaller t_high_, cyan). Combination of both modules (right) results in an optimum for the t_high_/t_low_ ratio.

## Discussion

Calcium (Ca^2+^) is an important second messenger whose mode of action is still only partially understood. The generation of calcium signals is well documented, with several calcium-permeable channels having been identified (Jin *et al*., 2025). The ongoing work of several groups on the fine tuning of these calcium channel activities will inevitably contribute to our understanding of calcium signalling and calcium signature formation. More elusive are the processes involved in calcium signal decoding in plants. In some very well-described experimental models, including pollen tubes, root hairs, guard cells and root cells, calcium oscillations are frequently observed, but their function is controversial and still largely unclear. One way to understand this fundamental aspect of plant physiology is to decipher the molecular mechanism of calcium signal decoding in order to have the basic tools for genetic and physiological experiments. In this study, we complemented these endeavours by adopting a theoretical bottom-up approach to gain insight into the mechanisms by which Ca^2+^ amplitudes and fluctuations can influence protein activity.

Although our approaches were developed for Ca^2+^, they are by no means limited to this ion. The comprehensive theory presented in the supplementary material can also be applied almost 1:1 to other ions or molecules that regulate protein function through binding reactions, such as abscisic acid that acts in a module-5-like manner via PYR/PYL/RCAR receptors on type 2C protein phosphatases (SY Park *et al*., 2009), or zinc, the most common metallic cofactor (Krämer, 2025). Although in these examples the time-dependence, a dynamic property of the universal modules, may not be as important as for Ca^2+^, the steady-state characteristics of TP are significant. Therefore, the conclusions apply not only for Ca^2+^ but also to other ions and molecules. In this context, it may be noted that H2O_2_ and pH signaling are also based on transient or oscillating signal signatures.

### A heuristic macroscopic equation covers a multitude of microscopic realities

The first important conclusion is that there is no quantitatively deducible, unambiguous correlation between the Ca^2+^-binding properties of EF-hands and the activity of the protein containing these EF-hands. Although both can be described by the sigmoidal Hill equation (Eqn 1), for isolated EF-hands the parameter *K* reflects the binding reaction and therefore has a biophysical significance, whereas for the whole protein, *K* is an empiric parameter. It depends, in a complex manner, not only on the binding reaction between Ca^2+^ and the EF hands but predominantly on the conformational changes that activate the protein after Ca^2+^ binding. Thus, even if all biophysical Ca^2+^-binding properties of the EF hands could be experimentally determined, the final activation reaction remains a largely inaccessible black box. The mathematical background presented in this study explains the elegant way out of this dilemma: Combinations of sigmoidal functions can be described again by a sigmoidal function (**Notes S2**) resulting finally in the empiric Hill equation that is often used as a satisfying heuristic description of the Ca^2+^ sensitivity of a CBP (Zauser *et al*., 2025). It is evident, however, that a Hill equation with 4 parameters is not unambiguous and can represent a multitude of microscopic realities.

### Ca^2+^-sensitivity is not an unambiguous property of a Ca^2+^ binding protein

Another important conclusion is that a CBP, like CPK, CBL or CML can hardly be characterized by a unique Ca^2+^ sensitivity, but that one and the same CBP can exhibit different target-dependent Ca^2+^ sensitivities in cellular processes. In wet-lab approaches, the Ca^2+^ sensitivity of a CBP is usually determined with a standard target protein as readout. The analyses presented in this study showed that it is predominantly the protein-protein interaction between CBP and TP (parameters *a* and *d*) that determines the Ca^2+^-dependent activity of the TP but not the original *K*-value of the CBP. The characterization of Ca^2+^ sensitivities of CBPs, such as kinases, with standard substrates may therefore not allow conclusions about their actual role *in vivo*. The same kinase may be characterized in biochemical assays as largely Ca^2+^ insensitive with one substrate but as highly Ca^2+^ sensitive with another (**Figure 4**). Furthermore, a higher amount of kinase is likely to result in greater apparent Ca^2+^ sensitivity than a lower amount (**Notes S1**).

### The key to understanding Ca^2+^ signal decoding lies at a meta-level

The theoretical findings presented in this study suggest that neither microscopic parameters of Ca^2+^ binding nor parameters of individual proteins are of primary importance for understanding the physiological aspects of Ca^2+^ signal decoding. Instead, there are universal general structures at a meta-level that exhibit peculiar emergent properties. We identified six different basic Ca^2+^ decoding modules (**Table 1**) formed by Ca^2+^ -binding proteins and their target proteins. Each of the six has unique properties and may be described at the meta-level by a characteristic set of parameters (*k*, *b*, *c*, *n*, *τ_0_*, *τ_∞_*, *k_τ_*, *n_τ_*). These parameters are not determined by the properties of a single protein, but by all units of the modules and the interactions between them. Thus, a Ca^2+^-dependent protein kinase or calmodulin may behave differently with different partners and in different modules, allowing great flexibility in setting the physiologically important parameters. The values for the steady-state activity of the TP at very low Ca^2+^ (parameter *b*) and at very high Ca^2+^ (parameter *c*), the slope of the Ca^2+^ dependence of the steady-state activity curve (*n*), and its midpoint Ca^2+^-value (*k*), together with the parameters of the time constant τ (*τ_0_*, Δ*τ* = *τ_∞_* - *τ_0_*, *k_τ_*, *n_τ_*), which could additionally be tuned via their expression levels (**Notes S1**), determine the sensitivity of the target protein not only to the amplitude of Ca^2+^ oscillations, but also to their frequencies.

### Combination of modules allow frequency fine-tuning

The six types of modules can be considered as fundamental building blocks that are assembled *in vivo* into more complex units in order to further optimize, refine, and adapt the Ca^2+^ decoding features to physiological requirements. For instance, two type 3 modules were found to be combined to a bi-kinase module with sensitized signaling properties (Köster *et al*., 2025). Another example is a combination of type 3 and type 5 modules involved in the regulation of CNGC5/6 channels (Ming *et al*., 2025), which shows enhanced sensitivity to the frequency of Ca^2+^ signals (**Figure 8**). These few examples of module combinations have only recently been reported, indicating that we are still in the early stages of identifying such complex structures. This makes it all the more important that these wet-lab findings are accompanied by robust theoretical dry-lab analyses, so that we can understand the complexity in depth. In this study, we have established a comprehensive theoretical foundation for decoding Ca^2+^ signals based on first principles, paving the way for future detailed analyses of module combinations. The continued parallel work in wet labs and dry labs is very promising for ultimately finding the “Rosetta Stone” for Ca^2+^ signals in plants.

## Acknowledgments

This work was supported by grants from the Agencia Nacional de Investigación y Desarrollo de Chile (ANID; grants No. 21220419 to F.V.-V., Fondecyt Regular No. 1220504, and Anillo No. ATE220043 to I.D., Fondecyt Regular No. 1251753 to E.M., Fondecyt Iniciacion No. 11251254 to M.E.R.-M.), and from the Biotechnology and Biological Research Council (BBSRC; grant No. BB/Z516843/1 to M.C.).

## Author contributions

F.V.-V. and I.D. conceived the study. All authors contributed to the design of the study. F.V.-V. and I.D. analysed the data. M.E.R.-M. participated in data visualization. F.V.-V. wrote the first draft of the manuscript. All authors contributed to writing and editing of the manuscript and approved its final version.

## Data availability

Further details about the modeling approach are available from Ingo Dreyer.

## Supporting Information

Additional Supporting Information may be found online in the Supporting Information section at the end of the article.

**Notes S1** with **Figs. S1-S15** and **Tables S1-S3** Ca^2+^-dependence of the Ca^2+^-decoding modules 1-6.

**Notes S2** with **Figs. S16-S17** Sigmoidal curves and the universal empiric Hill equation.

**Fig. S18** Mathematical description of Ca^2+^ sensitivity with a minimal set of parameters.

**Fig. S19** Frequency dependence of module 4.

**Fig. S20** Frequency dependence of module 5.

**Fig. S21** Frequency dependence of module 6.

**Fig. S22** Frequency dependence of combined modules.

## Abbreviations

TP: target protein
CBP: Ca^2+^-binding protein
CaM: calmodulin
apoCaM: apo-calmodulin
CML: calmodulin-like protein
CDPK: Ca^2+^-dependent protein kinase
CBL: calcineurin B-like (CBL) protein
CIPK: calcineurin B-like interacting protein kinase;
CCaMK: Ca^2+^/calmodulin-dependent protein kinase.

## References

Adams, P. J., Ben-Johny, M., Dick, I. E., Inoue, T., & Yue, D. T. (2014). Apocalmodulin itself promotes ion channel opening and Ca2+ regulation. Cell, 159(3), 608–622. 10.1016/j.cell.2014.09.047

Allan, C., Morris, R. J., & Meisrimler, C. N. (2022). Encoding, transmission, decoding, and specificity of calcium signals in plants. Journal of Experimental Botany, 73(11), 3372–3385. 10.1093/JXB/ERAC105

Bascom, C. S., Winship, L. J., & Bezanilla, M. (2018). Simultaneous imaging and functional studies reveal a tight correlation between calcium and actin networks. Proc. Natl. Acad. Sci. U.S.A., 115(12), E2869–E2878. 10.1073/PNAS.1711037115

Bayley, P., Ahlstrom, P., Martin, S. R., & Forsen, S. (1984). The kinetics of calcium binding to calmodulin: Quin 2 and ANS stopped-flow fluorescence studies. Biochemical and Biophysical Research Communications, 120(1), 185–191. 10.1016/0006-291X(84)91431-1

Brownlee, C., & Wheeler, G. L. (2025). Cellular calcium homeostasis and regulation of its dynamic perturbation. Quantitative Plant Biology, 6, e5. 10.1017/QPB.2025.2

Carafoli, E., & Krebs, J. (2016). Why calcium? How calcium became the best communicator. Journal of Biological Chemistry, 291(40), 20849–20857. 10.1074/jbc.R116.735894

Cárdenas, L., Lovy-Wheeler, A., Kunkel, J. G., & Hepler, P. K. (2008). Pollen Tube Growth Oscillations and Intracellular Calcium Levels Are Reversibly Modulated by Actin Polymerization. Plant Physiology, 146(4), 1611–1621. 10.1104/PP.107.113035

Christodoulous, J., Malmendal, A., Harper, J. F., & Chazin, W. J. (2004). Evidence for Differing Roles for Each Lobe of the Calmodulin-like Domain in a Calcium-dependent Protein Kinase. Journal of Biological Chemistry, 279(28), 29092–29100. 10.1074/JBC.M401297200

Contador-Álvarez, L., Rojas-Rocco, T., Rodríguez-Gómez, T., Rubio-Meléndez, M. E., Riedelsberger, J., Michard, E., & Dreyer, I. (2025). Dynamics of homeostats: the basis of electrical, chemical, hydraulic, pH and calcium signaling in plants. Quantitative Plant Biology, 6, e8. 10.1017/QPB.2025.6

Cook, N. M., Gobbato, G., Jacott, C. N., Marchal, C., Hsieh, C. Y., Lam, A. H. C., Simmonds, J., del Cerro, P., Gomez, P. N., Rodney, C., Cruz-Mireles, N., Uauy, C., Haerty, W., Lawson, D. M., & Charpentier, M. (2025). Autoactive CNGC15 enhances root endosymbiosis in legume and wheat. Nature. 10.1038/s41586-024-08424-7

Daniel-Mozo, M., Rombolá-Caldentey, B., Mendoza, I., Ragel, P., de, A., Carranco, R., Alcaide, A. M., Ausili, A., Schumacher, K., Albert, A., & Pardo, J. M. (2024). The vacuolar K^+^/H^+^ exchangers and calmodulin-like CML18 constitute a pH-sensing module that regulates K^+^ status in Arabidopsis. Sci Adv. 10.1126/sciadv.adp7658

DeFalco, T. A., Bender, K. W., & Snedden, W. A. (2010). Breaking the code: Ca2+ sensors in plant signalling. Biochemical Journal, 425(1), 27–40. 10.1042/BJ20091147

DeFalco, T. A., Marshall, C. B., Munro, K., Kang, H. G., Moeder, W., Ikura, M., Snedden, W. A., & Yoshioka, K. (2016). Multiple calmodulin-binding sites positively and negatively regulate arabidopsis CYCLIC NUCLEOTIDE-GATED CHANNEL12. Plant Cell, 28(7), 1738–1751. 10.1105/tpc.15.00870

Del Cerro, P., Cook, N. M., Huisman, R., Dangeville, P., Grubb, L. E., Marchal, C., Ho, A., Lam, C., & Charpentier, M. (2022). Engineered CaM2 modulates nuclear calcium oscillation and enhances legume root nodule symbiosis. Proc. Natl. Acad. Sci. U.S.A., 119 (13) e2200099119, 10.1073/pnas.2200099119

Dupont, G., & Goldbeter, A. (1998). CaM kinase II as frequency decoder of Ca2+ oscillations. BioEssays, 20(8), 607–610. 10.1002/(SICI)1521-1878(199808)20:8<607::AID-BIES2>3.0.CO;2-F

Eisner, D. A., Neher, E., Taschenberger, H., & Smith, G. (2023). Physiology of intracellular calcium buffering. Physiol Rev. 103(4):2767-2845. 10.1152/PHYSREV.00042.2022

Gall, D., Baus, E., & Dupont, G. (2000). Activation of the liver glycogen phosphorylase by Ca(2+)oscillations: a theoretical study. Journal of Theoretical Biology, 207(4), 445–454. 10.1006/jtbi.2000.2139

Gifford, J. L., Walsh, M. P., & Vogel, H. J. (2007). Structures and metal-ion-binding properties of the Ca2+-binding helix–loop–helix EF-hand motifs. Biochemical Journal, 405(2), 199–221. 10.1042/BJ20070255

Goldbeter, A., Dupont, G., & Berridge, M. J. (1990). Minimal model for signal-induced Ca2+ oscillations and for their frequency encoding through protein phosphorylation. Proc. Natl. Acad. Sci. U.S.A., 87(4), 1461–1465. 10.1073/pnas.87.4.1461

Grenzi, M., Resentini, F., Vanneste, S., Zottini, M., Bassi, A., & Costa, A. (2021). Illuminating the hidden world of calcium ions in plants with a universe of indicators. Plant Physiology, 187(2), 550–571. 10.1093/PLPHYS/KIAB339

Harper, J. R., Breton, G., & Harmon, A. (2004). Decoding Ca2+ signals through plant protein kinases. Annual Review of Plant Biology, vol. 55, 263–288. 10.1146/ANNUREV.ARPLANT.55.031903.141627

Hashimoto, K., & Kudla, J. (2011). Calcium decoding mechanisms in plants. Biochimie, 93(12), 2054–2059. 10.1016/J.BIOCHI.2011.05.019

Hepler, P. K., Vidali, L., & Cheung, A. Y. (2001). Polarized cell growth in higher plants. Annual Review of Cell and Developmental Biology, vol. 17, 159–187. 10.1146/ANNUREV.CELLBIO.17.1.159

Hill, A. V. (1910). A new mathematical treatment of changes of ionic concentration in muscle and nerve under the action of electric currents, with a theory as to their mode of excitation. The Journal of Physiology, 40(3), 190–224. 10.1113/JPHYSIOL.1910.SP001366

Ikura, M. (1996). Calcium binding and conformational response in EF-hand proteins. Trends in Biochemical Sciences, 21(1), 14–17. 10.1016/S0968-0004(06)80021-6

Ishida, H., & Vogel, H. J. (2006). Protein-Peptide Interaction Studies Demonstrate the Versatility of Calmodulin Target Protein Binding. Protein & Peptide Letters, 13(5), 455–465. 10.2174/092986606776819600

Jin, S., Zhong, X., Hu, Z., & Jiang, Z. (2025). Ca2+ flux in plant responses to abiotic stress. Journal of Plant Physiology, 315, 154648. 10.1016/J.JPLPH.2025.154648

Jurado, L. A., Chockalingam, S., & Jarrett, H. W. (1999). Apocalmodulin. Physiol Rev., 79(3):661–82. 10.1152/physrev.1999.79.3.661

Kader, M. A., & Lindberg, S. (2010). Cytosolic calcium and pH signaling in plants under salinity stress. Plant Signaling & Behavior, 5(3), 233–238. 10.4161/PSB.5.3.10740

Kaya, H., Nakajima, R., Iwano, M., Kanaoka, M. M., Kimura, S., Takeda, S., Kawarazaki, T., Senzaki, E., Hamamura, Y., Higashiyama, T., Takayama, S., Abe, M., & Kuchitsu, K. (2014). Ca2+-Activated Reactive Oxygen Species Production by Arabidopsis RbohH and RbohJ Is Essential for Proper Pollen Tube Tip Growth. The Plant Cell, 26(3), 1069–1080. 10.1105/TPC.113.120642

Keeler, C., Poon, G., Kuo, I. Y., Ehrlich, B. E., & Hodsdon, M. E. (2013). An Explicit Formulation Approach for the Analysis of Calcium Binding to EF-Hand Proteins Using Isothermal Titration Calorimetry. Biophysical Journal, 105(12), 2843–2853. 10.1016/J.BPJ.2013.11.017

Kintzer, A. F., & Stroud, R. M. (2016). Structure, inhibition and regulation of two-pore channel TPC1 from Arabidopsis thaliana. Nature, 531(7593), 258–264. 10.1038/nature17194

Klotz, I. M. (2004). Ligand-Receptor Complexes: Origin and Development of the Concept. Journal of Biological Chemistry, 279(1), 1–12. 10.1074/jbc.X300006200

Köster, P., He, G., Liu, C., Dong, Q., Hake, K., Schmitz-Thom, I., Heinkow, P., Eirich, J., Wallrad, L., Hashimoto, K., Schültke, S., Finkemeier, I., Romeis, T., & Kudla, J. (2025). A bi-kinase module sensitizes and potentiates plant immune signaling. Science Advances, 11(4). 10.1126/SCIADV.ADT9804

Krämer, U. (2025). Changing paradigms for the micronutrient zinc, a known protein cofactor, as a signal relaying also cellular redox state. Quantitative Plant Biology, 6, e7. 10.1017/QPB.2025.4

Kudla, J., Becker, D., Grill, E., Hedrich, R., Hippler, M., Kummer, U., Parniske, M., Romeis, T., & Schumacher, K. (2018). Advances and current challenges in calcium signaling. New Phytologist, vol. 218 (2), 414–431. 10.1111/nph.14966

La Verde, V., Dominici, P., & Astegno, A. (2018). Towards understanding plant calcium signaling through calmodulin-like proteins: A biochemical and structural perspective. International Journal of Molecular Sciences, vol. 19(5). 10.3390/ijms19051331

Larsen, A. Z., Olsen, L. F., & Kummer, U. (2004). On the encoding and decoding of calcium signals in hepatocytes. Biophysical Chemistry, 107(1), 83–99. 10.1016/j.bpc.2003.08.010

Latz, A., Mehlmer, N., Zapf, S., Mueller, T. D., Wurzinger, B., Pfister, B., Csaszar, E., Hedrich, R., Teige, M., & Becker, D. (2013). Salt Stress Triggers Phosphorylation of the Arabidopsis Vacuolar K+ Channel TPK1 by Calcium-Dependent Protein Kinases (CDPKs). Molecular Plant, 6(4), 1274–1289. 10.1093/MP/SSS158

Lee, H. J., & Seo, P. J. (2021). Ca2+talyzing Initial Responses to Environmental Stresses. Trends in Plant Science, 26(8), 849–870. 10.1016/J.TPLANTS.2021.02.007

Lee, J. Y., Yoo, B. C., & Harmon, A. C. (1998). Kinetic and calcium-binding properties of three calcium-dependent protein kinase isoenzymes from soybean. Biochemistry, 37(19), 6801–6809. 10.1021/BI980062Q

Liese, A., Eichstädt, B., Lederer, S., Schulz, P., Oehlschläger, J., Matschi, S., Feijó, J. A., Schulze, W. X., Konrad, K. R., & Romeis, T. (2024). Imaging of plant calcium-sensor kinase conformation monitors real time calcium-dependent decoding in planta. Plant Cell vol. 36(2), 276–297. 10.1093/plcell/koad196

Luan, S., & Wang, C. (2021). Calcium Signaling Mechanisms Across Kingdoms. Annual Review of Cell and Developmental Biology, 57(23), 23. 10.1146/annurev-cellbio-120219

Malhó, R., Moutinho, A., Van Der Luit, A., & Trewavas, A. J. (1998). Spatial characteristics to calcium signalling; the calcium wave as a basic unit in plant cell calcium signalling. Philosophical Transactions of the Royal Society of London. Series B: Biological Sciences, 353(1374), 1463–1473. 10.1098/RSTB.1998.0302

Martins, T. V., Evans, M. J., Woolfenden, H. C., & Morris, R. J. (2013). Towards the Physics of Calcium Signalling in Plants. Plants*, vol.* 2(4), 541–588. 10.3390/PLANTS2040541

Matsui, T. (2022). Calcium wave propagation during cell extrusion. Current Opinion in Cell Biology, 76, 102083. 10.1016/J.CEB.2022.102083

Mazzei, L., Ciurli, S., & Zambelli, B. (2016). Isothermal Titration Calorimetry to Characterize Enzymatic Reactions. Methods in Enzymology, 567, 215–236. 10.1016/BS.MIE.2015.07.022

McAinsh, M. R., Brownlee, C., & Hetherington, A. M. (1997). Calcium ions as second messengers in guard cell signal transduction. Physiologia Plantarum, 100(1), 16–29. 10.1111/J.1399-3054.1997.TB03451.X

Mérida-Quesada, F., Vergara-Valladares, F., Rubio-Meléndez, M. E., Hernández-Rojas, N., González-González, A., Michard, E., Navarro-Retamal, C., & Dreyer, I. (2022). TPC1-Type Channels in Physcomitrium patens: Interaction between EF-Hands and Ca2+. Plants, 11(24), 3527. 10.3390/PLANTS11243527

Ming, Y., Peng, Y., Liu, Q., Fu, D., Lin, Q., You, P., Wang, X., Zhang, X., Wang, Y., Gong, Z., Song, W., Yang, S., & Ding, Y. (2025). Coordinated control of calcium signaling by CPK3 and CaM2 via CNGCs in response to cold stress in Arabidopsis. Developmental Cell. 10.1016/J.DEVCEL.2025.06.020

Mukherjee, S., Kalra, G., & Bhatla, S. C. (2022). Trifluoperazine (TFP)-mediated fluorescence imaging approach reveals a probable calmodulin (CaM)-independent calcium signaling accompanying differential protein phosphorylation in NaCl-stressed sunflower seedlings (Helianthus annuus L. var. KBSH44). South African Journal of Botany, 150, 596–606. 10.1016/j.sajb.2022.08.008

Neamtu, A., Serban, D. N., Barritt, G. J., Isac, D. L., Vasiliu, T., Laaksonen, A., & Serban, I. L. (2023). Molecular dynamics simulations reveal the hidden EF-hand of EF-SAM as a possible key thermal sensor for STIM1 activation by temperature. Journal of Biological Chemistry, 299(8), 104970. 10.1016/J.JBC.2023.104970

Nguyen, C. T., Kurenda, A., Stolz, S., Chételat, A., & Farmer, E. E. (2018). Identification of cell populations necessary for leaf-toleaf electrical signaling in a wounded plant. Proc. Natl. Acad. Sci. U.S.A., 115(40), 10178–10183. 10.1073/PNAS.1807049115

Northrop, D. B., & Simpson, F. B. (1997). New concepts in bioorganic chemistry beyond enzyme kinetics: Direct determination of mechanisms by stopped-flow mass spectrometry. Bioorganic & Medicinal Chemistry, 5(4), 641–644. 10.1016/S0968-0896(97)00020-5

Ogasawara, Y., Kaya, H., Hiraoka, G., Yumoto, F., Kimura, S., Kadota, Y., Hishinuma, H., Senzaki, E., Yamagoe, S., Nagata, K., Nara, M., Suzuki, K., Tanokura, M., & Kuchitsu, K. (2008). Synergistic Activation of the Arabidopsis NADPH Oxidase AtrbohD by Ca2+ and Phosphorylation. Journal of Biological Chemistry, 283(14), 8885–8892. 10.1074/JBC.M708106200

Ogawa, S. T., Zhang, W., Staiger, C. J., & Kessler, S. A. (2025). MLO-mediated Ca2+ influx regulates root hair tip growth in Arabidopsis. New Phytologist, 248(6), 3024–3039. 10.1111/NPH.70378

Oh, M. H., Wu, X., Kim, H. S., Harper, J. F., Zielinski, R. E., Clouse, S. D., & Huber, S. C. (2012). CDPKs are dual-specificity protein kinases and tyrosine autophosphorylation attenuates kinase activity. FEBS Letters, 586(23), 4070–4075. 10.1016/J.FEBSLET.2012.09.040

Pan, Y., Chai, X., Gao, Q., Zhou, L., Zhang, S., Li, L., & Luan, S. (2019). Dynamic Interactions of Plant CNGC Subunits and Calmodulins Drive Oscillatory Ca 2+ Channel Activities. Developmental Cell, 48(5), 710–725.e5. 10.1016/j.devcel.2018.12.025

Park, H. Y., Kim, S. A., Korlach, J., Rhoades, E., Kwok, L. W., Zipfel, W. R., Waxham, M. N., Webb, W. W., & Pollack, L. (2008). Conformational changes of calmodulin upon Ca2+ binding studied with a microfluidic mixer. Proc. Natl. Acad. Sci. U.S.A., 105(2), 542–547. 10.1073/PNAS.0710810105

Park, S. Y., Fung, P., Nishimura, N., Jensen, D. R., Fujii, H., Zhao, Y., Lumba, S., Santiago, J., Rodrigues, A., Chow, T. F. F., Alfred, S. E., Bonetta, D., Finkelstein, R., Provart, N. J., Desveaux, D., Rodriguez, P. L., McCourt, P., Zhu, J. K., Schroeder, J. I., … Cutler, S. R. (2009). Abscisic acid inhibits type 2C protein phosphatases via the PYR/PYL family of START proteins. Science, 324(5930), 1068–1071. 10.1126/SCIENCE.1173041

Perochon, A., Aldon, D., Galaud, J. P., & Ranty, B. (2011). Calmodulin and calmodulin-like proteins in plant calcium signaling. Biochimie, 93(12), 2048–2053. 10.1016/J.BIOCHI.2011.07.012

Persechini, A., Moncrief, N. D., & Kretsinger, R. H. (1989). The EF-hand family of calcium-modulated proteins. Trends in Neurosciences, 12(11), 462–467. 10.1016/0166-2236(89)90097-0

Pirayesh, N., Giridhar, M., Ben Khedher, A., Vothknecht, U. C., & Chigri, F. (2021). Organellar calcium signaling in plants: An update. Biochimica et Biophysica Acta - Molecular Cell Research, vol. 1868(4). 10.1016/j.bbamcr.2021.118948

Praat, M., De Smet, I., & Van Zanten, M. (2021). Protein kinase and phosphatase control of plant temperature responses. Journal of Experimental Botany, 72(21), 7459–7473. 10.1093/JXB/ERAB345

Prank, K., Läer, L., Mühlen, A. von zur, Brabant, G., Schöfl, C., A., B. M. . J. and G., Schöfl C., P. K. and B. G., Berridge, M. J., Berridge, M. J., H., H. P. I. and S., R., C. R. J. and S. T., Schöfl C., R. L., L. H., M. T., V. Z. M. A. and B. G., Panten U., S. M. and S. C., Hanson P. I., M. T., S. L. and S. H., Schöfl C., B. G., H. R. D., V. Z. M. A., C. P. H. and C. K. S. R., Chay T. R., L. Y. S. and F. Y. S., Iida Y., S. T., M. Y., O. K., M. J. I., S. H., N. Y., H. H. and N. I., Krueger K. A., B. H., L. M. and E. R. A., Hisatomi M., H. H. and N. I., … Prank K., S. C., L. L., W. M., V. Z. M. A., B. G. and G. F. (1998). Decoding of intracellular calcium spike trains. Europhysics Letters (EPL), 42(2), 143–148. 10.1209/epl/i1998-00220-2

Project, E., Friedman, R., Nachliel, E., & Gutman, M. (2006). A Molecular Dynamics Study of the Effect of Ca2+ Removal on Calmodulin Structure. Biophysical Journal, 90(11), 3842–3850. 10.1529/BIOPHYSJ.105.077792

Quinn, C. F., Carpenter, M. C., Croteau, M. L., & Wilcox, D. E. (2016). Isothermal Titration Calorimetry Measurements of Metal Ions Binding to Proteins. Methods in Enzymology, 567, 3–21. 10.1016/BS.MIE.2015.08.021

Ranty, B., Aldon, D., & Galaud, J. P. (2006). Plant Calmodulins and Calmodulin-Related Proteins. Plant Signaling & Behavior, 1(3), 96–104. 10.4161/PSB.1.3.2998

Resentini, F., Grenzi, M., Ancora, D., Cademartori, M., Luoni, L., Franco, M., Bassi, A., Bonza, M. C., & Costa, A. (2021). Simultaneous imaging of ER and cytosolic Ca2+ dynamics reveals long-distance ER Ca2+ waves in plants. Plant Physiology, 187(2), 603–617. 10.1093/PLPHYS/KIAB251

Sánchez-Barrena, M. J., Martínez-Ripoll, M., Zhu, J. K., & Albert, A. (2005). The Structure of the Arabidopsis thaliana SOS3: Molecular Mechanism of Sensing Calcium for Salt Stress Response. Journal of Molecular Biology, 345(5), 1253–1264. 10.1016/J.JMB.2004.11.025

Schuster, S., Knoke, B., & Marhl, M. (2005). Differential regulation of proteins by bursting calcium oscillations--a theoretical study. Bio Systems, 81(1), 49–63. 10.1016/j.biosystems.2005.02.004

Senadheera, P., & Maathuis, F. J. M. (2009). Differentially regulated kinases and phosphatases in roots may contribute to inter-cultivar difference in rice salinity tolerance. Plant Signaling & Behavior, 4(12), 2553–2563. 10.4161/PSB.4.12.9969

Shabala, S. N., & Newman, I. A. (1997). Proton and calcium flux oscillations in the elongation region correlate with root nutation. Physiologia Plantarum, 100(4), 917–926. 10.1111/J.1399-3054.1997.TB00018.X

Swainsbury, D. J. K., Zhou, L., Oldroyd, G. E. D., & Bornemann, S. (2012). Calcium ion binding properties of medicago truncatula calcium/calmodulin-dependent protein kinase. Biochemistry, 51(35), 6897–6907. 10.1021/BI300826M

Tang, R. J., Wang, C., Li, K., & Luan, S. (2020). The CBL–CIPK Calcium Signaling Network: Unified Paradigm from 20 Years of Discoveries. Trends in Plant Science, 25(6), 604–617. 10.1016/J.TPLANTS.2020.01.009

Teleman, A., Drakenberg, T., & Forsén, S. (1986). Kinetics of Ca2+ binding to calmodulin and its tryptic fragments studied by 43Ca-NMR. Biochimica et Biophysica Acta (BBA) - Protein Structure and Molecular Enzymology, 873(2), 204–213. 10.1016/0167-4838(86)90047-6

Tian, W., Wang, C., Gao, Q., Li, L., & Luan, S. (2020). Calcium spikes, waves and oscillations in plant development and biotic interactions. Nature Plants, 6(7), 750–759. 10.1038/s41477-020-0667-6

Toyota, M., Spencer, D., Sawai-Toyota, S., Jiaqi, W., Zhang, T., Koo, A. J., Howe, G. A., & Gilroy, S. (2018). Glutamate triggers long-distance, calcium-based plant defense signaling. Science, 361(6407), 1112–1115. 10.1126/SCIENCE.AAT7744

Troilo, F., Pedretti, M., Travaglini-Allocatelli, C., Astegno, A., & Di Matteo, A. (2022). Rapid kinetics of calcium dissociation from plant calmodulin and calmodulin-like proteins and effect of target peptides. Biochemical and Biophysical Research Communications, 590, 103–108. 10.1016/J.BBRC.2021.12.077

Tso, S. C., Jowitt, T. A., & Brautigam, C. A. (2022). The feasibility of determining kinetic constants from isothermal titration calorimetry data. Biophysical Journal, 121(12), 2474–2484. 10.1016/J.BPJ.2022.04.035

Weinl, S., & Kudla, J. (2009). The CBL–CIPK Ca2+-decoding signaling network: function and perspectives. New Phytologist, 184(3), 517–528. 10.1111/J.1469-8137.2009.02938.X

Williams, R. J. P. (2002). Calcium. Methods in Molecular Biology, 172, 21–49. 10.1385/1-59259-183-3:021

Zauser, M., Liese, A., Feldman-Salit, A., Krebs, M., Schumacher, K., Romeis, T., Kummer, U., & Pahle, J. (2025). Differential regulation of calcium-activated plant kinases in Arabidopsis thaliana. Plant Journal, 123(5), e70413. 10.1111/TPJ.70413

