## Supplementary Material for "Universal modules for decoding amplitude and frequency of Ca^2+^ signals in plants"

#### **Notes S1** $Ca^{2+}$-dependence of the $Ca^{2+}$-decoding modules 1-6.

Six different modules determine the $Ca^{2+}$-dependence of the target protein. This dependency is derived in the following and the influence of the various parameters is illustrated. We consider 4 different cases: (0) the direct binding of $Ca^{2+}$ to the TP with an (almost) immediate effect; (i) the rapid direct binding of $Ca^{2+}$ to the TP with a slower secondary modulatory effect; (ii) the rapid modulation of a calcium binding protein (CBP) by $Ca^{2+}$ and the subsequent catalytic action of the CBP on the TP; (iii) the rapid modulation of a calcium binding protein (CBP) by $Ca^{2+}$ and the subsequent binding of the CBP to the TP.

##### **(0) Direct binding of $Ca^{2+}$ to the TP that causes an (almost) immediate effect**

The simplest scenario describes the direct binding of $Ca^{2+}$ to the target protein, *e.g.* to EF hands. Compared to other cellular processes, the direct interaction of $Ca^{2+}$ ions with a target protein is so fast that it can be considered to be almost instantaneous and always in steady state. In this condition, the binding of the ligand $Ca^{2+}$ to a receptor protein can be described with the theoretical framework developed for ligand-receptor complexes, which confirms the experimentally observed sigmoidal curves. To substantially reduce the number of free parameters, the sigmoidal dependency can be satisfactorily approximated by the empirical Hill equation.

Modules 1 & 2:

The first two modules describe the cases, in which (Module 1) a target protein is activated by calcium or (Module 2) a target protein is inactivated by calcium. In order to consider the most general case, the TP also can remain residual (in)activity at zero calcium and at saturating $Ca^{2+}$. In such cases, the $Ca^{2+}$-dependence of the fraction of active target proteins can be described mathematically by:

$$TP^*_{modules1\&2} = \frac{\beta \cdot K^n + \gamma \cdot \left[Ca^{2+}\right]^n}{K^n + \left[Ca^{2+}\right]^n} \qquad \text{Eqn S1}$$

with $\beta < \gamma$ for module 1 and $\beta > \gamma$ for module 2. Here, *K* denotes the apparent dissociation constant, *n* the Hill-coefficient, $0 \leq \beta \leq 1$ the fraction of active target proteins in the absence of $Ca^{2+}$, and $0 \leq \gamma \leq 1$ the fraction of active target proteins under saturating $Ca^{2+}$. $TP^*_{modules1\&2}$ takes values between $\beta$ (0 $Ca^{2+}$) and $\gamma$ (high $Ca^{2+}$). For module 1 ($\beta < \gamma$) it increases with increasing [$Ca^{2+}$], while for module 2 ($\beta > \gamma$) it decreases with increasing [$Ca^{2+}$]. The parameter *K* determines the $Ca^{2+}$ concentration at which the curve has its midpoint ($TP^*_{module1} = (\gamma + \beta)/2$), while the parameter *n* determines the steepness of the curve. **Fig. S1** illustrates the dependence on the different parameters.

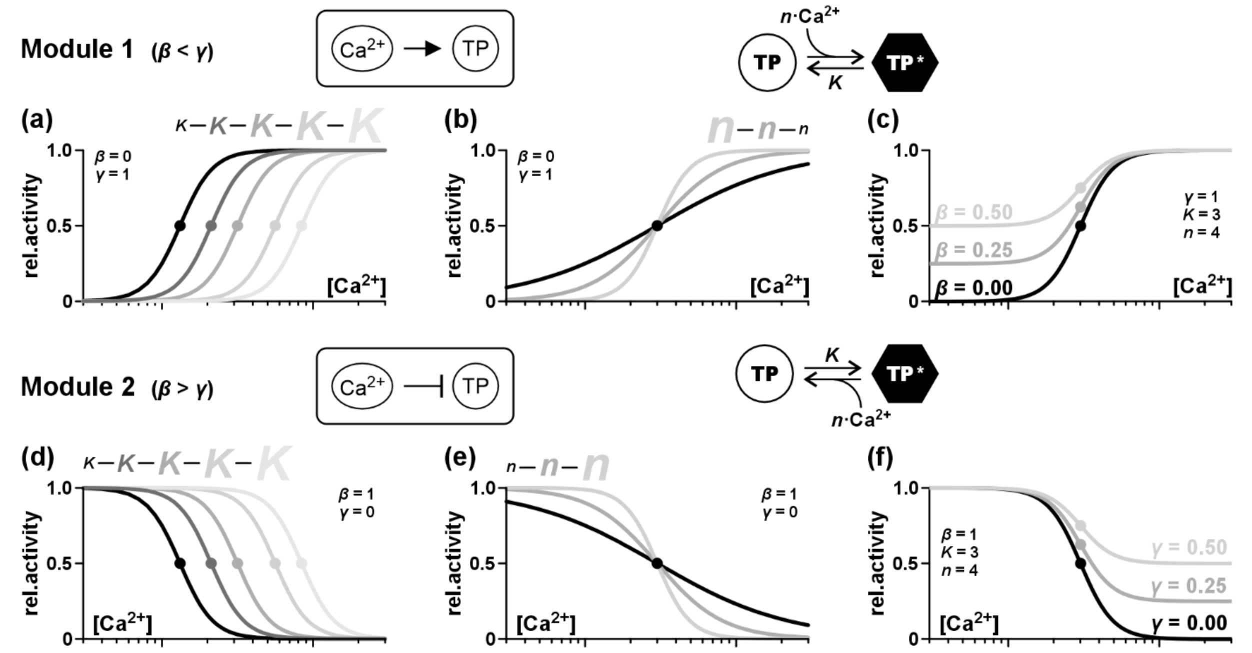

**Fig. S1** $Ca^{2+}$-dependent activity of the target protein in module 1 ($\beta < \gamma$, a-c) and module 2 ($\beta > \gamma$, d-f) illustrated for different parameters (a, d) *K*, (b, e) *n*, (c) *β*, and (f) *γ*. (a-c) Module 1. Here, $\gamma = 1$, *i.e.* the TP-activity at high $Ca^{2+}$ is 1. (d-f) Module 2. Here, $\beta = 1$, *i.e.* the TP-activity at low $Ca^{2+}$ is 1. In both modules, the $Ca^{2+}$-sensitivity decreases with increasing *K*, and with increasing *n* the steepness of the curve increases. With increasing *β* the activity at very low [$Ca^{2+}$] increases, while with increasing *γ* the activity at very high [$Ca^{2+}$] increases. The dots indicate the midpoints of the respective curves.

**(i) Rapid direct binding of $Ca^{2+}$ to the TP with a slower subsequent modulatory effect**

The first more complex scenario is a time-dependent version of the scenario described before. Also here, $Ca^{2+}$ binds directly to the target protein, *e.g.* to EF hands. This time, however, the activity change of the TP does not occur instantaneously but with a significant delay. The underlying changes in the protein conformation proceed on a much slower time scale. There are four different cases to be considered: (Module i.3) the inactive TP binds $Ca^{2+}$ and then undergoes a conformational change that activates it; (Module i.4) the inactive TP can undergo an activating conformational change in the absence of $Ca^{2+}$ but $Ca^{2+}$-binding inhibits this modification; (Module i.5) The active TP binds $Ca^{2+}$ and then undergoes a conformational change that inactivates it; (Module i.6) The active TP can undergo a conformational change that inactivates it, but binding of $Ca^{2+}$ prevents inactivation.

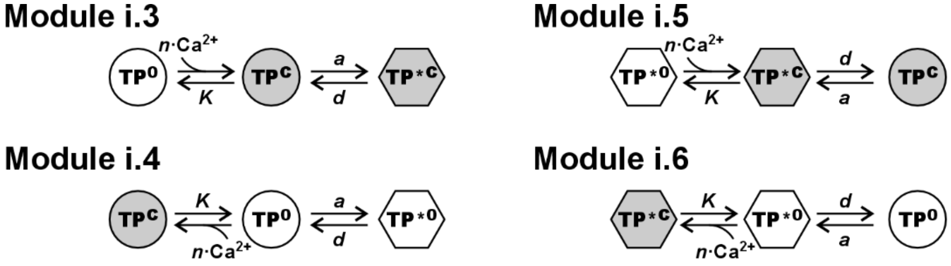


The variation of the fraction of activated target proteins (*TP**) in time is determined by the following differential equation systems

$$\left\{\begin{aligned}\frac{d}{dt}TP^{*C}(t) &= -d \cdot TP^{*C}(t) + a \cdot TP^{C}(t)\\ \frac{d}{dt}TP^{C}(t) &= d \cdot TP^{*C}(t) - (a+D) \cdot TP^{C}(t) + A \cdot [1 - TP^{*C}(t) - TP^{C}(t)]\end{aligned}\right\} \quad TP^{*}_{i.3}(t) = TP^{*C}(t) \qquad \text{Eqn S2}$$

$$\left\{\begin{aligned}\frac{d}{dt}TP^{*0}(t) &= -d \cdot TP^{*0}(t) + a \cdot [1 - TP^{*0}(t) - TP^{C}(t)]\\ \frac{d}{dt}TP^{C}(t) &= -D \cdot TP^{C}(t) + A \cdot [1 - TP^{*0}(t) - TP^{C}(t)]\end{aligned}\right\} \quad TP^{*}_{i.4}(t) = TP^{*0}(t) \qquad \text{Eqn S3}$$

$$\left\{\begin{aligned}\frac{d}{dt}TP^{C}(t) &= -a \cdot TP^{C}(t) + d \cdot TP^{*C}(t)\\ \frac{d}{dt}TP^{*C}(t) &= a \cdot TP^{C}(t) - (d+D) \cdot TP^{*C}(t) + A \cdot [1 - TP^{*C}(t) - TP^{C}(t)]\end{aligned}\right\} \quad TP^{*}_{i.5}(t) = 1 - TP^{C}(t) \qquad \text{Eqn S4}$$

$$\left\{\begin{aligned}\frac{d}{dt}TP^{0}(t) &= -a \cdot TP^{0}(t) + d \cdot TP^{*0}(t)\\ \frac{d}{dt}TP^{*0}(t) &= a \cdot TP^{0}(t) - (d+A) \cdot TP^{*0}(t) + D \cdot [1 - TP^{*0}(t) - TP^{0}(t)]\end{aligned}\right\} \quad TP^{*}_{i.6}(t) = 1 - TP^{0}(t) \qquad \text{Eqn S5}$$

with $[A, D] \gg [a, d]$, ($A = n\text{Ca}^{2+}$, $D = K$), *i.e.* the $Ca^{2+}$ binding is much faster than the conformational change (*a*, *d*). For a complex time-dependent $Ca^{2+}$ signal, these differential equation systems can only be solved numerically. Nevertheless, a transient $Ca^{2+}$ signal can be interpreted in a staircase manner as a sequence of short $Ca^{2+}$ pulses with constant amplitude. For such time intervals, the differential equation systems have constant coefficients and can be solved analytically: $TP^*(t) = TP^*_{ss} + C_1 \cdot e^{-t/\tau_1} + C_2 \cdot e^{-t/\tau_2}$, with coefficients $C_1$ and $C_2$ that depend on the situation at $t = 0$ and the time constants

$$\tau_{1,2} = 2 \cdot \left(A + D + a + d \pm (A + D - a - d) \cdot \sqrt{1 + \frac{4xY}{(A+D-a-d)^2}}\right)^{-\frac{1}{2}} \qquad \text{Eqn S6}$$

Here, $xY = aD$ for Module i.3, $xY = aA$ for Module i.4, $xY = dD$ for Module i.5, and $xY = dA$ for Module i.6. Considering that $A + D \gg a + d$, it can be approximated $\sqrt{1 + \frac{4xY}{(A+D-a-d)^2}} \approx 1 + \frac{2xY}{(A+D-a-d)^2}$, so that $\tau_1 \approx 1/(a + d - xY/(A + D))$ and $\tau_2 \approx 1/(A + D)$. The second time constant is much smaller than the first one and represents the (almost) instantaneous calcium binding reaction. Therefore, for physiologically relevant time intervals the solution of the equation systems can be written as

$$TP^*(t) = TP^*_{ss} + (TP^*_0 - TP^*_{ss}) \cdot e^{-t/\tau} \qquad \text{Eqn S7}$$

Here $TP^*_0$ is the *TP** value at the beginning of the $Ca^{2+}$ pulse and $TP^*_{ss}$ the value in steady state equilibrium. While $TP^*_0$ depends on the history of the system, $TP^*_{ss}$ and the time constant $\tau$ ($= \tau_1$) are characteristic parameters of the respective module. In analogy to eqn. (S1), the steady state of the fast $Ca^{2+}$-binding reaction can be described in generalized form as $\{\beta \cdot K^n + \gamma \cdot [\mathrm{Ca}^{2+}]^n\}/\{K^n + [\mathrm{Ca}^{2+}]^n\}$ with $0 \le \beta, \gamma \le 1$ to also include residual activities of the $Ca^{2+}$-driven reaction at zero $Ca^{2+}$ ($\beta$) and at very high $Ca^{2+}$ ($\gamma$). We could describe both $A/(A + D)$ and $D/(A + D)$ by the same general function, but it needs to be kept in mind that $A/(A + D)$ implies $\beta < \gamma$ (module 1) and $D/(A + D)$ implies $\beta > \gamma$ (module 2).

Module i.3

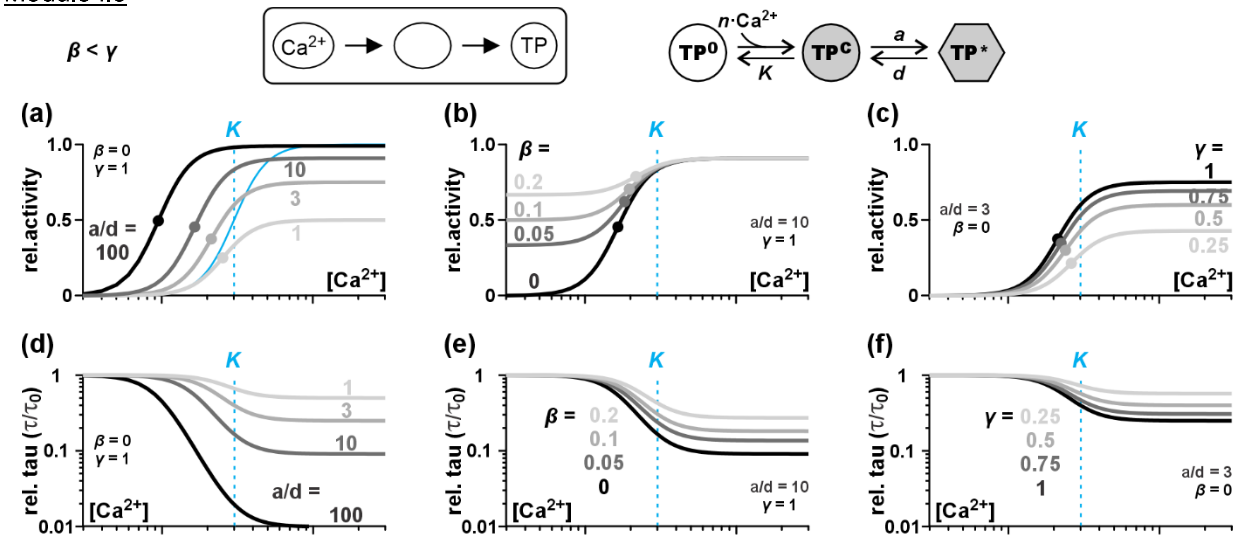


**Fig. S2** Effect of the parameters *a/d*, *β*, and *γ* on the $Ca^{2+}$-dependence of module i.3. The parameters *K*=3 and *n*=4 were fixed. (a-c) $Ca^{2+}$-dependent steady state activity of the target protein in module i.3 illustrated for different parameters (a) *a/d*, (b) *β* (=activity at low $Ca^{2+}$), and *γ* (=activity at high $Ca^{2+}$). The solid blue curve shows the $Ca^{2+}$-dependence of the $Ca^{2+}$-binding reaction for *β*=0/*γ*=1 and the dashed blue lines indicate the midpoint (*K*) of this curve. The midpoints of the $TP_{ss}$ curves are indicated as dots and grey-scale coded with the curves and the indicated values for the parameters. Please note that with increasing *a/d*-ratio the midpoint of the $TP_{ss}$ curve shifts to lower $Ca^{2+}$-values, i.e. the sensitivity of the TP to $Ca^{2+}$ increases. Panel (b) shows the dependency on *β* for *a/d*=10 and *γ*=1, while panel (c) illustrates the dependency on *γ* for *a/d*=3 and *β*=0. An increase in *β* and a decrease in *γ* reduce the $Ca^{2+}$ sensitivity of the TP, *i.e.* the midpoint shifts towards the *K* value of the original $Ca^{2+}$-binding reaction. (d-f) $Ca^{2+}$-dependence of the time constant $\tau$ displayed relative to the $\tau$-value at zero $Ca^{2+}$, $\tau_0$. Please note the logarithmic scale of the y-axes. (d) With increasing *a/d*-ratio the time constant of the module becomes smaller, i.e. the module reacts faster at higher $Ca^{2+}$. (e-f) An increase in the basal activity at zero $Ca^{2+}$ (parameter *β*) or a decrease in the activity at high $Ca^{2+}$ (parameter *γ*) reduce the difference between the time constants at high and at low $Ca^{2+}$.

$$TP^*_{ss,i.3} = \frac{a}{a + d \cdot \frac{A+D}{A}} = \frac{\frac{\beta \cdot a/d}{1+\beta \cdot a/d} \cdot \left(\frac{1+\beta \cdot a/d}{1+\gamma \cdot a/d}\right) \cdot K^n + \frac{\gamma \cdot a/d}{1+\gamma \cdot a/d}\left[\mathrm{Ca}^{2+}\right]^n}{\left(\frac{1+\beta \cdot a/d}{1+\gamma \cdot a/d}\right) \cdot K^n + \left[\mathrm{Ca}^{2+}\right]^n} \qquad \text{Eqn S8}$$

$$\tau_{i.3} = \frac{1}{d+a\cdot\frac{A}{A+D}} = \frac{1}{d+\beta\cdot a}\cdot\left(1-\frac{(\gamma-\beta)\cdot a/d}{1+\gamma\cdot a/d}\cdot\frac{\left[\mathrm{Ca}^{2+}\right]^n}{\left(\frac{1+\beta\cdot a/d}{1+\gamma\cdot a/d}\right)\cdot K^n+\left[\mathrm{Ca}^{2+}\right]^n}\right) \qquad \text{Eqn S9}$$

$TP^*_{ss,i.3}$ increases with increasing [$Ca^{2+}$]. The midpoint $Ca^{2+}$-concentration of the curve is smaller than *K*, the midpoint $Ca^{2+}$-concentration of the calcium-binding reaction, $\left[\mathrm{Ca}^{2+}\right]_{mid,i.3} \leq K$. Thus, with the additional step of a conformational change the TP gains $Ca^{2+}$ sensitivity. The time constant of module i.3, $\tau_{i.3}$, decreases with increasing [$Ca^{2+}$]. The midpoint of the curve is identical to the midpoint of the $TP^*_{ss,i.3}$ curve and the slope is determined by the parameter *n*. **Fig. S2** illustrates the influence of the parameters $\beta$, $\gamma$ and $a/d$ on the $Ca^{2+}$-dependence of the curves $TP^*_{ss,i.3}$ and $\tau_{i.3}$ for fixed *K* and *n*.

Module i.4

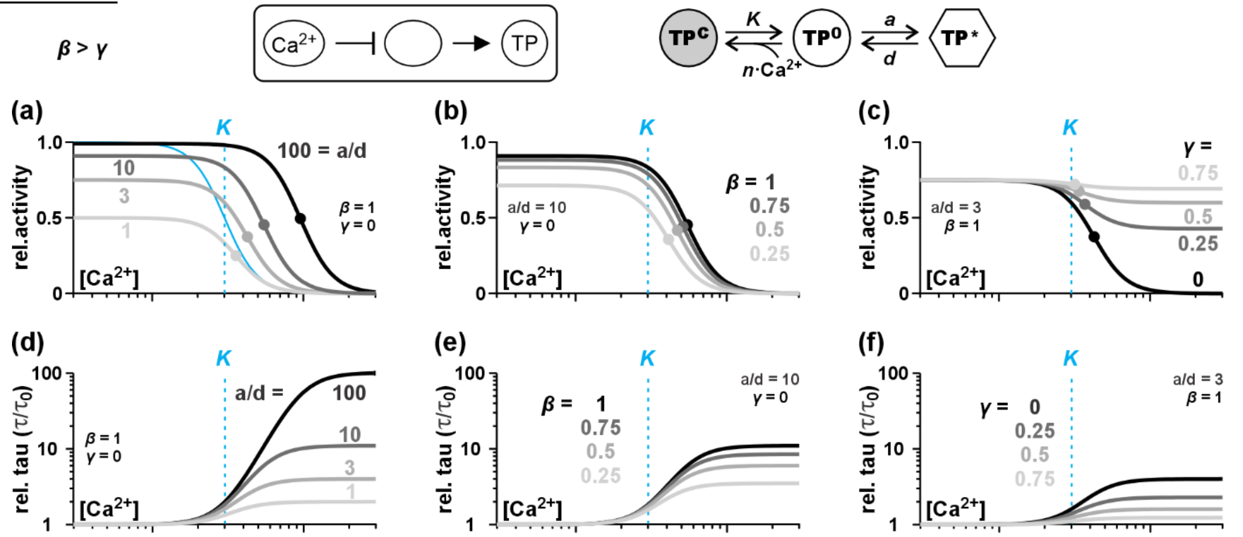


**Fig. S3** Effect of the parameters *a/d*, *β*, and *γ* on the $Ca^{2+}$-dependence of module i.4. The parameters *K*=3 and *n*=4 were fixed. (a-c) $Ca^{2+}$-dependent steady state activity of the target protein in module i.4 illustrated for different parameters (a) *a/d*, (b) *β* (=activity at low $Ca^{2+}$), and *γ* (=activity at high $Ca^{2+}$). The solid blue curve shows the $Ca^{2+}$-dependence of the $Ca^{2+}$-binding reaction for *β*=1/*γ*=0 and the dashed blue lines indicate the midpoint (*K*) of this curve. The midpoints of the $TP_{ss}$ curves are indicated as dots and grey-scale coded with the curves and the indicated values for the parameters. Please note that with increasing *a/d*-ratio the midpoint of the $TP_{ss}$ curve shifts to higher $Ca^{2+}$-values, i.e. the sensitivity of the TP to $Ca^{2+}$ decreases. Panel (b) shows the dependency on *β* for *a/d*=10 and *γ*=0, while panel (c) illustrates the dependency on *γ* for *a/d*=3 and *β*=1. A decrease in *β* and an increase in *γ* increase the $Ca^{2+}$ sensitivity of the TP, *i.e.* the midpoint shifts towards the *K* value of the original $Ca^{2+}$-binding reaction. (d-f) $Ca^{2+}$-dependence of the time constant $\tau$ displayed relative to the $\tau$-value at zero $Ca^{2+}$, $\tau_0$. Please note the logarithmic scale of the y-axes. (d) With increasing *a/d*-ratio the time constant of the module increases, i.e. the module reacts more slowly at higher $Ca^{2+}$. (e-f) A decrease in the activity at zero $Ca^{2+}$ (parameter *β*) or a increase in the activity at high $Ca^{2+}$ (parameter *γ*) reduce the difference between the time constants at high and at low $Ca^{2+}$.

$$TP^*_{ss,i.4} = \frac{a}{a+d\cdot\frac{A+D}{D}} = \frac{\frac{\beta\cdot a/d}{1+\beta\cdot a/d}\cdot\left(\frac{1+\beta\cdot a/d}{1+\gamma\cdot a/d}\right)\cdot K^n+\frac{\gamma\cdot a/d}{1+\gamma\cdot a/d}\left[\mathrm{Ca}^{2+}\right]^n}{\left(\frac{1+\beta\cdot a/d}{1+\gamma\cdot a/d}\right)\cdot K^n+\left[\mathrm{Ca}^{2+}\right]^n} \qquad \text{Eqn S10}$$

$$\tau_{i.4} = \frac{1}{d+a\cdot\frac{D}{A+D}} = \frac{1}{d+\beta\cdot a}\cdot\left(1+\frac{(\beta-\gamma)\cdot a/d}{1+\gamma\cdot a/d}\cdot\frac{\left[\mathrm{Ca}^{2+}\right]^n}{\left(\frac{1+\beta\cdot a/d}{1+\gamma\cdot a/d}\right)\cdot K^n+\left[\mathrm{Ca}^{2+}\right]^n}\right) \qquad \text{Eqn S11}$$

$TP^*_{ss,i.4}$ decreases with increasing [$Ca^{2+}$]. The midpoint $Ca^{2+}$-concentration of the curve is larger than *K*, the midpoint $Ca^{2+}$-concentration of the calcium-binding reaction, $\left[\mathrm{Ca}^{2+}\right]_{mid,i4} \geq K$. Thus, with the additional step of a conformational change the TP loses $Ca^{2+}$ sensitivity. The time constant of module i.4, $\tau_{i.4}$, increases with increasing [$Ca^{2+}$]. The midpoint of the curve is identical to the midpoint of the $TP^*_{ss,i.4}$ curve and the slope is determined by the parameter *n*. **Fig. S3** illustrates the influence of the parameters $\beta$, $\gamma$ and $a/d$ on the $Ca^{2+}$-dependence of the curves $TP^*_{ss,i.4}$ and $\tau_{i.4}$ for fixed *K* and *n*.

Module i.5

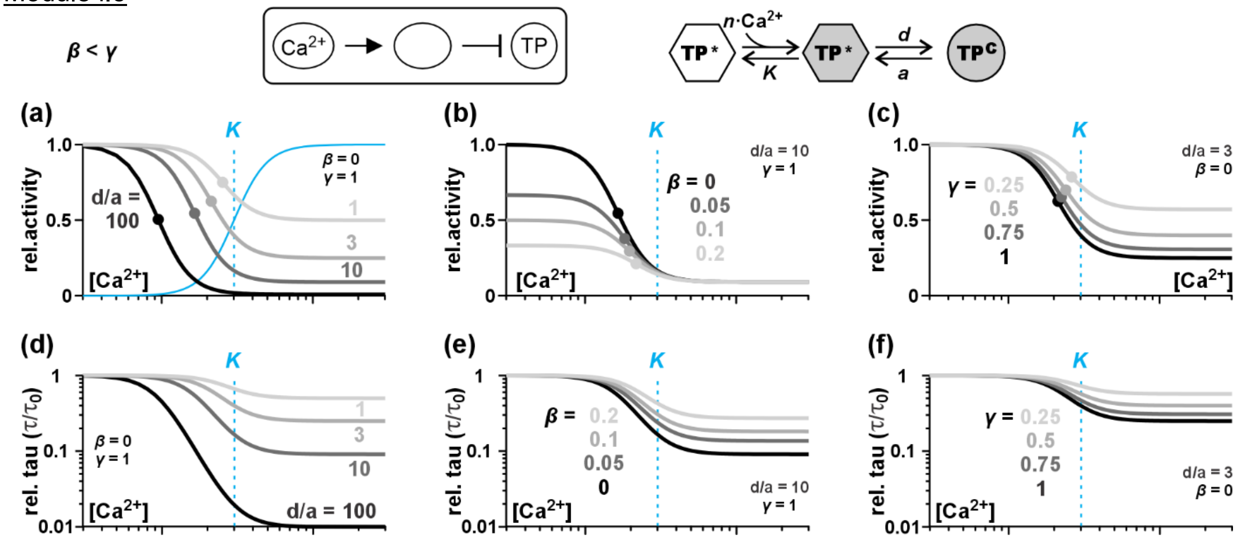


**Fig. S4** Effect of the parameters *d/a*, *β*, and *γ* on the $Ca^{2+}$-dependence of module i.5. The parameters *K*=3 and *n*=4 were fixed. (a-c) $Ca^{2+}$-dependent steady state activity of the target protein in module i.5 illustrated for different parameters (a) *d/a*, (b) *β* (=activity at low $Ca^{2+}$), and *γ* (=activity at high $Ca^{2+}$). The solid blue curve shows the $Ca^{2+}$-dependence of the $Ca^{2+}$-binding reaction for *β*=0/*γ*=1 and the dashed blue lines indicate the midpoint (*K*) of this curve. The midpoints of the $TP_{ss}$ curves are indicated as dots and grey-scale coded with the curves and the indicated values for the parameters. Please note that with increasing *d/a*-ratio the midpoint of the $TP_{ss}$ curve shifts to lower $Ca^{2+}$-values, i.e. the sensitivity of the TP to $Ca^{2+}$ increases. Panel (b) shows the dependency on *β* for *d/a*=10 and *γ*=1, while panel (c) illustrates the dependency on *γ* for *d/a*=3 and *β*=0. An increase in *β* and a decrease in *γ* reduce the $Ca^{2+}$ sensitivity of the TP, *i.e.* the midpoint shifts towards the *K* value of the original $Ca^{2+}$-binding reaction. (d-f) $Ca^{2+}$-dependence of the time constant $\tau$ displayed relative to the $\tau$-value at zero $Ca^{2+}$, $\tau_0$. Please note the logarithmic scale of the y-axes. (d) With increasing *d/a*-ratio the time constant of the module decreases, i.e. the module reacts faster at higher $Ca^{2+}$. (e-f) An increase in the basal activity (parameter *β*) or a decrease in the activity at high $Ca^{2+}$ (parameter *γ*) reduce the difference between the time constants at high and at low $Ca^{2+}$.

$$TP^*_{ss,i.5} = 1 - \frac{d}{d + a \cdot \frac{A+D}{A}} = \frac{\frac{1}{1+\beta\cdot d/a}\cdot\left(\frac{1+\beta\cdot d/a}{1+\gamma\cdot d/a}\right)\cdot K^n + \frac{1}{1+\gamma\cdot d/a}\cdot\left[\mathrm{Ca}^{2+}\right]^n}{\left(\frac{1+\beta\cdot d/a}{1+\gamma\cdot d/a}\right)\cdot K^n + \left[\mathrm{Ca}^{2+}\right]^n} \qquad \text{Eqn S12}$$

$$\tau_{i.5} = \frac{1}{a + d\cdot\frac{A}{A+D}} = \frac{1}{a+\beta\cdot d}\cdot\left(1 - \frac{(\gamma-\beta)\cdot d/a}{1+\gamma\cdot d/a}\cdot\frac{\left[\mathrm{Ca}^{2+}\right]^n}{\left(\frac{1+\beta\cdot d/a}{1+\gamma\cdot d/a}\right)\cdot K^n + \left[\mathrm{Ca}^{2+}\right]^n}\right) \qquad \text{Eqn S13}$$

$TP^*_{ss,i.5}$ decreases with increasing [$Ca^{2+}$]. The midpoint $Ca^{2+}$-concentration of the curve is smaller than *K*, the midpoint $Ca^{2+}$-concentration of the calcium-binding reaction, $\left[\mathrm{Ca}^{2+}\right]_{mid,i.5} \leq K$. Thus, with the additional step of a conformational change the TP gains $Ca^{2+}$ sensitivity. The time constant of module i.5, $\tau_{i.5}$, decreases with increasing [$Ca^{2+}$]. The midpoint of the curve is identical to the midpoint of the $TP^*_{ss,i.5}$ curve and the slope is determined by the parameter *n*. **Fig. S4** illustrates the influence of the parameters $\beta$, $\gamma$ and $d/a$ on the $Ca^{2+}$-dependence of the curves $TP^*_{ss,i.5}$ and $\tau_{i.5}$ for fixed *K* and *n*.

Module i.6

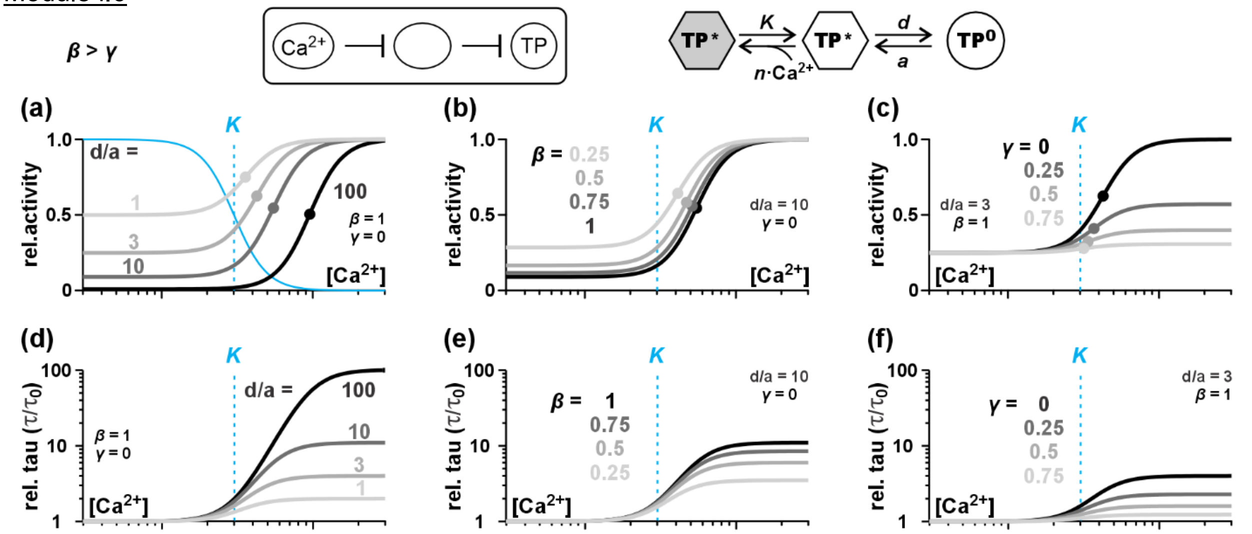


**Fig. S5** Effect of the parameters *d/a*, *β*, and *γ* on the $Ca^{2+}$-dependence of module i.6. The parameters *K*=3 and *n*=4 were fixed. (a-c) $Ca^{2+}$-dependent steady state activity of the target protein in module i.6 illustrated for different parameters (a) *d/a*, (b) *β* (=activity at low $Ca^{2+}$), and *γ* (=activity at high $Ca^{2+}$). The solid blue curve shows the $Ca^{2+}$-dependence of the $Ca^{2+}$-binding reaction for *β*=1/*γ*=0 and the dashed blue lines indicate the midpoint (*K*) of this curve. The midpoints of the $TP_{ss}$ curves are indicated as dots and grey-scale coded with the curves and the indicated values for the parameters. Please note that with increasing *d/a*-ratio the midpoint of the $TP_{ss}$ curve shifts to higher $Ca^{2+}$-values, i.e. the sensitivity of the TP to $Ca^{2+}$ decreases. Panel (b) shows the dependency on *β* for *d/a*=10 and *γ*=0, while panel (c) illustrates the dependency on *γ* for *d/a*=3 and *β*=1. A decrease in *β* and an increase in *γ* increase the $Ca^{2+}$ sensitivity of the TP, *i.e.* the midpoint shifts towards the *K* value of the original $Ca^{2+}$-binding reaction. (d-f) $Ca^{2+}$-dependence of the time constant $\tau$ displayed relative to the $\tau$-value at zero $Ca^{2+}$, $\tau_0$. Please note the logarithmic scale of the y-axes. (d) With increasing *d/a*-ratio the time constant of the module increases, i.e. the module reacts more slowly at higher $Ca^{2+}$. (e-f) A decrease in the activity at zero $Ca^{2+}$ (parameter *β*) or an increase in the activity at high $Ca^{2+}$ (parameter *γ*) reduce the difference between the time constants at high and at low $Ca^{2+}$.

$$TP^*_{ss,i.6} = 1 - \frac{d}{d + a\cdot\frac{A+D}{D}} = \frac{\frac{1}{1+\beta\cdot d/a}\cdot\left(\frac{1+\beta\cdot d/a}{1+\gamma\cdot d/a}\right)\cdot K^n + \frac{1}{1+\gamma\cdot d/a}\cdot\left[\mathrm{Ca}^{2+}\right]^n}{\left(\frac{1+\beta\cdot d/a}{1+\gamma\cdot d/a}\right)\cdot K^n + \left[\mathrm{Ca}^{2+}\right]^n} \qquad \text{Eqn S14}$$

$$\tau_{i.6} = \frac{1}{a + d\cdot\frac{D}{A+D}} = \frac{1}{a+\beta\cdot d}\cdot\left(1 + \frac{(\beta-\gamma)\cdot d/a}{1+\gamma\cdot d/a}\cdot\frac{\left[\mathrm{Ca}^{2+}\right]^n}{\left(\frac{1+\beta\cdot d/a}{1+\gamma\cdot d/a}\right)\cdot K^n+\left[\mathrm{Ca}^{2+}\right]^n}\right) \qquad \text{Eqn S15}$$

$TP^*_{ss,i.6}$ increases with increasing [$Ca^{2+}$]. The midpoint $Ca^{2+}$-concentration of the curve is larger than *K*, the midpoint $Ca^{2+}$-concentration of the calcium-binding reaction, $\left[\mathrm{Ca}^{2+}\right]_{mid,i.6} \geq K$. Thus, with the additional step of a conformational change the TP loses $Ca^{2+}$ sensitivity. The time constant of module i.6, $\tau_{i.6}$, increases with increasing [$Ca^{2+}$]. The midpoint of the curve is identical to the midpoint of the $TP^*_{ss,i.6}$ curve and the slope is determined by the parameter *n*. **Fig. S5** illustrates the influence of the parameters $\beta$, $\gamma$ and $d/a$ on the $Ca^{2+}$-dependence of the curves $TP^*_{ss,i.6}$ and $\tau_{i.6}$ for fixed *K* and *n*.

For all four modules i.3-i.6, $TP^*_{ss}$ and $\tau$ could be written as $TP^*_{ss} = \frac{b\cdot k^n + c\cdot\left[\mathrm{Ca}^{2+}\right]^n}{k^n + \left[\mathrm{Ca}^{2+}\right]^n}$ and $\tau = \tau_0\left(1 + \Delta\tau\cdot\frac{\left[\mathrm{Ca}^{2+}\right]^n}{k^n+\left[\mathrm{Ca}^{2+}\right]^n}\right)$, with the six physiologically significant parameters *n*, *k*, *b*, *c*, $\tau_0$, and $\Delta\tau$. These six parameters are unequivocally determined by the six mechanistic parameters *n*, *K*, *β*, *γ*, *a*, and *d* (**Tab. S1**). Thus, by adjusting the mechanistic parameters a cell can set the physiological parameters.

**Table S1**. Conversion between the parameter sets $(n, K, \beta, \gamma, a, d)$ and $(n, k, b, c, \tau_0, \Delta\tau)$ for the case of rapid direct binding of $Ca^{2+}$ to the TP with a slower subsequent modulatory effect.

| | | | |
|---|---|---|---|
| $TP^*(t) = TP_{ss}^* + (TP_0^* - TP_{ss}^*) \cdot e^{-t/\tau}$ | | | |
| $TP_{ss}^* = \frac{b \cdot k^n + c \cdot [Ca^{2+}]^n}{k^n + [Ca^{2+}]^n}$ $\quad \tau = \tau_0 \cdot \left(1 + \Delta\tau \cdot \frac{[Ca^{2+}]^n}{k^n + [Ca^{2+}]^n}\right)$ | | | |
| $\boldsymbol{\beta < \gamma}$ | **Module 1** 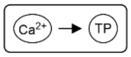  | **Module i.3** 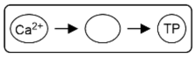  | **Module i.5** 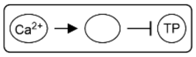  |
| $n$ Hill coefficient | $n$ | $n$ | $n$ |
| $k$ curve midpoint | $k = K$ | $k = \sqrt[n]{\frac{\beta \cdot a + d}{\gamma \cdot a + d}} \cdot K$ | $k = \sqrt[n]{\frac{a + \beta \cdot d}{a + \gamma \cdot d}} \cdot K$ |
| $b$ activity at 0 $Ca^{2+}$ | $b = \beta$ | $b = \frac{\beta \cdot a}{\beta \cdot a + d}$ | $b = \frac{a}{a + \beta \cdot d}$ |
| $c$ activity at high $Ca^{2+}$ | $c = \gamma$ | $c = \frac{\gamma \cdot a}{\gamma \cdot a + d}$ | $c = \frac{a}{a + \gamma \cdot d}$ |
| $\tau_0$ time constant a 0 $Ca^{2+}$ | --- | $\tau_0 = \frac{1}{\beta \cdot a + d}$ | $\tau_0 = \frac{1}{a + \beta \cdot d}$ |
| $\Delta\tau$ change of $\tau$ with $Ca^{2+}$ | --- | $\Delta\tau = -\frac{(\gamma - \beta) \cdot a}{\gamma \cdot a + d}$ | $\Delta\tau = -\frac{(\gamma - \beta) \cdot d}{a + \gamma \cdot d}$ |
| $\boldsymbol{\beta > \gamma}$ | **Module 2** 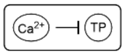  | **Module i.4** 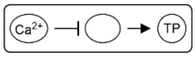  | **Module i.6** 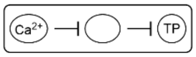  |
| $n$ Hill coefficient | $n$ | $n$ | $n$ |
| $k$ curve midpoint | $k = K$ | $k = \sqrt[n]{\frac{\beta \cdot a + d}{\gamma \cdot a + d}} \cdot K$ | $k = \sqrt[n]{\frac{a + \beta \cdot d}{a + \gamma \cdot d}} \cdot K$ |
| $b$ activity at 0 $Ca^{2+}$ | $b = \beta$ | $b = \frac{\beta \cdot a}{\beta \cdot a + d}$ | $b = \frac{a}{a + \beta \cdot d}$ |
| $c$ activity at high $Ca^{2+}$ | $c = \gamma$ | $c = \frac{\gamma \cdot a}{\gamma \cdot a + d}$ | $c = \frac{a}{a + \gamma \cdot d}$ |
| $\tau_0$ time constant a 0 $Ca^{2+}$ | --- | $\tau_0 = \frac{1}{\beta \cdot a + d}$ | $\tau_0 = \frac{1}{a + \beta \cdot d}$ |
| $\Delta\tau$ change of $\tau$ with $Ca^{2+}$ | --- | $\Delta\tau = \frac{(\beta - \gamma) \cdot a}{d + \gamma \cdot a}$ | $\Delta\tau = \frac{(\beta - \gamma) \cdot d}{a + \gamma \cdot d}$ |

**(ii) Catalytic activity of a CBP on the TP**

The second scenario describes the cases, in which a CBP is rapidly modulated by $Ca^{2+}$ (Modules 1 and 2). In a second, slower step the CBP then modulates catalytically the activity of the TP. $Ca^{2+}$-dependent phosphorylation by $Ca^{2+}$-dependent protein kinases, for example, would fall under this scenario. There are four different cases to be considered: (Module ii.3) the CBP is activated by $Ca^{2+}$ and an active CBP activates the TP; (Module ii.4) the CBP is inactivated by $Ca^{2+}$ and an active CBP activates the TP; (Module ii.5) the CBP is activated by $Ca^{2+}$ and an active CBP inactivates the TP; (Module ii.6) the CBP is inactivated by $Ca^{2+}$ and an active CBP inactivates the TP.

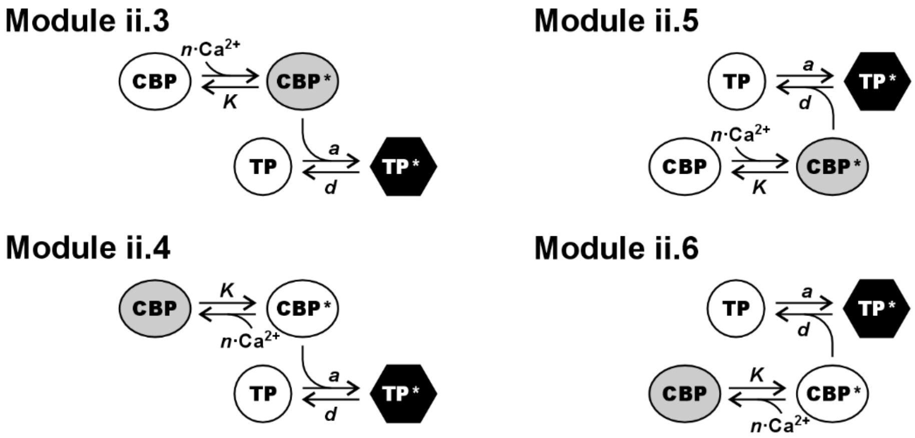


As in the scenario i before, also here, the rapid $Ca^{2+}$-binding reaction was described in generalized form as $\{\beta \cdot K^n + \gamma \cdot [\mathrm{Ca}^{2+}]^n\}/\{K^n + [\mathrm{Ca}^{2+}]^n\}$ with $0 \leq \beta, \gamma \leq 1$. The modules ii.3 and ii.5 are defined for $\beta < \gamma$ and the modules ii.4 and ii.6 are defined for $\beta > \gamma$. If the TP is activated by an active CBP (modules ii.3 and ii.4), the fraction of activated target proteins (*TP**) is determined by the differential equation

$$\frac{d}{dt} TP^*_{ii.x}(t) = a \cdot \frac{\beta \cdot K^n + \gamma \cdot [\mathrm{Ca}^{2+}]^n}{K^n + [\mathrm{Ca}^{2+}]^n} \cdot \left(1 - TP^*_{ii.x}(t)\right) - d \cdot TP^*_{ii.x}(t) \qquad \text{Eqn S16}$$

with *x*=3 or 4 and the reaction constants *a* and *d* that strongly depend on the affinity between CBP and TP and the expression level of CBP; here, *a* contains implicitly the factor $CBP_{total}$, *i.e.* the expression level of the CBP.

In the case that a target protein gets inactivated by an active CBP (modules 5 and 6), the fraction of activated target proteins (*TP**) is determined by the differential equation:

$$\frac{d}{dt} TP^*_{ii.y}(t) = a \cdot \left(1 - TP^*_{ii.y}(t)\right) - d \cdot \frac{\beta \cdot K^n + \gamma \cdot [\mathrm{Ca}^{2+}]^n}{K^n + [\mathrm{Ca}^{2+}]^n} \cdot TP^*_{ii.y}(t) \qquad \text{Eqn S17}$$

with y=5 or 6 and the reaction constants *a* and *d* that strongly depend on the affinity between CBP and TP and the expression level of CBP; here, *d* contains implicitly the factor $CBP_{total}$.

For a complex time-dependent $Ca^{2+}$ signal, these differential equations can only be solved numerically. Nevertheless, a transient $Ca^{2+}$ signal can be interpreted in a staircase manner as a sequence of short $Ca^{2+}$ pulses with constant amplitude. For such time intervals, the differential equations have constant coefficients and can be solved analytically. Upon a $Ca^{2+}$ pulse the time course of the fraction TP* follows in general an exponential decay function (see Eq S7): $TP^*(t) = TP^*_{ss} + (TP^*_0 - TP^*_{ss}) \cdot e^{-t/\tau}$.

Here $TP^*_0$ is the *TP** value at the beginning of the $Ca^{2+}$ pulse and $TP^*_{ss}$ the value in steady state equilibrium. While $TP^*_0$ depends solely on the history of the system, $TP^*_{ss}$ and the time constant $\tau$ are characteristic parameters of the respective module that are presented in the following in more detail.

Module ii.3

$$TP^*_{ss,ii.3} = \frac{\frac{\beta\cdot a}{\beta\cdot a+d}\cdot\left(\frac{\beta\cdot a+d}{\gamma\cdot a+d}\right)\cdot K^n + \frac{\gamma\cdot a}{\gamma\cdot a+d}\cdot\left[\mathrm{Ca}^{2+}\right]^n}{\left(\frac{\beta\cdot a+d}{\gamma\cdot a+d}\right)\cdot K^n + \left[\mathrm{Ca}^{2+}\right]^n} = \frac{\frac{\beta\cdot a/d}{1+\beta\cdot a/d}\cdot\left(\frac{1+\beta\cdot a/d}{1+\gamma\cdot a/d}\right)\cdot K^n + \frac{\gamma\cdot a/d}{1+\gamma\cdot a/d}\cdot\left[\mathrm{Ca}^{2+}\right]^n}{\left(\frac{1+\beta\cdot a/d}{1+\gamma\cdot a/d}\right)\cdot K^n + \left[\mathrm{Ca}^{2+}\right]^n} \qquad \text{Eqn S18}$$

$TP^*_{ss,ii.3}$ increases with increasing [$Ca^{2+}$]. The midpoint of the curve is at $\left[\mathrm{Ca}^{2+}\right]^n_{mid} = \frac{1+\beta\cdot a/d}{1+\gamma\cdot a/d}\cdot K^n$, with $K$ being the midpoint $Ca^{2+}$-concentration and *n* the Hill-coefficient of the calcium-binding protein (CBP). For $\beta < \gamma$, $\left[\mathrm{Ca}^{2+}\right]_{mid,TP3} \le \left[\mathrm{Ca}^{2+}\right]_{mid,CBP}$. Thus, the TP is more sensitive to $Ca^{2+}$ than the CBP.

$$\tau_{ii.3} = \frac{1}{\beta\cdot a+d}\cdot\left(1 - \frac{(\gamma-\beta)\cdot a/d}{1+\gamma\cdot a/d}\cdot\frac{\left[\mathrm{Ca}^{2+}\right]^n}{\left(\frac{1+\beta\cdot a/d}{1+\gamma\cdot a/d}\right)\cdot K^n + \left[\mathrm{Ca}^{2+}\right]^n}\right) \qquad \text{Eqn S19}$$

The time constant of module ii.3, $\tau_{ii.3}$, decreases with increasing [$Ca^{2+}$]. The midpoint of the curve is identical to the midpoint of the $TP^*_{ss,ii.3}$ curve and the slope is determined by the parameter *n*. **Fig. S6** illustrates the influence of the parameters $\beta$, $\gamma$ and $a/d$ (affinity of the CBP to the TP & expression level of the CBP relative to the TP) on the $Ca^{2+}$-dependence of the curves $TP^*_{ss,ii.3}$ and $\tau_{ii.3}$ for fixed *K* and *n*.

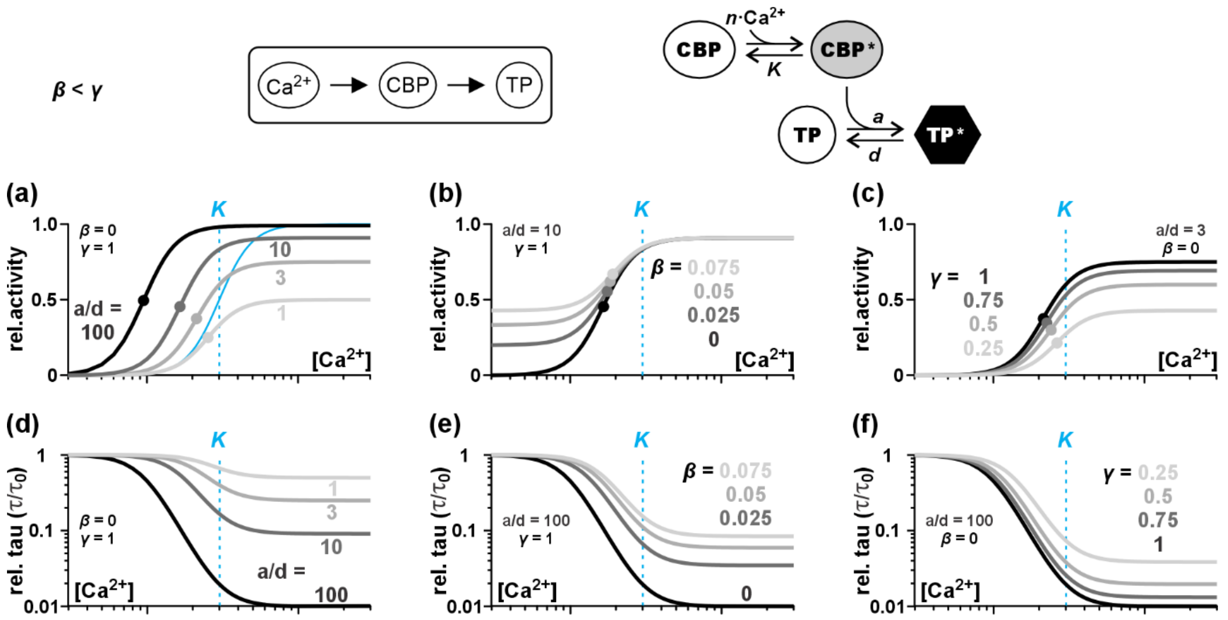


**Fig. S6** Effect of the parameters *β*, *γ* and *a/d* on the $Ca^{2+}$-dependence of module ii.3. The parameters *K*=3 and *n*=4 were fixed. (a-c) $Ca^{2+}$-dependent steady state activity of the target protein in module ii.3 illustrated for different parameters (a) *a/d* (=affinity of the CBP to the TP & expression level of the CBP relative to the TP), (b) *β* (=activity of the CBP at low $Ca^{2+}$), and (c) *γ* (=activity of the CBP at high $Ca^{2+}$). The solid blue curve shows the $Ca^{2+}$-dependence of the CBP for *β*=0/*γ*=1 and the dashed blue lines indicate the midpoint (*K*) of the CBP-curves. The midpoints of the $TP_{ss}$ curves are indicated as dots and grey-scale coded with the curves and the indicated values for the parameters. Please note that with increasing affinity or a higher CBP expression level the midpoint of the $TP_{ss}$ curve shifts to lower $Ca^{2+}$-values, *i.e.* the sensitivity of the TP to $Ca^{2+}$ increases. Panel (b) shows the curves at different *β* for *a/d*=10/*γ*=1, while panel (c) illustrates the dependency on *γ* for *a/d*=3/*β*=0. An increase in *β* and a decrease in *γ* reduce the $Ca^{2+}$ sensitivity of the TP, *i.e.* the midpoint shifts towards the *K* value of the CBP. (d-f) $Ca^{2+}$-dependence of the time constant $\tau$ displayed relative to the $\tau$-value at zero $Ca^{2+}$, $\tau_0$. Please note the logarithmic scale of the y-axes. (d) With increasing affinity of the CBP to the TP or a higher CBP expression (parameter *a/d*) the time constant of the module becomes smaller, i.e. the module reacts faster at higher $Ca^{2+}$. (e-f) An increase in the basal activity of the CBP at low $Ca^{2+}$ (parameter *β*) or a decrease in activity of the CBP at high $Ca^{2+}$ (parameter *γ*) reduce the difference between the time constants at high and at low $Ca^{2+}$.

Module ii.4

$$TP^*_{ss,ii.4} = \frac{\frac{\beta\cdot a}{\beta\cdot a+d}\cdot\left(\frac{\beta\cdot a+d}{\gamma\cdot a+d}\right)\cdot K^n+\frac{\gamma\cdot a}{\gamma\cdot a+d}\cdot\left[\mathrm{Ca}^{2+}\right]^n}{\left(\frac{\beta\cdot a+d}{\gamma\cdot a+d}\right)\cdot K^n+\left[\mathrm{Ca}^{2+}\right]^n} = \frac{\frac{\beta\cdot a/d}{1+\beta\cdot a/d}\cdot\left(\frac{1+\beta\cdot a/d}{1+\gamma\cdot a/d}\right)\cdot K^n+\frac{\gamma\cdot a/d}{1+\gamma\cdot a/d}\cdot\left[\mathrm{Ca}^{2+}\right]^n}{\left(\frac{1+\beta\cdot a/d}{1+\gamma\cdot a/d}\right)\cdot K^n+\left[\mathrm{Ca}^{2+}\right]^n} \qquad \text{Eqn S20}$$

$TP^*_{ss,ii.4}$ decreases with increasing [$Ca^{2+}$]. The midpoint of the curve is at $\left[\mathrm{Ca}^{2+}\right]^n_{mid} = \frac{1+\beta\cdot a/d}{1+\gamma\cdot a/d}\cdot K^n$, with *K* being the midpoint $Ca^{2+}$-concentration and *n* the Hill-coefficient of the calcium-binding protein (CBP). For $\beta > \gamma$, $\left[\mathrm{Ca}^{2+}\right]_{mid,TP4} \geq \left[\mathrm{Ca}^{2+}\right]_{mid,CBP}$. Thus, the TP is less sensitive to $Ca^{2+}$ than the CBP.

$$\tau_{ii.4} = \frac{1}{\beta\cdot a+d}\cdot\left(1+\frac{(\beta-\gamma)\cdot a/d}{1+\gamma\cdot a/d}\cdot\frac{\left[\mathrm{Ca}^{2+}\right]^n}{\left(\frac{1+\beta\cdot a/d}{1+\gamma\cdot a/d}\right)\cdot K^n+\left[\mathrm{Ca}^{2+}\right]^n}\right) \qquad \text{Eqn S21}$$

The time constant of module ii.4, $\tau_{ii4}$, increases with increasing [$Ca^{2+}$]. The midpoint of the curve is identical to the midpoint of the $TP^*_{ss,ii.4}$ curve and the slope is determined by the parameter *n*. **Fig. S7** illustrates the influence of the parameters $\beta$, $\gamma$ and $a/d$ (affinity of the CBP to the TP & expression level of the CBP relative to the TP) on the $Ca^{2+}$-dependence of the curves $TP^*_{ss,ii.4}$ and $\tau_{ii.4}$ for fixed *K* and *n*.

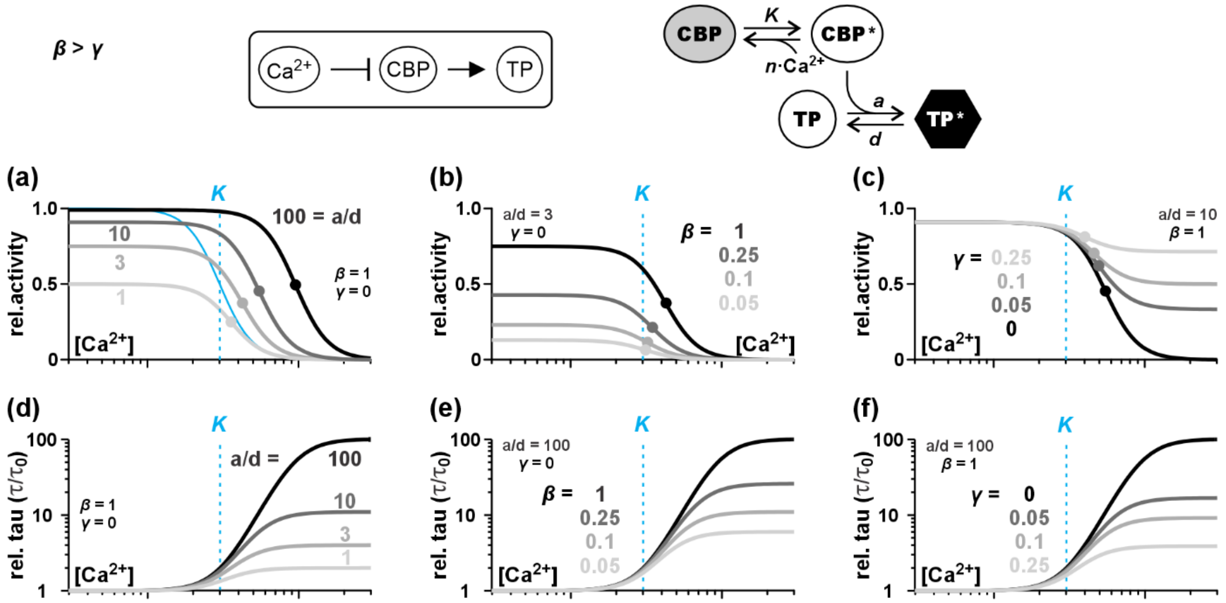


**Fig. S7** Effect of the parameters *β*, *γ* and *a/d* on the $Ca^{2+}$-dependence of module ii.4. The parameters *K*=3 and *n*=4 were fixed. (a-c) $Ca^{2+}$-dependent steady state activity of the target protein in module ii.4 illustrated for different parameters (a) *a/d* (=affinity of the CBP to the TP & expression level of the CBP relative to the TP), (b) *β* (=activity of the CBP at low $Ca^{2+}$), and (c) *γ* (=activity of the CBP at high $Ca^{2+}$). The solid blue curve shows the $Ca^{2+}$-dependence of the CBP for *β*=1/*γ*=0 and the dashed blue lines indicate the midpoint (*K*) of the CBP-curves. The midpoints of the $TP_{ss}$ curves are indicated as dots and grey-scale coded with the curves and the indicated values for the parameters. Please note that with increasing affinity or a higher CBP expression level the midpoint of the $TP_{ss}$ curve shifts to higher $Ca^{2+}$-values, *i.e.* the sensitivity of the TP to $Ca^{2+}$ decreases. Panel (b) shows the curves at different *β* for *a/d*=3/*γ*=0, while panel (c) illustrates the dependency on *γ* for *a/d*=10/*β*=1. A decrease in *β* and an increase in *γ* enhance the $Ca^{2+}$ sensitivity of the TP, *i.e.* the midpoint shifts towards the *K* value of the CBP. (d-f) $Ca^{2+}$-dependence of the time constant $\tau$ displayed relative to the $\tau$-value at zero $Ca^{2+}$, $\tau_0$. Please note the logarithmic scale of the y-axes. (d) With increasing affinity of the CBP to the TP or a higher CBP expression (parameter *a/d*) the time constant of the module increases, *i.e.* the module reacts more slowly at higher $Ca^{2+}$. (e-f) A decrease in the basal activity of the CBP at low $Ca^{2+}$ (parameter *β*) or an increase in activity of the CBP at high $Ca^{2+}$ (parameter *γ*) reduce the difference between the time constants at high and at low $Ca^{2+}$.

Module ii.5

$$TP^*_{ss,ii.5} = \frac{\frac{a}{a+\beta\cdot d}\cdot\left(\frac{a+\beta\cdot d}{a+\gamma\cdot d}\right)\cdot K^n+\frac{a}{a+\gamma\cdot d}\cdot\left[\mathrm{Ca}^{2+}\right]^n}{\left(\frac{a+\beta\cdot d}{a+\gamma\cdot d}\right)\cdot K^n+\left[\mathrm{Ca}^{2+}\right]^n} = \frac{\frac{1}{1+\beta\cdot d/a}\cdot\left(\frac{1+\beta\cdot d/a}{1+\gamma\cdot d/a}\right)\cdot K^n+\frac{1}{1+\gamma\cdot d/a}\cdot\left[\mathrm{Ca}^{2+}\right]^n}{\left(\frac{1+\beta\cdot d/a}{1+\gamma\cdot d/a}\right)\cdot K^n+\left[\mathrm{Ca}^{2+}\right]^n} \qquad \text{Eqn S22}$$

$TP^*_{ss,ii.5}$ decreases with increasing [$Ca^{2+}$]. The midpoint of the curve is at $\left[\mathrm{Ca}^{2+}\right]^n_{mid} = \frac{1+\beta\cdot d/a}{1+\gamma\cdot d/a}\cdot K^n$, with *K* being the midpoint $Ca^{2+}$-concentration and *n* the Hill-coefficient of the calcium-binding protein (CBP). For $\beta < \gamma$, $\left[\mathrm{Ca}^{2+}\right]_{mid,TP5} \leq \left[\mathrm{Ca}^{2+}\right]_{mid,CBP}$. Thus, the TP is more sensitive to $Ca^{2+}$ than the CBP.

$$\tau_{ii.5} = \frac{1}{a+\beta\cdot d}\cdot\left(1-\frac{(\gamma-\beta)\cdot d/a}{1+\gamma\cdot d/a}\cdot\frac{\left[\mathrm{Ca}^{2+}\right]^n}{\left(\frac{1+\beta\cdot d/a}{1+\gamma\cdot d/a}\right)\cdot K^n+\left[\mathrm{Ca}^{2+}\right]^n}\right) \qquad \text{Eqn S23}$$

The time constant of module ii.5, $\tau_{ii.5}$, decreases with increasing [$Ca^{2+}$]. The midpoint of the curve is identical to the midpoint of the $TP^*_{ss,ii.5}$ curve and the slope is determined by the parameter *n*. **Fig. S8** illustrates the influence of the parameters $\beta$, $\gamma$ and $d/a$ (affinity of the CBP to the TP & expression level of the CBP relative to the TP) on the $Ca^{2+}$-dependence of the curves $TP^*_{ss,ii5}$ and $\tau_{ii5}$ for fixed *K* and *n*.

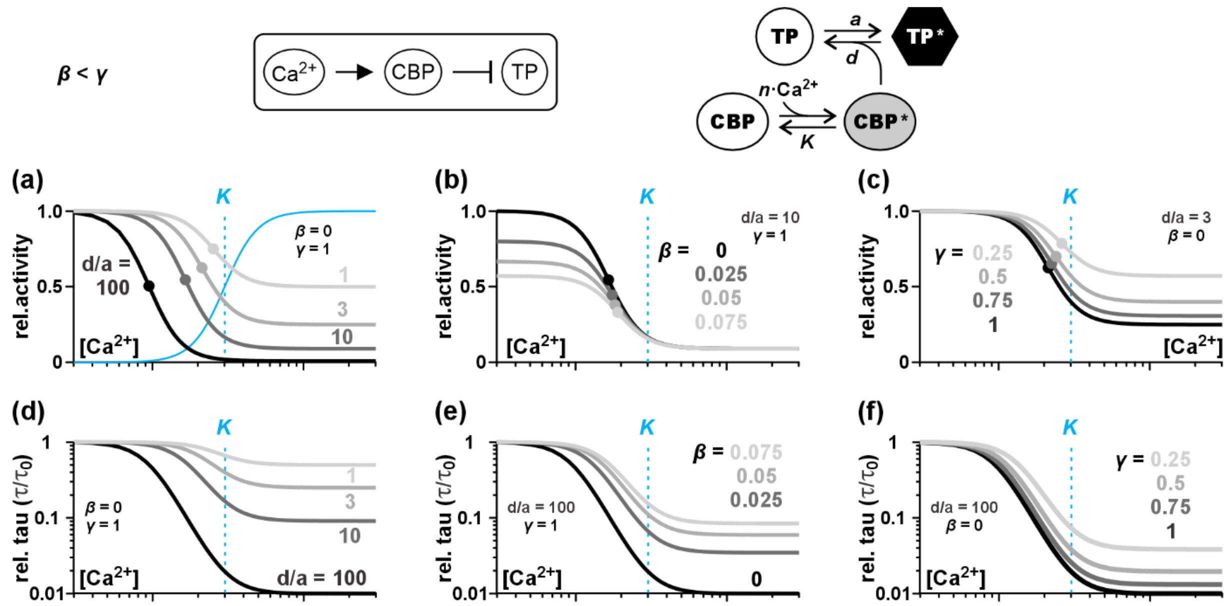


**Fig. S8** Effect of the parameters *β*, *γ* and *d/a* on the $Ca^{2+}$-dependence of module ii.5. The parameters *K*=3 and *n*=4 were fixed. (a-c) $Ca^{2+}$-dependent steady state activity of the target protein in module ii.5 illustrated for different parameters (a) *d/a* (=affinity of the CBP to the TP & expression level of the CBP relative to the TP), (b) *β* (=activity of the CBP at low $Ca^{2+}$), and (c) *γ* (=activity of the CBP at high $Ca^{2+}$). The solid blue curve shows the $Ca^{2+}$-dependence of the CBP for *β*=0/*γ*=1 and the dashed blue lines indicate the midpoint (*K*) of the CBP-curves. The midpoints of the $TP_{ss}$ curves are indicated as dots and grey-scale coded with the curves and the indicated values for the parameters. Please note that with increasing affinity or a higher CBP expression level the midpoint of the $TP_{ss}$ curve shifts to lower $Ca^{2+}$-values, *i.e.* the sensitivity of the TP to $Ca^{2+}$ increases. Panel (b) shows the curves at different *β* for *d/a*=10/*γ*=1, while panel (c) illustrates the dependency on *γ* for *d/a*=3/*β*=0. An increase in *β* and a decrease in *γ* reduce the $Ca^{2+}$ sensitivity of the TP, *i.e.* the midpoint shifts towards the *K* value of the CBP. (d-f) $Ca^{2+}$-dependence of the time constant $\tau$ displayed relative to the $\tau$-value at zero $Ca^{2+}$, $\tau_0$. Please note the logarithmic scale of the y-axes. (d) With increasing affinity of the CBP to the TP or a higher CBP expression (parameter *d/a*) the time constant of the module becomes smaller, i.e. the module reacts faster at higher $Ca^{2+}$. (e-f) An increase in the basal activity of the CBP at low $Ca^{2+}$ (parameter *β*) or a decrease in activity of the CBP at high $Ca^{2+}$ (parameter *γ*) reduce the difference between the time constants at high and at low $Ca^{2+}$.

Module ii.6

$$TP^*_{ss,ii.6} = \frac{\frac{a}{a+\beta\cdot d}\cdot\left(\frac{a+\beta\cdot d}{a+\gamma\cdot d}\right)\cdot K^n + \frac{a}{a+\gamma\cdot d}\cdot\left[\mathrm{Ca}^{2+}\right]^n}{\left(\frac{a+\beta\cdot d}{a+\gamma\cdot d}\right)\cdot K^n+\left[\mathrm{Ca}^{2+}\right]^n} = \frac{\frac{1}{1+\beta\cdot d/a}\cdot\left(\frac{1+\beta\cdot d/a}{1+\gamma\cdot d/a}\right)\cdot K^n+\frac{1}{1+\gamma\cdot d/a}\cdot\left[\mathrm{Ca}^{2+}\right]^n}{\left(\frac{1+\beta\cdot d/a}{1+\gamma\cdot d/a}\right)\cdot K^n+\left[\mathrm{Ca}^{2+}\right]^n} \qquad \text{Eqn S24}$$

$TP^*_{ss,ii.6}$ increases with increasing [$Ca^{2+}$]. The midpoint of the curve is at $\left[\mathrm{Ca}^{2+}\right]^n_{mid} = \frac{1+\beta\cdot d/a}{1+\gamma\cdot d/a}\cdot K^n$, with *K* being the midpoint $Ca^{2+}$-concentration and *n* the Hill-coefficient of the calcium-binding protein (CBP). For $\beta > \gamma$, $\left[\mathrm{Ca}^{2+}\right]_{mid,TP6} \geq \left[\mathrm{Ca}^{2+}\right]_{mid,CBP}$. Thus, the TP is less sensitive to $Ca^{2+}$ than the CBP.

$$\tau_{ii.6} = \frac{1}{a+\beta\cdot d}\cdot\left(1+\frac{(\beta-\gamma)\cdot d/a}{1+\gamma\cdot d/a}\cdot\frac{\left[\mathrm{Ca}^{2+}\right]^n}{\left(\frac{1+\beta\cdot d/a}{1+\gamma\cdot d/a}\right)\cdot K^n+\left[\mathrm{Ca}^{2+}\right]^n}\right) \qquad \text{Eqn S25}$$

The time constant of module 6, $\tau_{ii.6}$, increases with increasing [$Ca^{2+}$]. The midpoint of the curve is identical to the midpoint of the $TP^*_{ss,ii.6}$ curve and the slope is determined by the parameter *n*. **Fig. S9** illustrates the influence of the parameters $\beta$, $\gamma$ and $d/a$ (affinity of the CBP to the TP & expression level of the CBP relative to the TP) on the $Ca^{2+}$-dependence of the curves $TP^*_{ss,ii.6}$ and $\tau_{ii.6}$ for fixed *K* and *n*.

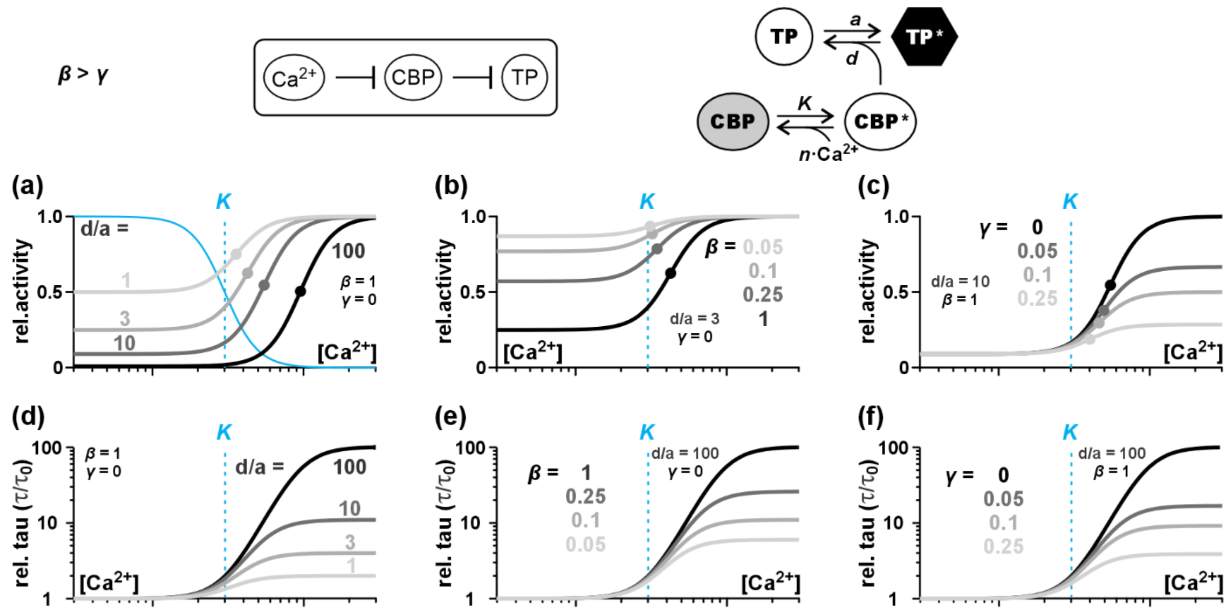


**Fig. S9** Effect of the parameters *β*, *γ* and *d/a* on the $Ca^{2+}$-dependence of module ii.6. The parameters *K*=3 and *n*=4 were fixed. (a-c) $Ca^{2+}$-dependent steady state activity of the target protein in module ii.6 illustrated for different parameters (a) *d/a* (=affinity of the CBP to the TP & expression level of the CBP relative to the TP), (b) *β* (=activity of the CBP at low $Ca^{2+}$), and (c) *γ* (=activity of the CBP at high $Ca^{2+}$). The solid blue curve shows the $Ca^{2+}$-dependence of the CBP for *β*=1/*γ*=0 and the dashed blue lines indicate the midpoint (*K*) of the CBP-curves. The midpoints of the $TP_{ss}$ curves are indicated as dots and grey-scale coded with the curves and the indicated values for the parameters. Please note that with increasing affinity or a higher CBP expression level the midpoint of the $TP_{ss}$ curve shifts to higher $Ca^{2+}$-values, *i.e.* the sensitivity of the TP to $Ca^{2+}$ decreases. Panel (b) shows the curves at different *β* for *d/a*=3/*γ*=0, while panel (c) illustrates the dependency on *γ* for *d/a*=10/*β*=1. A decrease in *β* and an increase in *γ* enhance the $Ca^{2+}$ sensitivity of the TP, *i.e.* the midpoint shifts towards the *K* value of the CBP. (d-f) $Ca^{2+}$-dependence of the time constant $\tau$ displayed relative to the $\tau$-value at zero $Ca^{2+}$, $\tau_0$. Please note the logarithmic scale of the y-axes. (d) With increasing affinity of the CBP to the TP or a higher CBP expression (parameter *d/a*) the time constant of the module increases, *i.e.* the module reacts more slowly at higher $Ca^{2+}$. (e-f) A decrease in the basal activity of the CBP at low $Ca^{2+}$ (parameter *β*) or an increase in activity of the CBP at high $Ca^{2+}$ (parameter *γ*) reduce the difference between the time constants at high and at low $Ca^{2+}$.

**Table S2**. Conversion between the parameter sets $(n, K, \beta, \gamma, a, d)$ and $(n, k, b, c, \tau_0, \Delta\tau)$ for the case of catalytic modulation of the TP by the CBP.

$$TP^*(t) = TP^*_{ss} + (TP^*_0 - TP^*_{ss}) \cdot e^{-t/\tau}$$

$$TP^*_{ss} = \frac{b \cdot k^n + c \cdot [\mathrm{Ca}^{2+}]^n}{k^n + [\mathrm{Ca}^{2+}]^n} \qquad \tau = \tau_0 \cdot \left(1 + \Delta\tau \cdot \frac{[\mathrm{Ca}^{2+}]^n}{k^n + [\mathrm{Ca}^{2+}]^n}\right)$$

| $\boldsymbol{\beta < \gamma}$ | **Module 1**<br>Ca$^{2+}$ → TP | **Module ii.3 ***<br>Ca$^{2+}$ → CBP → TP | **Module ii.5 ****<br>Ca$^{2+}$ → CBP ⊣ TP |
|---|---|---|---|
| *n*<br>Hill coefficient | *n* | *n* | *n* |
| *k*<br>curve midpoint | $k = K$ | $k = \sqrt[n]{\frac{\beta \cdot a + d}{\gamma \cdot a + d}} \cdot K$ | $k = \sqrt[n]{\frac{a + \beta \cdot d}{a + \gamma \cdot d}} \cdot K$ |
| *b*<br>activity at 0 Ca$^{2+}$ | $b = \beta$ | $b = \frac{\beta \cdot a}{\beta \cdot a + d}$ | $b = \frac{a}{a + \beta \cdot d}$ |
| *c*<br>activity at high Ca$^{2+}$ | $c = \gamma$ | $c = \frac{\gamma \cdot a}{\gamma \cdot a + d}$ | $c = \frac{a}{a + \gamma \cdot d}$ |
| $\tau_0$<br>time constant a 0 Ca$^{2+}$ | --- | $\tau_0 = \frac{1}{\beta \cdot a + d}$ | $\tau_0 = \frac{1}{a + \beta \cdot d}$ |
| $\Delta\tau$<br>change of $\tau$ with Ca$^{2+}$ | --- | $\Delta\tau = -\frac{(\gamma - \beta) \cdot a}{\gamma \cdot a + d} < 0$ | $\Delta\tau = -\frac{(\gamma - \beta) \cdot d}{a + \gamma \cdot d} < 0$ |
| $\boldsymbol{\beta > \gamma}$ | **Module 2**<br>Ca$^{2+}$ ⊣ TP | **Module ii.4 ***<br>Ca$^{2+}$ ⊣ CBP → TP | **Module ii.6 ****<br>Ca$^{2+}$ ⊣ CBP ⊣ TP |
| *n*<br>Hill coefficient | *n* | *n* | *n* |
| *k*<br>curve midpoint | $k = K$ | $k = \sqrt[n]{\frac{\beta \cdot a + d}{\gamma \cdot a + d}} \cdot K$ | $k = \sqrt[n]{\frac{a + \beta \cdot d}{a + \gamma \cdot d}} \cdot K$ |
| *b*<br>activity at 0 Ca$^{2+}$ | $b = \beta$ | $b = \frac{\beta \cdot a}{\beta \cdot a + d}$ | $b = \frac{a}{a + \beta \cdot d}$ |
| *c*<br>activity at high Ca$^{2+}$ | $c = \gamma$ | $c = \frac{\gamma \cdot a}{\gamma \cdot a + d}$ | $c = \frac{a}{a + \gamma \cdot d}$ |
| $\tau_0$<br>time constant a 0 Ca$^{2+}$ | --- | $\tau_0 = \frac{1}{\beta \cdot a + d}$ | $\tau_0 = \frac{1}{a + \beta \cdot d}$ |
| $\Delta\tau$<br>change of $\tau$ with Ca$^{2+}$ | --- | $\Delta\tau = \frac{(\beta - \gamma) \cdot a}{\gamma \cdot a + d} > 0$ | $\Delta\tau = \frac{(\beta - \gamma) \cdot d}{a + \gamma \cdot d} > 0$ |

* Please note: The parameter *a* contains implicitly the factor $CBP_{total}$, *i.e.* the expression level of the CBP.
** Please note: The parameter *d* contains implicitly the factor $CBP_{total}$, *i.e.* the expression level of the CBP.
Thus, the parameter set $(k, b, c, \tau_0, \Delta\tau)$ varies with the expression level of the CBP.

**(iii) Binding of a CBP to the TP**

The third scenario describes the cases, in which a CBP is rapidly modulated by $Ca^{2+}$ (Modules 1 and 2) in a first step. In a second, slower step the CBP then binds to the TP and modulates its activity. Interaction of the target protein with calmodulin, for example, would fall under this scenario. There are four different cases to be considered: (Module iii.3) the CBP is activated by $Ca^{2+}$, an active CBP binds to the TP and activates it; (Module iii.4) the CBP is inactivated by $Ca^{2+}$, an active CBP binds to the TP and activates it; (Module iii.5) the CBP is activated by $Ca^{2+}$, an active CBP binds to the TP and inactivates it; (Module iii.6) the CBP is inactivated by $Ca^{2+}$, an active CBP binds to the TP and inactivates it.

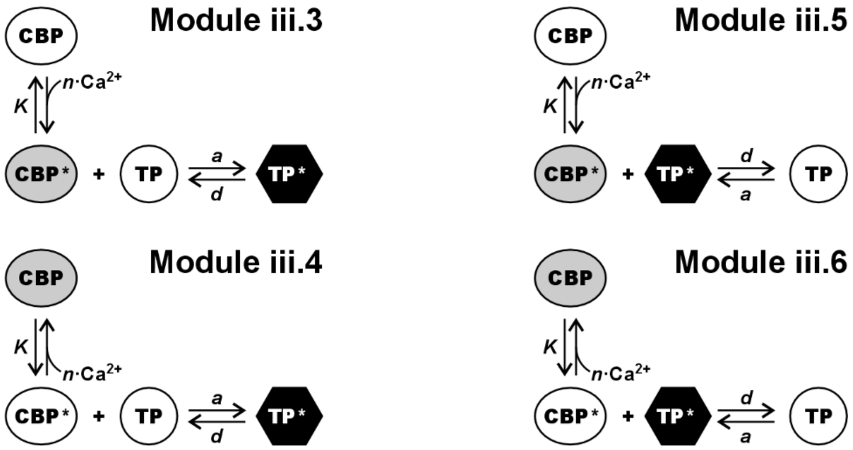


The Target-Protein (total expression level $TP_{total}$) and the Calcium-Binding-Protein (total expression level $CBP_{total}$) interact in a bimolecular chemical reaction. When *TargetP* stands for the amount of inactive and *TargetP** for the amount of active target proteins, the definitions $TP^* = TargetP^*/TP_{total}$ and $TP = TargetP/TP_{total}$ indicate the fractions of active and inactive target proteins respectively. The concentration of CBP* is determined by $CBP^*_{iii.3} = f\cdot(CBP_{total} - TargetP^*)$ in Module iii.3, $CBP^*_{iii.4} = f\cdot(CBP_{total} - TargetP^*)$ in Module iii.4, $CBP^*_{iii.5} = f\cdot(CBP_{total} - [TP_{total} - TargetP^*])$ in Module iii.5, and $CBP^*_{iii.6} = f\cdot(CBP_{total} - [TP_{total} - TargetP^*])$ in Module iii.6, with $f = \left(\beta\cdot K^n + \gamma\cdot[\text{Ca}^{2+}]^n\right)/\left(K^n + [\text{Ca}^{2+}]^n\right)$ ($\beta < \gamma$ in modules iii.3 and iii.5; $\beta > \gamma$ in modules iii.4 and iii.6). With $CT = CBP_{total}/TP_{total}$ (expression relation between CBP and TP), the change of *TP** in time is given by the differential equations:

$$\frac{d}{dt}TP^*_{iii.x} = a\cdot(1-TP^*_{iii.x})\cdot f\cdot(CT-TP^*_{iii.x}) - d\cdot TP^*_{iii.x} \qquad \text{Eqn S26}$$

with *x*=3 or 4; here, the rate constant *a* contains implicitly the factor $TP_{total}$.

$$\frac{d}{dt}TP^*_{iii.y} = a\cdot\left(1-TP^*_{iii.y}\right) - d\cdot TP^*_{iii.y}\cdot f\cdot\left(CT-\left(1-TP^*_{iii.y}\right)\right) \qquad \text{Eqn S27}$$

with *x*=5 or 6; here, the rate constant *d* contains implicitly the factor $TP_{total}$. Under constant $[Ca^{2+}]$ conditions (see remark in section ii), the solutions of these equations are

$$TP^*_{iii.x}(t) = \frac{1}{2}\cdot\left[CT + \frac{a\cdot f+d}{a\cdot f} - \sqrt{\left(CT+\frac{a\cdot f+d}{a\cdot f}\right)^2 - 4\cdot CT}\cdot\tanh\left(\frac{a\cdot f}{2}\cdot\sqrt{\left(CT+\frac{a\cdot f+d}{a\cdot f}\right)^2 - 4\cdot CT}\cdot t\right)\right] \qquad \text{Eqn S28}$$

$$TP^*_{iii.y}(t) = \frac{1}{2}\cdot\left[\frac{d\cdot f-a}{d\cdot f} - CT + \sqrt{\left(CT+\frac{a+d\cdot f}{d\cdot f}\right)^2 - 4\cdot CT}\cdot\tanh\left(\frac{d\cdot f}{2}\cdot\sqrt{\left(CT+\frac{a+d\cdot f}{d\cdot f}\right)^2 - 4\cdot CT}\cdot t\right)\right] \qquad \text{Eqn S29}$$

These equations can be written as:

$$TP^*_{iii}(t) = TP^*_{0,iii} + \left(TP^*_{ss,iii} - TP^*_{0,iii}\right)\cdot\tanh\left(\ln 2\cdot {}^{t}/_{\tau_{iii}}\right) \qquad \text{Eqn S30}$$

Here $TP^*_{0,iii}$ is the $TP^*_{iii}$ value at the beginning of the $Ca^{2+}$ pulse and $TP^*_{ss,iii}$ the value in steady state equilibrium. While $TP^*_{0,iii}$ depends solely on the history of the system, $TP^*_{ss,iii}$ and the time constant $\tau_{iii}$ are characteristic parameters of the respective module that are presented in the following in more detail.

Module iii.3

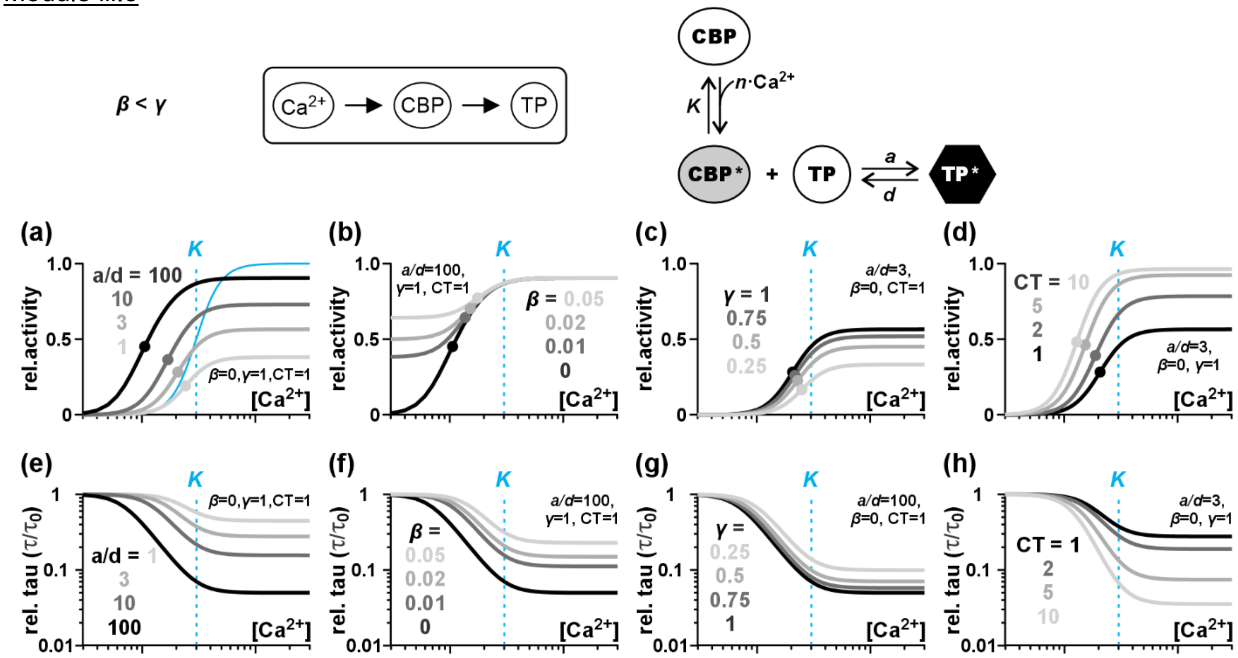


**Fig. S10** Effect of the parameters *a/d*, $\beta$, $\gamma$, and *CT* on the $Ca^{2+}$-dependence of module iii.3. The parameters *K*=3 and *n*=4 were fixed. (a-d) $Ca^{2+}$-dependent steady state activity of the target protein in module iii.3 illustrated for different parameters (a) *a/d* (=affinity of the CBP to the TP), (b) $\beta$ (=activity of the CBP at low $Ca^{2+}$), (c) $\gamma$ (=activity of the CBP at high $Ca^{2+}$), and (d) *CT* (= expression of the CBP relative to the TP, $CT = CBP_{total}/TP_{total}$). The solid blue curve shows the $Ca^{2+}$-dependence of the CBP for $\beta$=0/$\gamma$=1 and the dashed blue lines indicate the midpoint (*K*) of the CBP-curves. The midpoints of the $TP_{ss}$ curves are indicated as dots and grey-scale coded with the curves and the indicated values for the parameters. Please note that with increasing affinity or an increasing CBP expression the midpoint of the $TP_{ss}$ curve shifts to lower $Ca^{2+}$-values, *i.e.* the sensitivity of the TP to $Ca^{2+}$ increases. Panel (b) shows the curves at different $\beta$ for fixed values *a/d*=100/$\gamma$=1/*CT*=1, panel (c) for different $\gamma$ [*a/d*=3/$\beta$=0/*CT*=1], while panel (d) illustrates the dependency on CT [*a/d*=3/$\beta$=0/$\gamma$=1]. (e-h) $Ca^{2+}$-dependence of the time constant $\tau$ displayed relative to the $\tau$-value at zero $Ca^{2+}$, $\tau_0$. Please note the logarithmic scale of the y-axes. With increasing affinity of the CBP to the TP (panel e, parameter *a/d*), or a higher CBP expression (panel h, parameter *CT*) the time constant of the module becomes smaller, i.e. the module reacts faster at higher $Ca^{2+}$. (f,g) An increase in the basal activity of the CBP at low $Ca^{2+}$ (parameter $\beta$) or a decrease in activity of the CBP at high $Ca^{2+}$ (parameter $\gamma$) reduce the difference between the time constants at high and at low $Ca^{2+}$.

$$TP^*_{ss,iii.3} = \frac{1}{2}\cdot\left[CT + \frac{(1+\beta\cdot a/d)\cdot K^n + (1+\gamma\cdot a/d)\cdot\left[\mathrm{Ca}^{2+}\right]^n}{\beta\cdot a/d\cdot K^n + \gamma\cdot a/d\cdot\left[\mathrm{Ca}^{2+}\right]^n} - \sqrt{\left(CT + \frac{(1+\beta\cdot a/d)\cdot K^n + (1+\gamma\cdot a/d)\cdot\left[\mathrm{Ca}^{2+}\right]^n}{\beta\cdot a/d\cdot K^n + \gamma\cdot a/d\cdot\left[\mathrm{Ca}^{2+}\right]^n}\right)^2 - 4\cdot CT}\right] \quad \text{Eqn S31}$$

$$\tau_{iii.3} = \frac{2\cdot\ln 2}{a}\cdot\frac{K^n + \left[\mathrm{Ca}^{2+}\right]^n}{\beta\cdot K^n + \gamma\cdot\left[\mathrm{Ca}^{2+}\right]^n}\cdot\left(\left(CT + \frac{(1+\beta\cdot a/d)\cdot K^n + (1+\gamma\cdot a/d)\cdot\left[\mathrm{Ca}^{2+}\right]^n}{\beta\cdot a/d\cdot K^n + \gamma\cdot a/d\cdot\left[\mathrm{Ca}^{2+}\right]^n}\right)^2 - 4\cdot CT\right)^{-\frac{1}{2}} \quad \text{Eqn S32}$$

$TP^*_{ss,iii.3}$ increases with increasing [$Ca^{2+}$]. The midpoint calcium concentration of the curve is smaller than the midpoint $Ca^{2+}$-concentration of the calcium-binding protein (CBP), $[\mathrm{Ca}^{2+}]_{mid,TP3} \leq [\mathrm{Ca}^{2+}]_{mid,CBP}$. Thus, the TP is more sensitive to $Ca^{2+}$ than the CBP. The time constant of module iii.3, $\tau_{iii.3}$, decreases with increasing [$Ca^{2+}$]. Both, midpoint and slope of the curves can be different from midpoint and slope of the respective $TP^*_{ss,iii.3}$ curves. **Fig. S10** illustrates the influence of the parameters $a/d$ (affinity of the CBP to the TP), $\beta$ (=activity of the CBP at low $Ca^{2+}$), $\gamma$ (=activity of the CBP at high $Ca^{2+}$), and *CT* (= expression of the CBP relative to the TP) on the $Ca^{2+}$-dependence of the curves $TP^*_{ss,iii.3}$ and $\tau_{iii.3}$ for fixed *K* and *n*.

Module iii.4

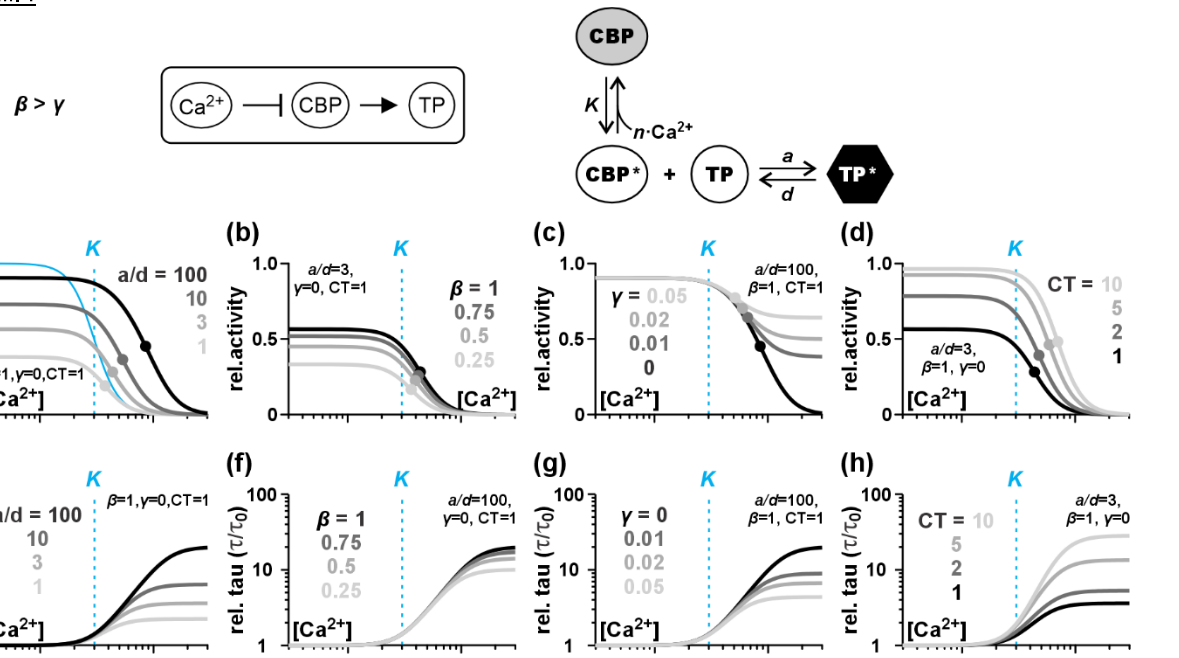


**Fig. S11** Effect of the parameters *a/d*, *β*, *γ*, and *CT* on the $Ca^{2+}$-dependence of module iii.4. The parameters *K*=3 and *n*=4 were fixed. (a-d) $Ca^{2+}$-dependent steady state activity of the target protein in module iii.4 illustrated for different parameters (a) *a/d* (=affinity of the CBP to the TP), (b) *β* (=activity of the CBP at low $Ca^{2+}$), (c) *γ* (=activity of the CBP at high $Ca^{2+}$), and (d) *CT* (= expression of the CBP relative to the TP, $CT = CBP_{total}/TP_{total}$). The solid blue curve shows the $Ca^{2+}$-dependence of the CBP for *β*=1/*γ*=0 and the dashed blue lines indicate the midpoint (*K*) of the CBP-curves. The midpoints of the $TP_{ss}$ curves are indicated as dots and grey-scale coded with the curves and the indicated values for the parameters. Please note that with increasing affinity or an increasing CBP expression the midpoint of the $TP_{ss}$ curve shifts to higher $Ca^{2+}$-values, *i.e.* the sensitivity of the TP to $Ca^{2+}$ decreases. Panel (b) shows the curves at different *β* for fixed values *a/d*=3/*γ*=0/*CT*=1, panel (c) for different *γ* [*a/d*=100/*β*=1/*CT*=1], while panel (d) illustrates the dependency on CT [*a/d*=3/*β*=1/*γ*=0]. (e-h) $Ca^{2+}$-dependence of the time constant $\tau$ displayed relative to the $\tau$-value at zero $Ca^{2+}$, $\tau_0$. Please note the logarithmic scale of the y-axes. With increasing affinity of the CBP to the TP (panel e, parameter *a/d*), or a higher CBP expression (panel h, parameter *CT*) the time constant of the module increases, *i.e.* the module reacts more slowly at higher $Ca^{2+}$. (f,g) An increase in the basal activity of the CBP at low $Ca^{2+}$ (parameter *β*) or a decrease in activity of the CBP at high $Ca^{2+}$ (parameter *γ*) reduce the difference between the time constants at high and at low $Ca^{2+}$.

$$TP^*_{ss,iii.4} = \frac{1}{2}\cdot\left[CT + \frac{(1+\beta\cdot a/d)\cdot K^n + (1+\gamma\cdot a/d)\cdot\left[\mathrm{Ca}^{2+}\right]^n}{\beta\cdot a/d\cdot K^n + \gamma\cdot a/d\cdot\left[\mathrm{Ca}^{2+}\right]^n} - \sqrt{\left(CT + \frac{(1+\beta\cdot a/d)\cdot K^n + (1+\gamma\cdot a/d)\cdot\left[\mathrm{Ca}^{2+}\right]^n}{\beta\cdot a/d\cdot K^n + \gamma\cdot a/d\cdot\left[\mathrm{Ca}^{2+}\right]^n}\right)^2 - 4\cdot CT}\right] \quad \text{Eqn S33}$$

$$\tau_{iii.4} = \frac{2\cdot\ln 2}{a}\cdot\frac{K^n + \left[\mathrm{Ca}^{2+}\right]^n}{\beta\cdot K^n + \gamma\cdot\left[\mathrm{Ca}^{2+}\right]^n}\cdot\left(\left(CT + \frac{(1+\beta\cdot a/d)\cdot K^n + (1+\gamma\cdot a/d)\cdot\left[\mathrm{Ca}^{2+}\right]^n}{\beta\cdot a/d\cdot K^n + \gamma\cdot a/d\cdot\left[\mathrm{Ca}^{2+}\right]^n}\right)^2 - 4\cdot CT\right)^{-\frac{1}{2}} \quad \text{Eqn S34}$$

$TP^*_{ss,iii.4}$ decreases with increasing [$Ca^{2+}$]. The midpoint calcium concentration of the curve is larger than the midpoint $Ca^{2+}$-concentration of the calcium-binding protein (CBP), $[\mathrm{Ca}^{2+}]_{mid,TP4} \geq [\mathrm{Ca}^{2+}]_{mid,CBP}$. Thus, the TP is less sensitive to $Ca^{2+}$ than the CBP. The time constant of module iii.4, $\tau_{iii.4}$, increases with increasing [$Ca^{2+}$]. Both, midpoint and slope of the curves can be different from midpoint and slope of the respective $TP^*_{ss,iii.4}$ curves. **Fig. S11** illustrates the influence of the parameters $a/d$ (affinity of the CBP to the TP, $\beta$ (=activity of the CBP at low $Ca^{2+}$), $\gamma$ (=activity of the CBP at high $Ca^{2+}$), and *CT* (= expression of the CBP relative to the TP) on the $Ca^{2+}$-dependence of the curves $TP^*_{ss,iii.4}$ and $\tau_{iii.4}$ for fixed *K* and *n*.

Module iii.5

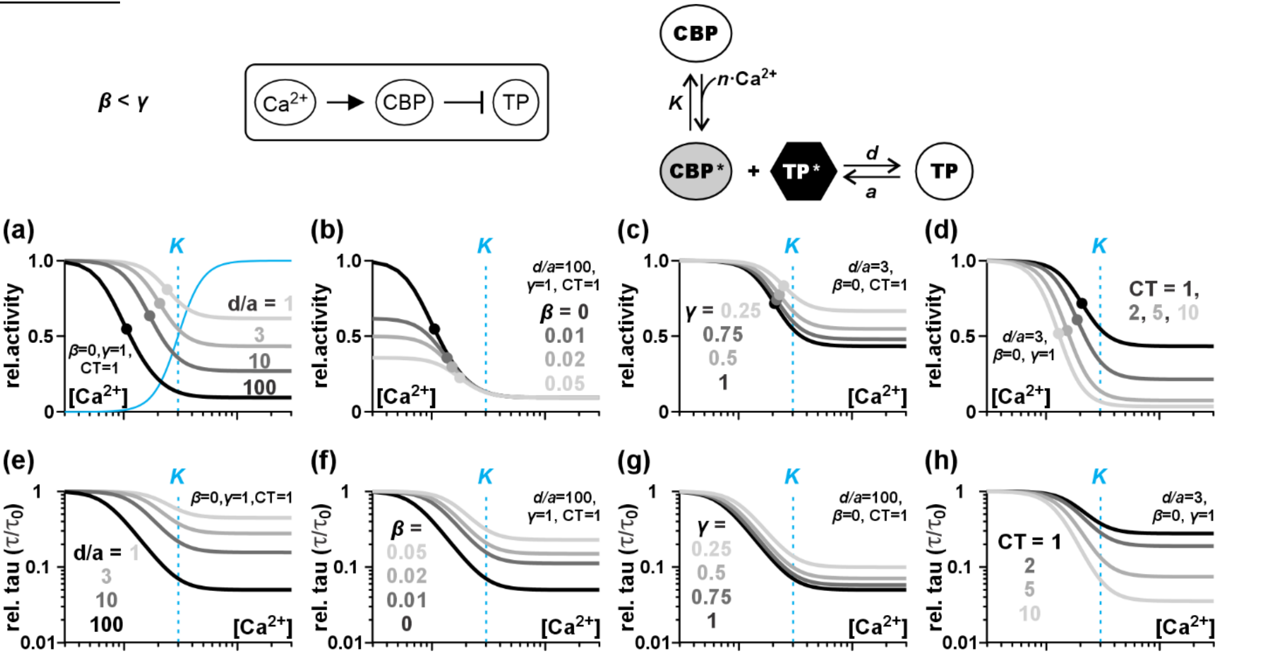


**Fig. S12** Effect of the parameters *d/a*, *β*, *γ*, and *CT* on the Ca$^{2+}$-dependence of module iii.5. The parameters *K*=3 and *n*=4 were fixed. (a-d) Ca$^{2+}$-dependent steady state activity of the target protein in module iii.5 illustrated for different parameters (a) *d/a* (=affinity of the CBP to the TP), (b) *β* (=activity of the CBP at low Ca$^{2+}$), (c) *γ* (=activity of the CBP at high Ca$^{2+}$), and (d) *CT* (= expression of the CBP relative to the TP, $CT = CBP_{total}/TP_{total}$). The solid blue curve shows the Ca$^{2+}$-dependence of the CBP for *β*=0/*γ*=1 and the dashed blue lines indicate the midpoint (*K*) of the CBP-curves. The midpoints of the $TP_{ss}$ curves are indicated as dots and grey-scale coded with the curves and the indicated values for the parameters. Please note that with increasing affinity or an increasing CBP expression the midpoint of the $TP_{ss}$ curve shifts to lower Ca$^{2+}$-values, *i.e.* the sensitivity of the TP to Ca$^{2+}$ increases. Panel (b) shows the curves at different *β* for fixed values *d/a*=100/*γ*=1/*CT*=1, panel (c) for different *γ* [*d/a*=3/*β*=0/*CT*=1], while panel (d) illustrates the dependency on CT [*d/a*=3/*β*=0/*γ*=1]. (e-h) Ca$^{2+}$-dependence of the time constant $\tau$ displayed relative to the $\tau$-value at zero Ca$^{2+}$, $\tau_0$. Please note the logarithmic scale of the y-axes. With increasing affinity of the CBP to the TP (panel e, parameter *d/a*), or a higher CBP expression (panel h, parameter *CT*) the time constant of the module becomes smaller, *i.e.* the module reacts faster at higher Ca$^{2+}$. (f,g) An increase in the basal activity of the CBP at low Ca$^{2+}$ (parameter *β*) or a decrease in activity of the CBP at high Ca$^{2+}$ (parameter *γ*) reduce the difference between the time constants at high and at low Ca$^{2+}$.

$$TP^*_{ss,iii.5} = \frac{1}{2}\cdot\left[\frac{(\beta\cdot d/a-1)\cdot K^n+(\gamma\cdot d/a-1)\cdot[\mathrm{Ca}^{2+}]^n}{\left(\beta\cdot d/a\cdot K^n+\gamma\cdot d/a\cdot[\mathrm{Ca}^{2+}]^n\right)} - CT + \sqrt{\left(CT+\frac{(1+\beta\cdot d/a)\cdot K^n+(1+\gamma\cdot d/a)\cdot[\mathrm{Ca}^{2+}]^n}{\beta\cdot d/a\cdot K^n+\gamma\cdot d/a\cdot[\mathrm{Ca}^{2+}]^n}\right)^2 - 4\cdot CT}\right] \qquad \text{Eqn S35}$$

$$\tau_{iii.5} = \frac{2\cdot\ln 2}{d}\cdot\frac{K^n+[\mathrm{Ca}^{2+}]^n}{\beta\cdot K^n+\gamma\cdot[\mathrm{Ca}^{2+}]^n}\cdot\left(\left(CT+\frac{(1+\beta\cdot d/a)\cdot K^n+(1+\gamma\cdot d/a)\cdot[\mathrm{Ca}^{2+}]^n}{\beta\cdot d/a\cdot K^n+\gamma\cdot d/a\cdot[\mathrm{Ca}^{2+}]^n}\right)^2 - 4\cdot CT\right)^{-\frac{1}{2}} \qquad \text{Eqn S36}$$

$TP^*_{ss,iii.5}$ decreases with increasing [Ca$^{2+}$]. The midpoint calcium concentration of the curve is smaller than the midpoint Ca$^{2+}$-concentration of the calcium-binding protein (CBP), $[\mathrm{Ca}^{2+}]_{mid,TP5} \leq [\mathrm{Ca}^{2+}]_{mid,CBP}$. Thus, the TP is more sensitive to Ca$^{2+}$ than the CBP. The time constant of module iii.5, $\tau_{iii.5}$, decreases with increasing [Ca$^{2+}$]. Both, midpoint and slope of the curves can be different from midpoint and slope of the respective $TP^*_{ss,iii.5}$ curves. **Fig. S12** illustrates the influence of the parameters $d/a$ (affinity of the CBP to the TP), $\beta$ (=activity of the CBP at low Ca$^{2+}$), $\gamma$ (=activity of the CBP at high Ca$^{2+}$), and *CT* (= expression of the CBP relative to the TP) on the Ca$^{2+}$-dependence of the curves $TP^*_{ss,iii.5}$ and $\tau_{iii.5}$ for fixed *K* and *n*.

Module iii.6

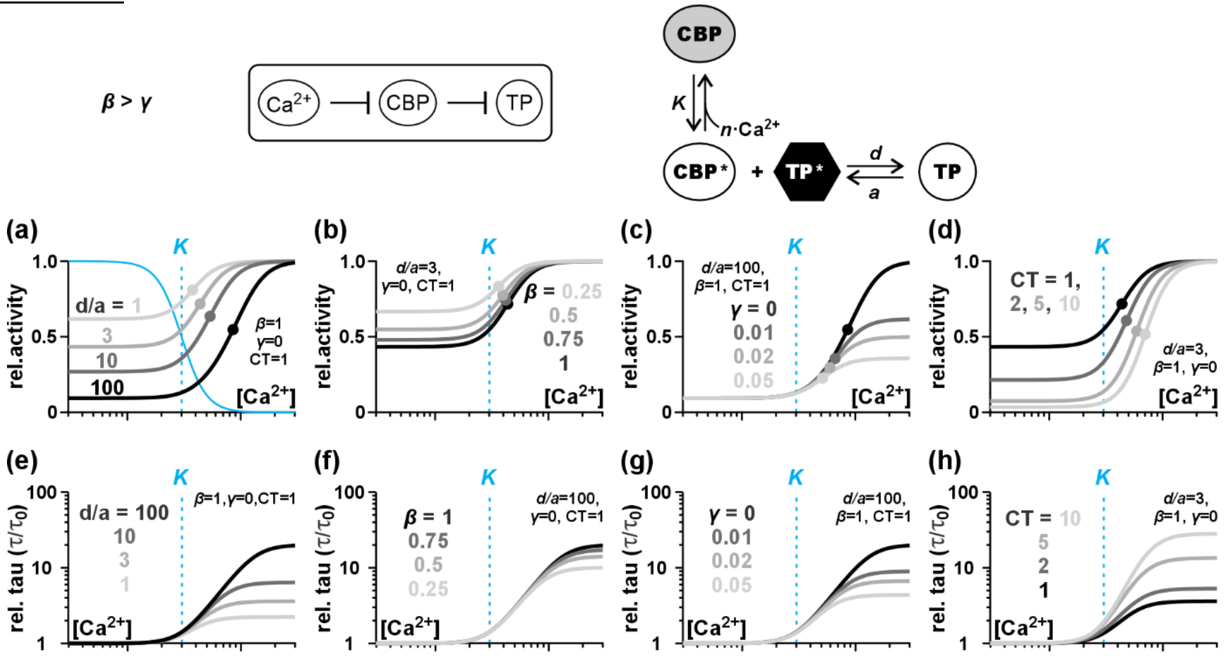


**Fig. S13** Effect of the parameters *d/a*, *β*, *γ*, and *CT* on the $Ca^{2+}$-dependence of module iii.6. The parameters *K*=3 and *n*=4 were fixed. (a-d) $Ca^{2+}$-dependent steady state activity of the target protein in module iii.6 illustrated for different parameters (a) *d/a* (=affinity of the CBP to the TP), (b) *β* (=activity of the CBP at low $Ca^{2+}$), (c) *γ* (=activity of the CBP at high $Ca^{2+}$), and (d) *CT* (= expression of the CBP relative to the TP, $CT = CBP_{total}/TP_{total}$). The solid blue curve shows the $Ca^{2+}$-dependence of the CBP for *β*=1/*γ*=0 and the dashed blue lines indicate the midpoint (*K*) of the CBP-curves. The midpoints of the $TP_{ss}$ curves are indicated as dots and grey-scale coded with the curves and the indicated values for the parameters. Please note that with increasing affinity or an increasing CBP expression the midpoint of the $TP_{ss}$ curve shifts to higher $Ca^{2+}$-values, *i.e.* the sensitivity of the TP to $Ca^{2+}$ decreases. Panel (b) shows the curves at different *β* for fixed values *d/a*=3/*γ*=0/*CT*=1, panel (c) for different *γ* [*d/a*=100/*β*=1/*CT*=1], while panel (d) illustrates the dependency on CT [*d/a*=3/*β*=1/*γ*=0]. (e-h) $Ca^{2+}$-dependence of the time constant $\tau$ displayed relative to the $\tau$-value at zero $Ca^{2+}$, $\tau_0$. Please note the logarithmic scale of the y-axes. With increasing affinity of the CBP to the TP (panel e, parameter *d/a*) or a higher CBP expression (panel h, parameter *CT*) the time constant of the module increases, *i.e.* the module reacts more slowly at higher $Ca^{2+}$. (f,g) An increase in the basal activity of the CBP at low $Ca^{2+}$ (parameter *β*) or a decrease in activity of the CBP at high $Ca^{2+}$ (parameter *γ*) reduce the difference between the time constants at high and at low $Ca^{2+}$.

$$TP^*_{ss,iii.6} = \frac{1}{2}\cdot\left[\frac{(\beta\cdot d/a-1)\cdot K^n+(\gamma\cdot d/a-1)\cdot\left[\mathrm{Ca}^{2+}\right]^n}{\left(\beta\cdot d/a\cdot K^n+\gamma\cdot d/a\cdot\left[\mathrm{Ca}^{2+}\right]^n\right)} - CT + \sqrt{\left(CT+\frac{(1+\beta\cdot d/a)\cdot K^n+(1+\gamma\cdot d/a)\cdot\left[\mathrm{Ca}^{2+}\right]^n}{\beta\cdot d/a\cdot K^n+\gamma\cdot d/a\cdot\left[\mathrm{Ca}^{2+}\right]^n}\right)^2 - 4\cdot CT}\right] \quad \text{Eqn S37}$$

$$\tau_{iii.6} = \frac{2\cdot\ln 2}{d}\cdot\frac{K^n+\left[\mathrm{Ca}^{2+}\right]^n}{\beta\cdot K^n+\gamma\cdot\left[\mathrm{Ca}^{2+}\right]^n}\cdot\left(\left(CT+\frac{(1+\beta\cdot d/a)\cdot K^n+(1+\gamma\cdot d/a)\cdot\left[\mathrm{Ca}^{2+}\right]^n}{\beta\cdot d/a\cdot K^n+\gamma\cdot d/a\cdot\left[\mathrm{Ca}^{2+}\right]^n}\right)^2 - 4\cdot CT\right)^{-\frac{1}{2}} \quad \text{Eqn S38}$$

$TP^*_{ss,iii.6}$ increases with increasing [$Ca^{2+}$]. The midpoint calcium concentration of the curve is larger than the midpoint $Ca^{2+}$-concentration of the calcium-binding protein (CBP), $\left[\mathrm{Ca}^{2+}\right]_{mid,TP6} \geq \left[\mathrm{Ca}^{2+}\right]_{mid,CBP}$. Thus, the TP is less sensitive to $Ca^{2+}$ than the CBP. The time constant of module iii.6, $\tau_{iii.6}$, increases with increasing [$Ca^{2+}$]. Both, midpoint and slope of the curves can be different from midpoint and slope of the respective $TP^*_{ss,iii.6}$ curves. **Fig. S13** illustrates the influence of the parameters $d/a$ (affinity of the CBP to the TP), $\beta$ (=activity of the CBP at low $Ca^{2+}$), $\gamma$ (=activity of the CBP at high $Ca^{2+}$), and *CT* (= expression of the CBP relative to the TP) on the $Ca^{2+}$-dependence of the curves $TP^*_{ss,iii.6}$ and $\tau_{iii.6}$ for fixed *K* and *n*.

Although the equations for the modules iii.3-iii.6 appear to be fundamentally different from those of the modules i.3-i.6 or ii.3-ii.6, the curves are highly similar exhibiting the same tendencies. This similarity is no coincidence, but has two (mathematical) reasons:

1) The hyperbolic tangent can be well approximated by $\tanh(\ln 2 \cdot x) \approx (1 - e^{-x})$ (**Fig. S14**). This allows the equation (S30) to be written as equation (S7): $TP^*_{iii}(t) \approx TP^*_{ss,iii} + \left(TP^*_{0,iii} - TP^*_{ss,iii}\right) \cdot e^{-t/\tau_{iii}}$.

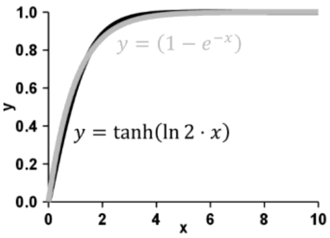


**Fig. S14** Equivalence between $\tanh(\ln 2 \cdot x)$ (*black curve*) and $(1 - e^{-x})$ (*grey curve*). Considering the noise in cellular reactions, the difference between both curves is negligible in physiological processes.

2) The function $\sqrt{(x+A)^2 - 4 \cdot x}$ can be approximated for $x > 0$ and $A > 1$ by (first-order Taylor approximation, **Fig. S15**)

$$\sqrt{(x+A)^2 - 4 \cdot x} = (x+A) \cdot \sqrt{1 - \frac{4 \cdot x}{(x+A)^2}} \approx (x+A) \cdot \left(1 - \frac{2 \cdot x}{(x+A)^2}\right) = x + A - \frac{2 \cdot x}{x+A}$$

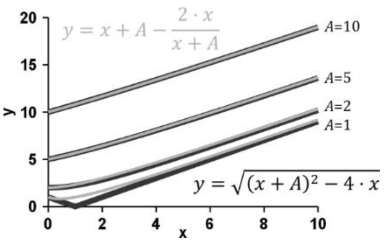


**Fig. S15** Equivalence between $\sqrt{(x+A)^2 - 4 \cdot x}$ (*black curves*) and $x + A - \frac{2 \cdot x}{x+A}$ (*grey curves*).

With these approximations, the steady state values can be written in standard form [eqn. (S1)] as follows:

$$TP^*_{ss,iii.x} \approx \frac{CT}{CT + \frac{a \cdot f + d}{a \cdot f}} = \frac{CT \cdot f \cdot a}{d + (1+CT) \cdot f \cdot a} = \frac{\frac{CT \cdot \beta \cdot a/d}{1+(1+CT) \cdot \beta \cdot a/d} \cdot \left(\frac{1+(1+CT) \cdot \beta \cdot a/d}{1+(1+CT) \cdot \gamma \cdot a/d}\right) \cdot K^n + \frac{CT \cdot \gamma \cdot a/d}{1+(1+CT) \cdot \gamma \cdot a/d} \cdot \left[\mathrm{Ca}^{2+}\right]^n}{\left(\frac{1+(1+CT) \cdot \beta \cdot a/d}{1+(1+CT) \cdot \gamma \cdot a/d}\right) \cdot K^n + \left[\mathrm{Ca}^{2+}\right]^n} \qquad \text{Eqn S39}$$

$$TP^*_{ss,iii.y} \approx 1 - \frac{CT}{CT + \frac{a + d \cdot f}{d \cdot f}} = \frac{a + d \cdot f}{CT \cdot d \cdot f + a + d \cdot f} = \frac{\frac{1+\beta \cdot d/a}{1+(1+CT) \cdot \beta \cdot d/a} \cdot \left(\frac{1+(1+CT) \cdot \beta \cdot d/a}{1+(1+CT) \cdot \gamma \cdot d/a}\right) \cdot K^n + \frac{1+\gamma \cdot d/a}{1+(1+CT) \cdot \gamma \cdot d/a} \cdot \left[\mathrm{Ca}^{2+}\right]^n}{\left(\frac{1+(1+CT) \cdot \beta \cdot d/a}{1+(1+CT) \cdot \gamma \cdot d/a}\right) \cdot K^n + \left[\mathrm{Ca}^{2+}\right]^n} \qquad \text{Eqn S40}$$

where *x* = 3 or 4, and *y* = 5 or 6. The time constants are then specified as:

$$\tau_{iii.x} = \frac{2 \cdot \ln 2/(a \cdot f)}{\sqrt{\left(CT + \frac{a \cdot f + d}{a \cdot f}\right)^2 - 4 \cdot CT}} \approx \frac{2 \cdot \ln 2}{a \cdot f} \cdot \frac{1}{CT + \frac{a \cdot f + d}{a \cdot f} - \frac{2 \cdot CT}{CT + \frac{a \cdot f + d}{a \cdot f}}} = \frac{2 \cdot \ln 2}{d} \cdot \frac{(1+CT) \cdot a/d \cdot f + 1}{\left((1+CT) \cdot a/d \cdot f + 1\right)^2 - 2 \cdot CT \cdot (a/d \cdot f)^2} \qquad \text{Eqn S41}$$

$$\tau_{iii.y} = \frac{2 \cdot \ln 2/(d \cdot f)}{\sqrt{\left(CT + \frac{d \cdot f + a}{d \cdot f}\right)^2 - 4 \cdot CT}} \approx \frac{2 \cdot \ln 2}{d \cdot f} \cdot \frac{1}{CT + \frac{d \cdot f + a}{d \cdot f} - \frac{2 \cdot CT}{CT + \frac{d \cdot f + a}{d \cdot f}}} = \frac{2 \cdot \ln 2}{a} \cdot \frac{(1+CT) \cdot d/a \cdot f + 1}{\left((1+CT) \cdot d/a \cdot f + 1\right)^2 - 2 \cdot CT \cdot (d/a \cdot f)^2} \qquad \text{Eqn S42}$$

which are monotonous sigmoidal functions of [$Ca^{2+}$] that have values of $\tau_0$ at zero and $\tau_\infty$ at very high calcium. Their midpoint is at $\left[\mathrm{Ca}^{2+}\right]_{mid,\tau} = k_\tau$ and the slope at the midpoint can be expressed as $slope_\tau@k_\tau = n_\tau \cdot (\tau_\infty - \tau_0)/4 \cdot k_\tau$. Such a sigmoidal curve can be approximated by:

$$\tau_{iii} \approx \tau_0 \cdot \left(1 + \Delta\tau \cdot \frac{\left[\mathrm{Ca}^{2+}\right]^{n_\tau}}{k_\tau^{n_\tau} + \left[\mathrm{Ca}^{2+}\right]^{n_\tau}}\right) \qquad \text{Eqn S43}$$

with $\Delta\tau = (\tau_\infty - \tau_0)/\tau_0$.

**Table S3**. Conversion between the parameter sets $(n, K, \beta, \gamma, CT, a, d)$ and $(n, k, b, c, \tau_0, \Delta\tau, n_\tau, k_\tau)$ for the case of binding of the CBP to the TP.

| | | | |
|---|---|---|---|
| $TP^*(t) = TP^*_{ss} + (TP^*_0 - TP^*_{ss}) \cdot e^{-t/\tau}$ | | | |
| $TP^*_{ss} = \frac{b \cdot k^n + c \cdot [\text{Ca}^{2+}]^n}{k^n + [\text{Ca}^{2+}]^n}$ $\quad \tau \approx \tau_0 \cdot \left(1 + \Delta\tau \cdot \frac{[\text{Ca}^{2+}]^{n_\tau}}{k_\tau^{n_\tau} + [\text{Ca}^{2+}]^{n_\tau}}\right)$ | | | |
| $\boldsymbol{\beta < \gamma}$ | **Module 1**<br>$Ca^{2+}$ → TP | **Module iii.3 ***<br>$Ca^{2+}$ → CBP → TP | **Module iii.5 ****<br>$Ca^{2+}$ → CBP ⊣ TP |
| $n$<br>Hill coefficient | $n$ | $n$ | $n$ |
| $k$<br>curve midpoint | $k = K$ | $k = \sqrt[n]{\frac{1 + (1 + CT) \cdot \beta \cdot a/d}{1 + (1 + CT) \cdot \gamma \cdot a/d}} \cdot K$ | $k = \sqrt[n]{\frac{1 + (1 + CT) \cdot \beta \cdot d/a}{1 + (1 + CT) \cdot \gamma \cdot d/a}} \cdot K$ |
| $b$<br>activity at 0 $Ca^{2+}$ | $b = \beta$ | $b = \frac{CT \cdot \beta \cdot a/d}{1 + (1 + CT) \cdot \beta \cdot a/d}$ | $b = \frac{1 + \beta \cdot d/a}{1 + (1 + CT) \cdot \beta \cdot d/a}$ |
| $c$<br>activity at high $Ca^{2+}$ | $c = \gamma$ | $c = \frac{CT \cdot \gamma \cdot a/d}{1 + (1 + CT) \cdot \gamma \cdot a/d}$ | $c = \frac{1 + \gamma \cdot d/a}{1 + (1 + CT) \cdot \gamma \cdot d/a}$ |
| $\tau_0$<br>time constant at 0 $Ca^{2+}$ | --- | $\tau_0 = \frac{2 \cdot \ln 2 \cdot ((CT + 1) \cdot a \cdot \beta + d)}{((CT + 1) \cdot a \cdot \beta + d)^2 - 2 \cdot CT \cdot (a \cdot \beta)^2}$ | $\tau_0 = \frac{2 \cdot \ln 2 \cdot ((CT + 1) \cdot d \cdot \beta + a)}{((CT + 1) \cdot d \cdot \beta + a)^2 - 2 \cdot CT \cdot (d \cdot \beta)^2}$ |
| $\tau_\infty$<br>$\tau$ at high $Ca^{2+}$ | --- | $\tau_\infty = \frac{2 \cdot \ln 2 \cdot ((CT + 1) \cdot a \cdot \gamma + d)}{((CT + 1) \cdot a \cdot \gamma + d)^2 - 2 \cdot CT \cdot (a \cdot \gamma)^2}$ | $\tau_\infty = \frac{2 \cdot \ln 2 \cdot ((CT + 1) \cdot d \cdot \gamma + a)}{((CT + 1) \cdot d \cdot \gamma + a)^2 - 2 \cdot CT \cdot (d \cdot \gamma)^2}$ |
| $\Delta\tau$<br>change of $\tau$ with $Ca^{2+}$ | --- | $\Delta\tau = \frac{\tau_\infty - \tau_0}{\tau_0} < 0$ | $\Delta\tau = \frac{\tau_\infty - \tau_0}{\tau_0} < 0$ |
| $k_\tau, n_\tau$ *** | --- | $k_\tau \neq k, k_\tau \neq K, n_\tau \neq n$ | $k_\tau \neq k, k_\tau \neq K, n_\tau \neq n$ |
| $\boldsymbol{\beta > \gamma}$ | **Module 2**<br>$Ca^{2+}$ ⊣ TP | **Module iii.4 ***<br>$Ca^{2+}$ ⊣ CBP → TP | **Module iii.6 ****<br>$Ca^{2+}$ ⊣ CBP ⊣ TP |
| $n$<br>Hill coefficient | $n$ | $n$ | $n$ |
| $k$<br>curve midpoint | $k = K$ | $k = \sqrt[n]{\frac{1 + (1 + CT) \cdot \beta \cdot a/d}{1 + (1 + CT) \cdot \gamma \cdot a/d}} \cdot K$ | $k = \sqrt[n]{\frac{1 + (1 + CT) \cdot \beta \cdot d/a}{1 + (1 + CT) \cdot \gamma \cdot d/a}} \cdot K$ |
| $b$<br>activity at 0 $Ca^{2+}$ | $b = \beta$ | $b = \frac{CT \cdot \beta \cdot a/d}{1 + (1 + CT) \cdot \beta \cdot a/d}$ | $b = \frac{1 + \beta \cdot d/a}{1 + (1 + CT) \cdot \beta \cdot d/a}$ |
| $c$<br>activity at high $Ca^{2+}$ | $c = \gamma$ | $c = \frac{CT \cdot \gamma \cdot a/d}{1 + (1 + CT) \cdot \gamma \cdot a/d}$ | $c = \frac{1 + \gamma \cdot d/a}{1 + (1 + CT) \cdot \gamma \cdot d/a}$ |
| $\tau_0$<br>time constant a 0 $Ca^{2+}$ | --- | $\tau_0 = \frac{2 \cdot \ln 2 \cdot ((CT + 1) \cdot a \cdot \beta + d)}{((CT + 1) \cdot a \cdot \beta + d)^2 - 2 \cdot CT \cdot (a \cdot \beta)^2}$ | $\tau_0 = \frac{2 \cdot \ln 2 \cdot ((CT + 1) \cdot d \cdot \beta + a)}{((CT + 1) \cdot d \cdot \beta + a)^2 - 2 \cdot CT \cdot (d \cdot \beta)^2}$ |
| $\tau_\infty$<br>$\tau$ at high $Ca^{2+}$ | --- | $\tau_\infty = \frac{2 \cdot \ln 2 \cdot ((CT + 1) \cdot a \cdot \gamma + d)}{((CT + 1) \cdot a \cdot \gamma + d)^2 - 2 \cdot CT \cdot (a \cdot \gamma)^2}$ | $\tau_\infty = \frac{2 \cdot \ln 2 \cdot ((CT + 1) \cdot d \cdot \gamma + a)}{((CT + 1) \cdot d \cdot \gamma + a)^2 - 2 \cdot CT \cdot (d \cdot \gamma)^2}$ |
| $\Delta\tau$<br>change of $\tau$ with $Ca^{2+}$ | --- | $\Delta\tau = \frac{\tau_\infty - \tau_0}{\tau_0} > 0$ | $\Delta\tau = \frac{\tau_\infty - \tau_0}{\tau_0} > 0$ |
| $k_\tau, n_\tau$ *** | --- | $k_\tau \neq k, k_\tau \neq K, n_\tau \neq n$ | $k_\tau \neq k, k_\tau \neq K, n_\tau \neq n$ |

* Please note: The parameter *a* contains implicitly the factor $TP_{total}$, *i.e.* the expression level of the TP.
** Please note: The parameter *d* contains implicitly the factor $TP_{total}$, *i.e.* the expression level of the TP.
*** The complex relationships are not further specified here.

#### **Notes S2** Sigmoidal curves and the universal empiric Hill equation

The fraction of $Ca^{2+}$-bound EF hands that follow the chemical reaction $Ca^{2+} + EF \rightleftharpoons EF^C$ is in steady state:

$$EF^C = \frac{[Ca^{2+}]}{K + [Ca^{2+}]} \qquad \text{Eqn S44}$$

The $Ca^{2+}$ binding then may induce a rather rapid conformational change in the EF hand $EF^C \underset{\leftarrow d -}{\overset{-a \rightarrow}{}} EF^*$ that activates the EF hand, with the forward and reverse rate constants *a* and *d*, respectively. In steady state the fraction of active EF hands (EF*) is given by (compare Notes S1, module i.3):

$$EF^* = \frac{\frac{a}{a+d} \cdot [Ca^{2+}]}{\frac{d}{a+d} \cdot K + [Ca^{2+}]} = \frac{\gamma \cdot [Ca^{2+}]}{k + [Ca^{2+}]} \qquad \text{Eqn S45}$$

Both, eqns (S44) and (S45), describe sigmoidal curves. However, they differ in their midpoints and in the saturation level at high $Ca^{2+}$ ($\gamma = \frac{a}{a+d}$, $k = K \cdot \frac{d}{a+d}$; **Figure S16**). Thus, the dissociation constant *K* of the $Ca^{2+}$-binding is correlated to but not identical with the mechanistically relevant $Ca^{2+}$-sensitivity constant *k* of the EF hand. In general, the parameters *a* and *d* are difficult to determine experimentally/computationally, so that a prediction of the *k*-value from the *K*-value or *vice versa* is hardly possible.

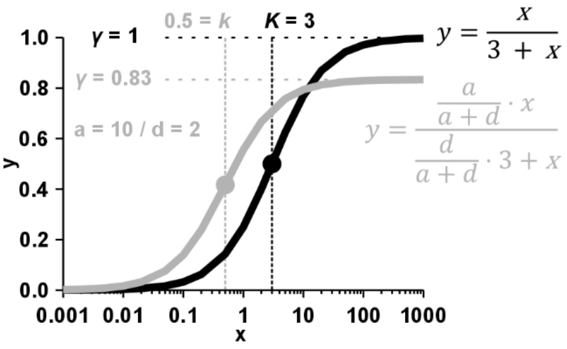


**Fig. S16** Illustration of equations S44 and S45. In this example *K* = 3, *a* = 10, *d* = 2. These values result in *k* = 0.5 and *γ* = 0.83.

Usually, two or four EF hands collaborate to generate a common outcome. If two identical EF hands act independently together to induce a subsequent reaction, the fraction of active EF hand pairs is given by:

$$EFpair^* = \left(\frac{\gamma \cdot [Ca^{2+}]}{k + [Ca^{2+}]}\right)^2 \qquad \text{Eqn S46}$$

This equation describes a sigmoidal curve with EFpair* = 0 in the absence of $Ca^{2+}$, EFpair* = $\gamma^2$ at high $Ca^{2+}$, and midpoint at $\kappa = (\sqrt{2} + 1) \cdot k$. This curve is hardly distinguishable from an empiric Hill function that has the same values at low and high $Ca^{2+}$, the same midpoint and the same slope at the midpoint [Hill-coefficient $n = 2 \cdot (2 - \sqrt{2}) \approx 1.17$] (**Figure S17**):

$$EFpair^* = \left(\frac{\gamma \cdot [Ca^{2+}]}{k + [Ca^{2+}]}\right)^2 \approx \frac{\gamma^2 \cdot [Ca^{2+}]^{2\cdot(2-\sqrt{2})}}{\left((\sqrt{2}+1) \cdot k\right)^{2\cdot(2-\sqrt{2})} + [Ca^{2+}]^{2\cdot(2-\sqrt{2})}} = \frac{c \cdot [Ca^{2+}]^n}{\kappa^n + [Ca^{2+}]^n} \qquad \text{Eqn S47}$$

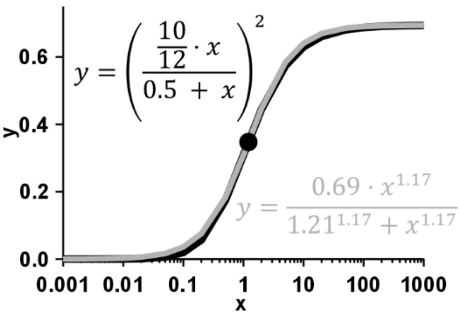


**Fig. S17** Illustration of equations S46 and S47. In this example *k* = 0.5, *γ* = 0.83, *κ* = 1.21, *c* = 0.69, *n* = 1.17. The empiric Hill function (grey) is almost indistinguishable from the "exact" function (black). It is not possible to judge whether equation S46 or S47 is better suited to describe data points that line up along this sigmoidal curve.

### Figure S18

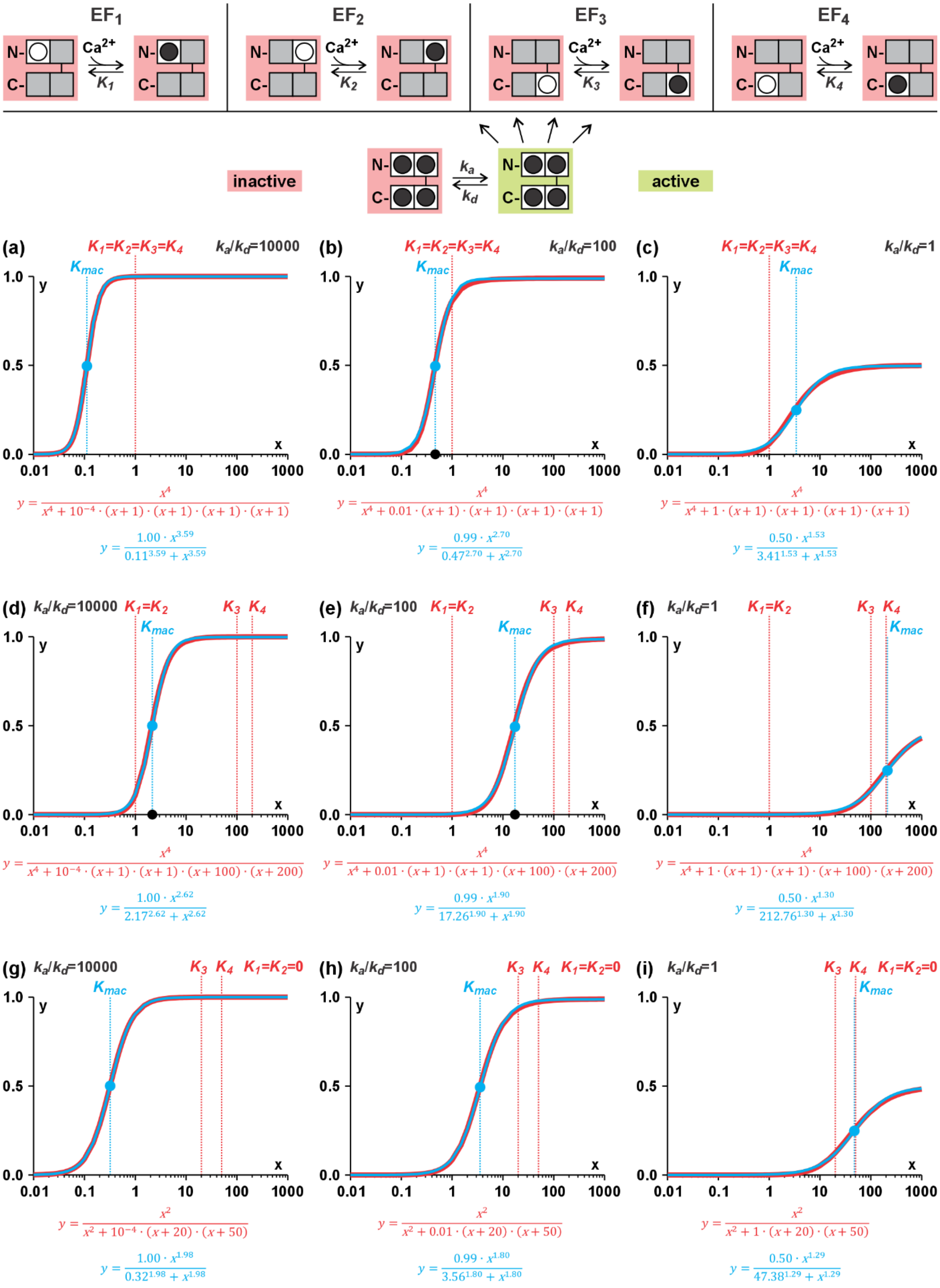
EF$_1$
EF$_2$
EF$_3$
EF$_4$
Ca$^{2+}$
$K_1$
$K_2$
$K_3$
$K_4$
inactive
active
$k_a$
$k_d$
(a) $K_1$=$K_2$=$K_3$=$K_4$ $k_a/k_d$=10000
$K_{mac}$
$y = \frac{x^4}{x^4 + 10^{-4}\cdot(x+1)\cdot(x+1)\cdot(x+1)\cdot(x+1)}$
$y = \frac{1.00\cdot x^{3.59}}{0.11^{3.59} + x^{3.59}}$
(b) $K_1$=$K_2$=$K_3$=$K_4$ $k_a/k_d$=100
$K_{mac}$
$y = \frac{x^4}{x^4 + 0.01\cdot(x+1)\cdot(x+1)\cdot(x+1)\cdot(x+1)}$
$y = \frac{0.99\cdot x^{2.70}}{0.47^{2.70} + x^{2.70}}$
(c) $K_1$=$K_2$=$K_3$=$K_4$ $k_a/k_d$=1
$K_{mac}$
$y = \frac{x^4}{x^4 + 1\cdot(x+1)\cdot(x+1)\cdot(x+1)\cdot(x+1)}$
$y = \frac{0.50\cdot x^{1.53}}{3.41^{1.53} + x^{1.53}}$
(d) $k_a/k_d$=10000 $K_1$=$K_2$ $K_3$ $K_4$
$K_{mac}$
$y = \frac{x^4}{x^4 + 10^{-4}\cdot(x+1)\cdot(x+1)\cdot(x+100)\cdot(x+200)}$
$y = \frac{1.00\cdot x^{2.62}}{2.17^{2.62} + x^{2.62}}$
(e) $k_a/k_d$=100 $K_1$=$K_2$ $K_3$ $K_4$
$K_{mac}$
$y = \frac{x^4}{x^4 + 0.01\cdot(x+1)\cdot(x+1)\cdot(x+100)\cdot(x+200)}$
$y = \frac{0.99\cdot x^{1.90}}{17.26^{1.90} + x^{1.90}}$
(f) $k_a/k_d$=1 $K_1$=$K_2$ $K_3$ $K_4$
$K_{mac}$
$y = \frac{x^4}{x^4 + 1\cdot(x+1)\cdot(x+1)\cdot(x+100)\cdot(x+200)}$
$y = \frac{0.50\cdot x^{1.30}}{212.76^{1.30} + x^{1.30}}$
(g) $k_a/k_d$=10000 $K_3$ $K_4$ $K_1$=$K_2$=0
$K_{mac}$
$y = \frac{x^2}{x^2 + 10^{-4}\cdot(x+20)\cdot(x+50)}$
$y = \frac{1.00\cdot x^{1.98}}{0.32^{1.98} + x^{1.98}}$
(h) $k_a/k_d$=100 $K_3$ $K_4$ $K_1$=$K_2$=0
$K_{mac}$
$y = \frac{x^2}{x^2 + 0.01\cdot(x+20)\cdot(x+50)}$
$y = \frac{0.99\cdot x^{1.80}}{3.56^{1.80} + x^{1.80}}$
(i) $k_a/k_d$=1 $K_3$ $K_4$ $K_1$=$K_2$=0
$K_{mac}$
$y = \frac{x^2}{x^2 + 1\cdot(x+20)\cdot(x+50)}$
$y = \frac{0.50\cdot x^{1.29}}{47.38^{1.29} + x^{1.29}}$

**Fig. S18** Mathematical description of $Ca^{2+}$ sensitivity with a minimal set of parameters. A protein has four EF hands that are arranged as pairs in an N- and a C-lobe. Individual $Ca^{2+}$ binding to each EF hand is characterized by the K-values $K_1$, $K_2$, $K_3$, and $K_4$. In a final step, the protein with $Ca^{2+}$-saturated EF hands can activate (*green*) with the rates $k_a$ (forward) and $k_d$ (backward). (a-i) Examples of the relative activity of the protein in steady state (y-axis) as a function of the $Ca^{2+}$ concentration (x-axis). The red curves (*mathematical descriptions in red*) were calculated according to the outlined model with the specified parameters $K_1$, $K_2$, $K_3$, $K_4$ (indicated also by the *red vertical dashed lines*), and $k_a/k_d$. The blue curves are fits of the red curves with the equation (1) (*mathematical descriptions in blue*). The vertical blue dashed lines indicate the macroscopic K-values ($K_{mac}$). Please note that the $Ca^{2+}$ sensitivity of the system increases (*n* larger, $K_{mac}$ smaller) with rising $k_a/k_d$ ratios, even though the *K* values of the EF hands remain unchanged.

#### Figure S19

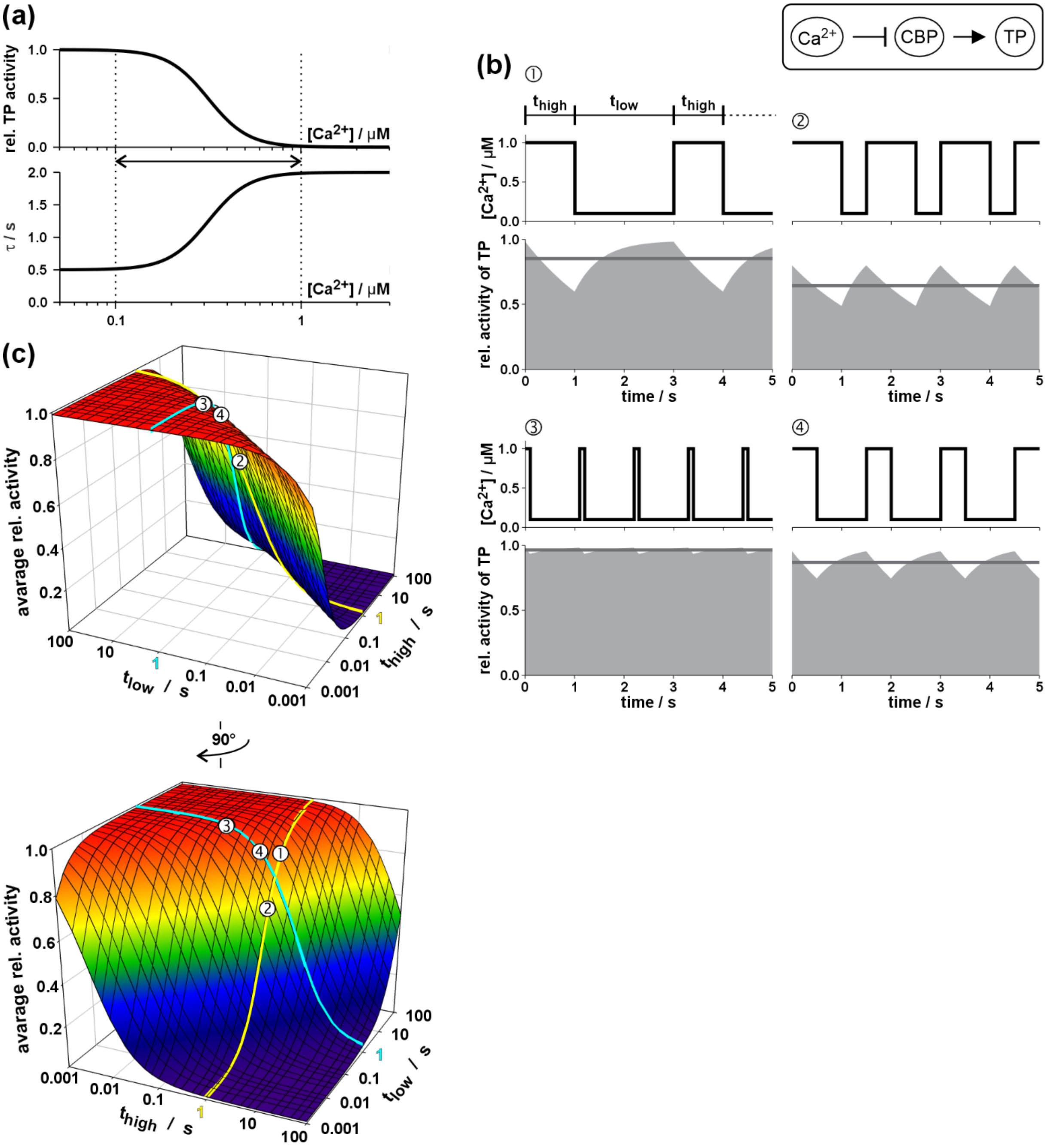


**Fig. S19** Frequency dependence of module 4. (a) $Ca^{2+}$ dependence of the steady state activity of the target protein (TP) and the time constant ($\tau$) of the relaxation processes. The dashed lines indicate the $Ca^{2+}$ values between which oscillations occurred in the various scenarios. (b) Four scenarios with repetitive pulses between high and low $Ca^{2+}$ (①-④) are shown with different durations $t_{high}$ and $t_{low}$. The average activity of TP over time (grey line) is given by the time-average over the grey surface under the curve. (c) 3D-plot of the average relative activity of TP as a function of the two parameters $t_{high}$ and $t_{low}$. The data of the four presented scenarios are indicated and cross-referenced by numbers ①-④. For better orientation the values for $t_{high}$ = 1s and $t_{low}$ = 1s are indicated by a yellow and a cyan line, respectively.

#### Figure S20

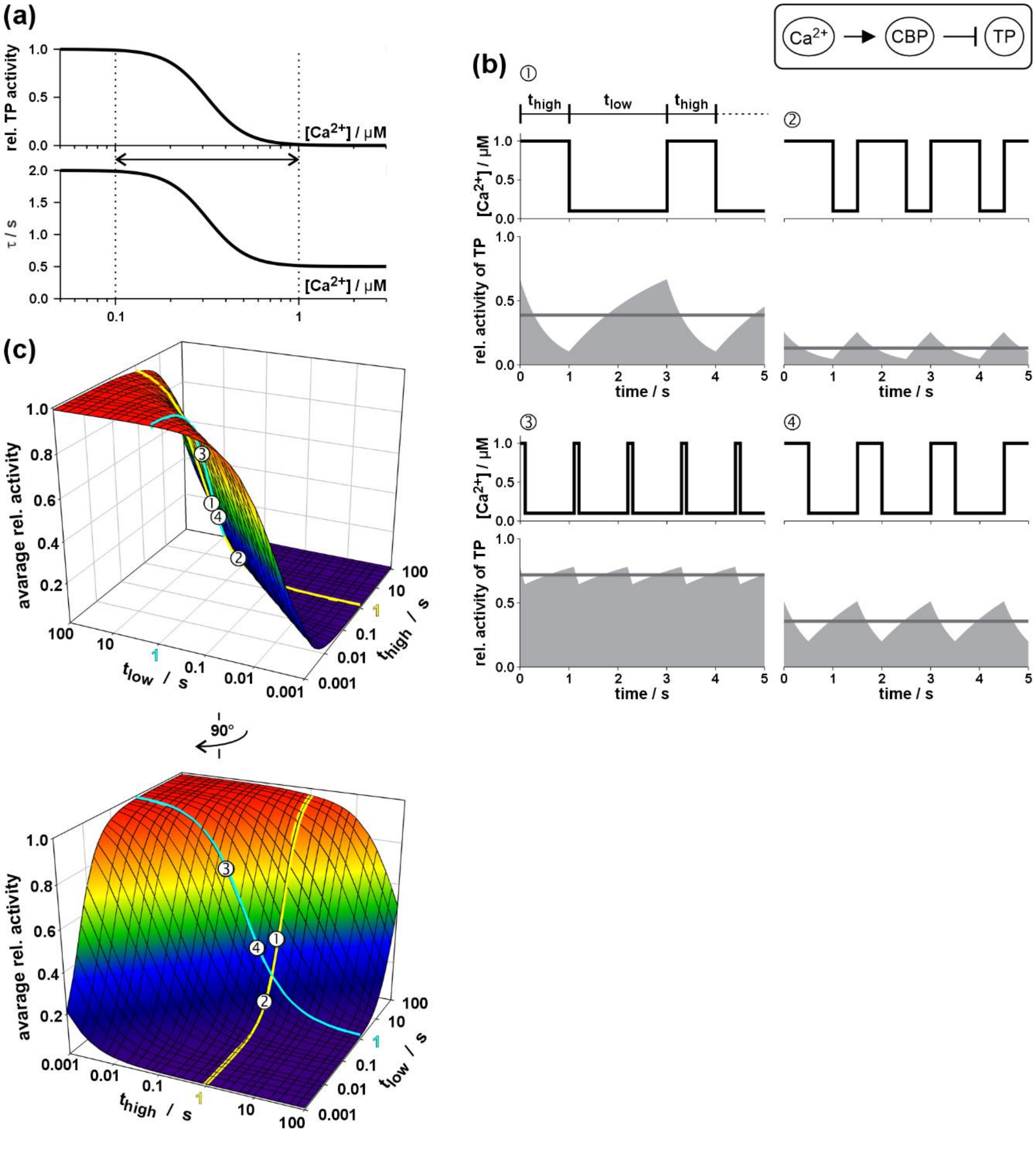


**Fig. S20** Frequency dependence of module 5. (a) $Ca^{2+}$ dependence of the steady state activity of the target protein (TP) and the time constant ($\tau$) of the relaxation processes. The dashed lines indicate the $Ca^{2+}$ values between which oscillations occurred in the various scenarios. (b) Four scenarios with repetitive pulses between high and low $Ca^{2+}$ (①-④) are shown with different durations $t_{high}$ and $t_{low}$. The average activity of TP over time (grey line) is given by the time-average over the grey surface under the curve. (c) 3D-plot of the average relative activity of TP as a function of the two parameters $t_{high}$ and $t_{low}$. The data of the four presented scenarios are indicated and cross-referenced by numbers ①-④. For better orientation the values for $t_{high}$ = 1s and $t_{low}$ = 1s are indicated by a yellow and a cyan line, respectively.

#### Figure S21

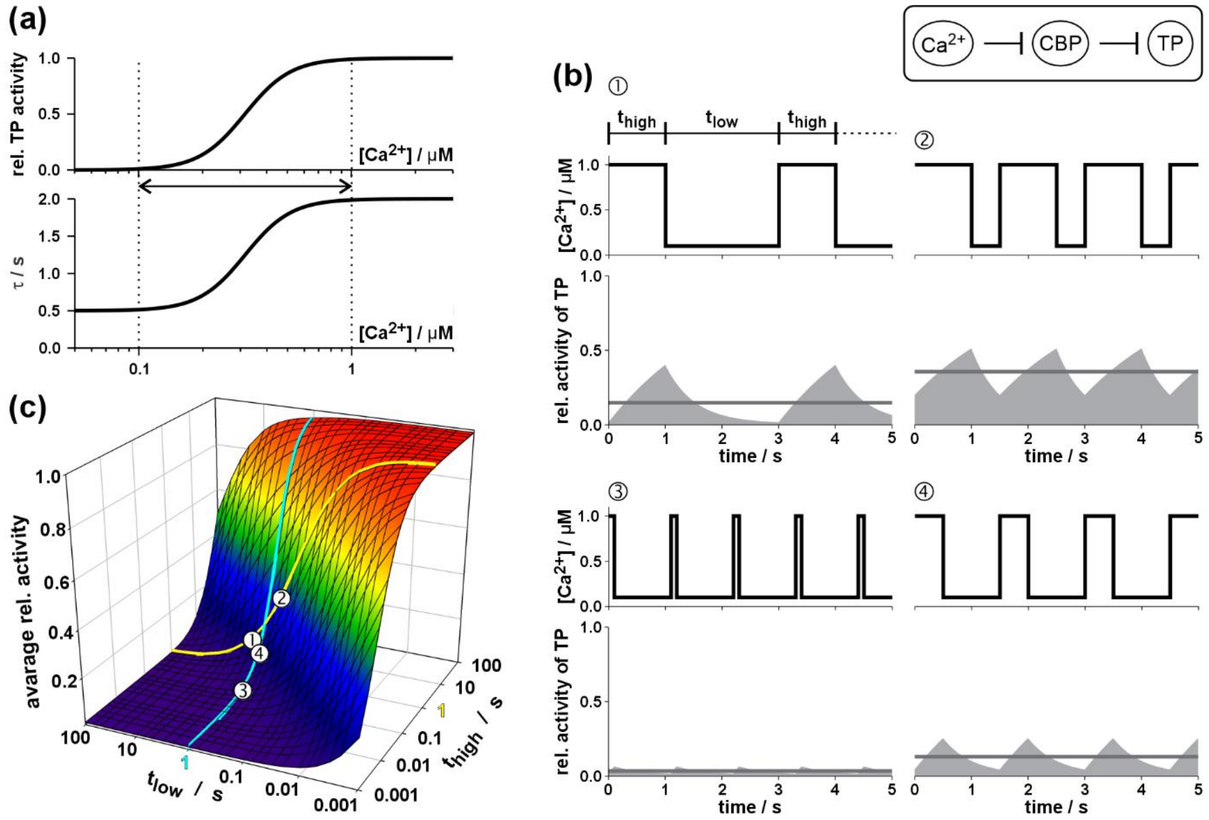


**Fig. S21** Frequency dependence of module 6. (a) $Ca^{2+}$ dependence of the steady state activity of the target protein (TP) and the time constant ($\tau$) of the relaxation processes. The dashed lines indicate the $Ca^{2+}$ values between which oscillations occurred in the various scenarios. (b) Four scenarios with repetitive pulses between high and low $Ca^{2+}$ (①-④) are shown with different durations $t_{high}$ and $t_{low}$. The average activity of TP over time (grey line) is given by the time-average over the grey surface under the curve. (c) 3D-plot of the average relative activity of TP as a function of the two parameters $t_{high}$ and $t_{low}$. The data of the four presented scenarios are indicated and cross-referenced by numbers ①-④. For better orientation the values for $t_{high}$ = 1s and $t_{low}$ = 1s are indicated by a yellow and a cyan line, respectively.

#### Figure S22

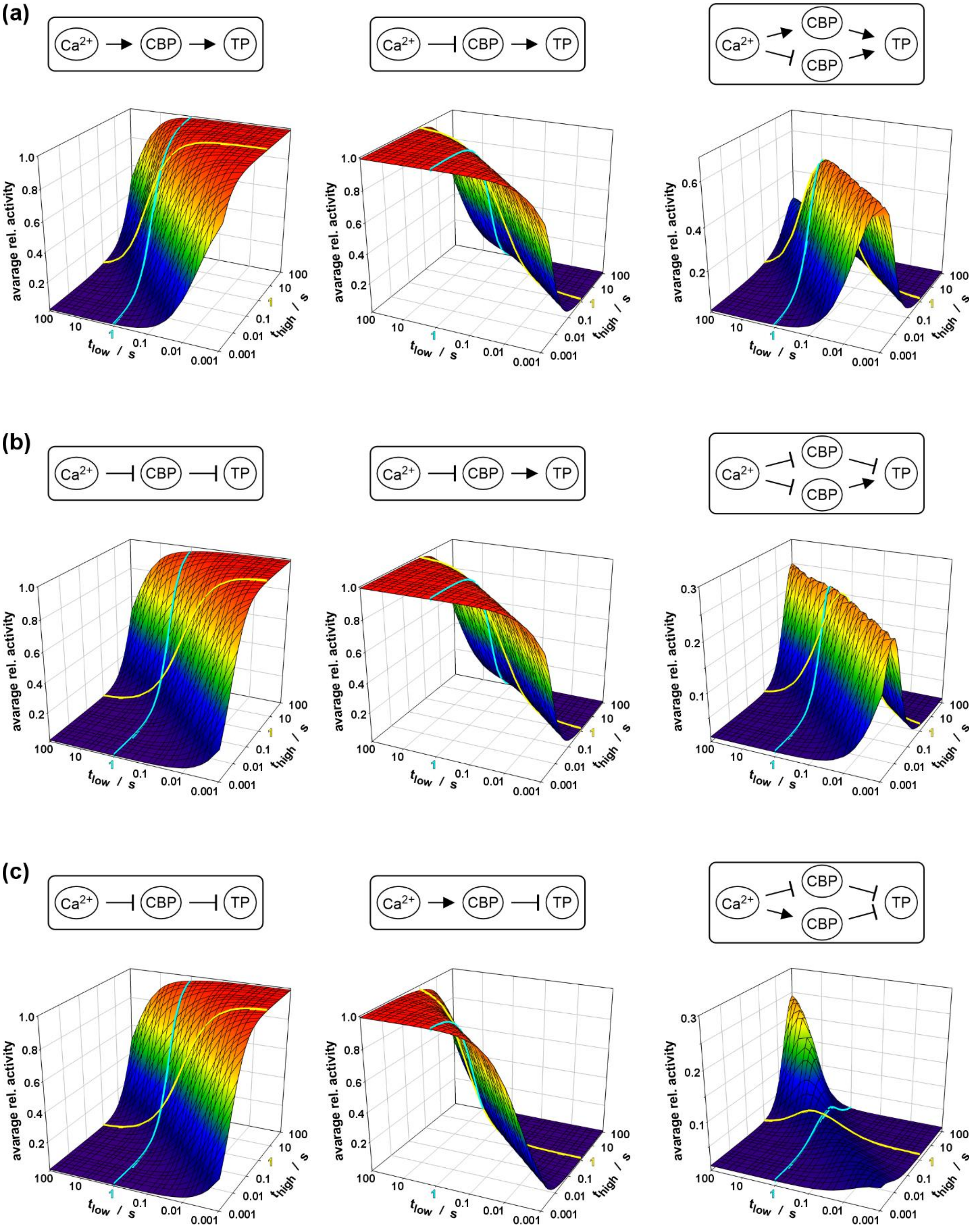


**Fig. S22** Frequency dependence of combined modules. (a) Combination of modules 3 (left) and 4 (middle). (b) Combination of modules 6 (left) and 4 (middle). (c) Combination of modules 6 (left) and 5 (middle).
